# A ‘through-DNA’ mechanism for metal uptake-vs.-efflux regulation

**DOI:** 10.1101/2023.12.05.570191

**Authors:** Udit Kumar Chakraborty, Youngchan Park, Kushal Sengupta, Won Jung, Chandra P. Joshi, Danielle H. Francis, Peng Chen

## Abstract

Transition metals like Zn are essential for all organisms including bacteria, but fluctuations of their concentrations in the cell can be lethal. Organisms have thus evolved complex mechanisms for cellular metal homeostasis. One mechanistic paradigm involves pairs of transcription regulators sensing intracellular metal concentrations to regulate metal uptake and efflux. Here we report that Zur and ZntR, a prototypical pair of regulators for Zn uptake and efflux in *E. coli*, respectively, can coordinate their regulation through DNA, besides sensing cellular Zn^2+^ concentrations. Using a combination of live-cell single-molecule tracking and *in vitro* single-molecule FRET measurements, we show that unmetallated ZntR can enhance the unbinding kinetics of Zur from DNA by directly acting on Zur-DNA complexes, possibly through forming heteromeric ternary and quaternary complexes that involve both protein-DNA and protein-protein interactions. This ‘through-DNA’ mechanism may functionally facilitate the switching in Zn uptake regulation when bacteria encounter changing Zn environments; it could also be relevant for regulating the uptake-vs.-efflux of various metals across different bacterial species and yeast.

## Main Text

For all life forms including bacteria, transition metals like Zn are essential but their excess is also detrimental^1–12^. Host cells can sequester metals to curb bacterial proliferation during infection, while metal stress can also be effective bactericidal treatments^8,10–13^. For growth and survival, bacteria have evolved exquisite mechanisms to regulate metal uptake and efflux^6,8,13–16^. Studying bacteria has thus produced mechanistic paradigms not only for understanding metal homeostasis in general but also for developing antibiotic treatments^7,17,18^.

One such paradigm is the ‘set-point’ mechanism that bacteria use to regulate cellular concentrations of a variety of transition metals (e.g., Zn^2+^, Fe^2+^, Ni^2+^, etc.)^1–6^. Here, the cellular free metal concentration [M^n+^]_free_ is bound by the metal-binding affinities of the respective uptake and efflux regulators. In *E. coli*, the Fur-family metalloregulator Zur^19^ and the MerR-family metalloregulator ZntR^20,21^ are the major Zn uptake and efflux regulators that control the free or bio-available Zn^2+^ concentration in the cell (i.e., [Zn^2+^]_free_) to a range set by their respective Zn^2+^ binding affinities (Fig. 1a). Under Zn deficiency ([Zn^2+^]_free_ < 0.2 fM), Zur has vacant regulatory Zn-binding sites and is a non-repressor that binds to non-consensus DNA sites^22^; here Zn uptake genes (e.g., *znuABC*) are actively transcribed, while ZntR is at its apo state (i.e., ZntR_apo_) and binds its cognate promoter tightly, repressing Zn efflux genes (e.g., *zntA*). Under Zn replete conditions where [Zn^2+^]_free_ exceeds 0.2 fM, Zur becomes fully metallated (i.e., Zur_Zn_) and binds to its cognate promoter sites tightly, repressing Zn uptake. When [Zn^2+^]_free_ rises further above 1.1 fM (i.e., Zn excess), ZntR is metallated (i.e., ZntR_Zn_) to become an activator at its cognate promoters, activating Zn efflux.

**Fig. 1.**
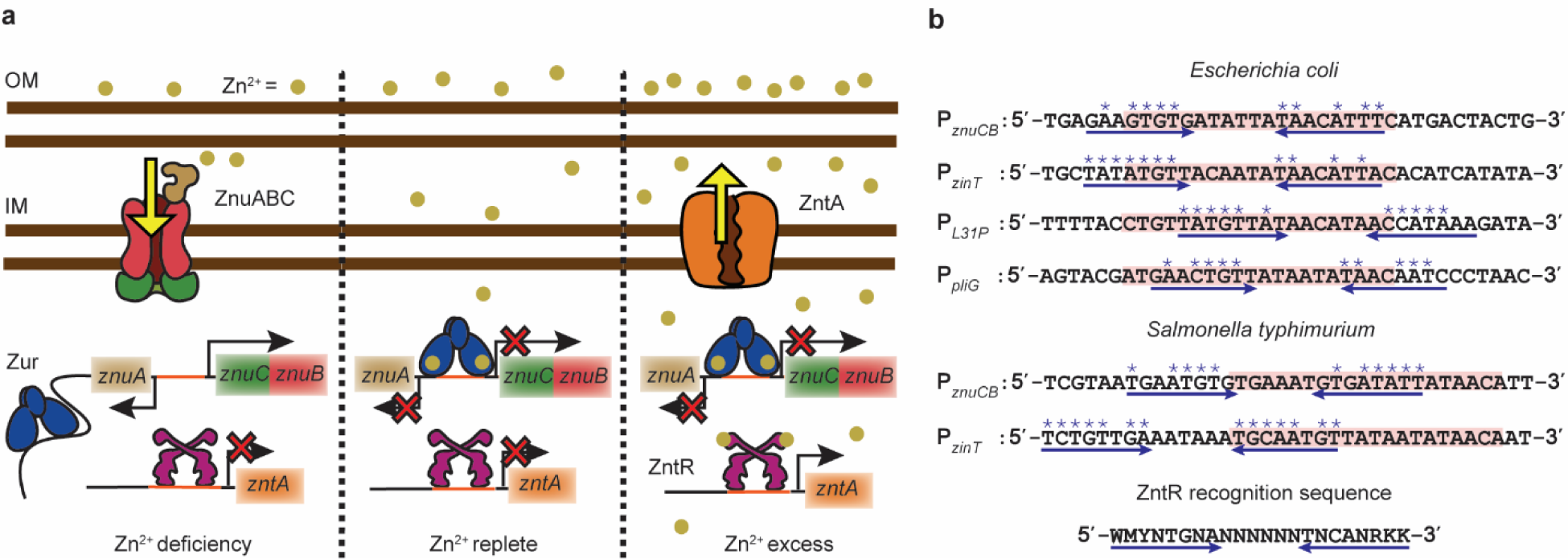
a, Present functional paradigm of Zur and ZntR in bacteria. Left: Under Zn^2+^ deficiency, Zur, without metallation at its regulatory sites, is a non-repressor that binds to non-consensus DNA sites but not to its recognition sequence (i.e., Zur box) in the promoter regions of its regulons (e.g., the divergent *znuABC* operon); here Zn^2+^ uptake genes are derepressed, while the non-metallated ZntR binds to its recognition sequence in the promoter regions of its regulons (e.g., *zntA*), repressing Zn^2+^ efflux. Center: Under Zn^2+^ replete conditions, Zur starts to be fully metallated (Zur_Zn_) and binds to Zur box, repressing Zn^2+^ uptake, while ZntR is still predominantly in its apo form, repressing Zn^2+^ efflux. Right: Under Zn^2+^ excess, fully metallated Zur_Zn_ keeps repressing Zn^2+^ uptake, while the metallated ZntR_Zn_ at its cognate promoters distorts the DNA to result in activation of Zn^2+^ efflux genes. Both Zur and ZntR also have a freely diffusing population in the cell (not shown). IM: inner membrane; OM: outer membrane. **b**, **Promoter region sequences of Zur regulons in two different bacteria**. Pink shades: Zur boxes. Double blue arrows: possible dyad symmetric sequences recognized by ZntR, whose consensus recognition sequence is shown at the bottom. Asterisk (*) denotes matches with the consensus sequence. Analysis of other species in Extended Data Fig. 1.

While the binding of Zur and ZntR to their cognate promoters leads to transcription repression or activation of Zn uptake or efflux genes, their unbinding is key to resetting regulation status when environmental and cellular Zn levels change. We have recently uncovered an unusual, facilitated unbinding mechanism for Zur and ZntR from DNA^22–24^. There, a homotypic freely diffusing protein can either assist the dissociation of the incumbent protein on DNA or directly substitute it, likely through an intermediate ternary complex enabled by the multivalent contact between the protein and DNA, leading to protein-concentration−enhanced unbinding kinetics. Such facilitated unbinding allows for more facile switching between repression and derepression of Zn uptake genes or between activation and deactivation of Zn efflux genes; it was also observed for other types of DNA-binding proteins^24–31^. Additionally, Zur shows an impeded unbinding from its oligomerization-rendered stabilization on DNA^22^, which allows for storing the non-repressor form of Zur at non-consensus chromosomal sites longer to not interfere with its repressor form at cognate promoter sites. Overall, the apparent first-order unbinding rate constant *k*_−1_ of Zur and ZntR from tight binding sites on DNA follows^22,23^:

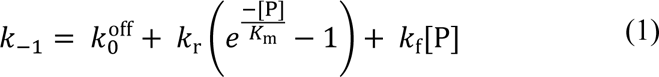

Here 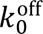 is a first-order intrinsic unbinding rate constant. The second term only applies to Zur and accounts for its impeded unbinding pathway, where *k*_r_ is a first-order rate constant, *K*_m_ is an effective dissociation constant of protein oligomer on DNA, and [P] is the free protein concentration. The third term is for the facilitated unbinding pathway for both Zur and ZntR with *k*_f_ being a second-order rate constant.

As both Zur and ZntR can act on themselves on DNA, we wondered whether they could also act on each other, leading to cross-communication directly on DNA between the two regulatory systems. Here, we report the discovery of partial ZntR recognition sequences that overlap with Zur binding boxes in the promoters of Zur regulons. Using single-molecule tracking and single-cell protein quantitation, we show that in live *E. coli* cells, the unmetallated ZntR_apo_ can enhance the unbinding kinetics of both repressor and non-repressor forms of Zur from DNA, whereas metallated ZntR_Zn_ cannot. We further show, through *in vitro* single-molecule FRET measurements, that ZntR_apo_ directly acts on Zur-DNA complexes, possibly through forming heteromeric ternary and quaternary complexes that involve both protein-DNA and protein-protein interactions; this direct action gives rise to a *first-of-its-kind* ‘through-DNA’ mechanism for their cross-actions in regulating Zn homeostasis. Moreover, this mechanism is likely functionally significant in facilitating the switching in Zn uptake regulation when an *E. coli* cell encounters changing Zn environments; it may even be broadly relevant for regulating uptake-vs.-efflux of Zn and other metals for different bacterial species and yeast.

### Zur cognate promoters contain partial ZntR recognition sequences

We examined the DNA sequences around Zur’s and ZntR’s operator sites in *E. coli*’s genome. Strikingly, at the promoters of the four known Zur regulons [i.e., *znuABC*, *zinT* (a periplasmic Zn chaperone), *l31p* and *l36p* (a pair of ribosomal proteins), and *pliG* (a periplasmic lysozyme inhibitor)]^32–35^, the Zur box overlaps with sequences that match significantly with ZntR’s recognition sequence (Fig. 1b; Supplementary Information 2), suggesting possible direct involvement of ZntR in Zur-DNA interactions. At promoters of known ZntR regulons (e.g., *zntA*), we did not identify clear Zur binding sites.

### ZntR_apo_ enhances unbinding of repressor Zur_Zn_ from DNA in cells

The discovery of partial ZntR recognition sequences around Zur boxes prompted us to examine whether ZntR can affect Zur unbinding from DNA. We first examined how ZntR_apo_ may affect Zur_Zn_ unbinding, as ZntR_apo_ and Zur_Zn_ coexist in the cell under the normal Zn replete conditions (Fig. 1a, center).

To visualize Zur_Zn_ in the cell, we tagged Zur at its C-terminus with the photoconvertible fluorescent protein mEos3.2 (i.e., Zur^mE^), either at its chromosomal locus or additionally on an inducible plasmid to access a broader range of cellular protein concentrations, as previously reported (Methods; Supplementary Information 1.1)^22^. We cultured and imaged cells in the presence of 20 μM Zn^2+^, under which cellular Zur is known to be dominantly in its metallated repressor form 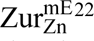. The unbinding kinetics of 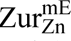 from DNA in the cell was measured using single-molecule tracking (SMT), as we reported (Extended Data Fig. 2a; Supplementary Information 1.2.2)^22^. Briefly, we used controlled photoconversion coupled with time-lapse stroboscopic imaging to track single 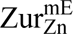 molecules until the fluorescent tag photobleached (Fig. 2a). The displacement length distribution obtained from SMT can resolve its three diffusion states: freely diffusing in the cytosol (FD), non-specifically bound to DNA (NB), and tightly bound to DNA (TB), including their effective diffusion constants and fractional populations (Fig. 2b; Supplementary Information 4). Thresholding the displacement-vs.-time trajectories allowed us to extract Zur’s microscopic residence times *τ* on DNA that are dominated by protein residing at tight binding sites (e.g., operator sites) (Supplementary Fig. 11). Using a three-state kinetic model previously validated (Fig. 2c)^22^, we can analyze the distribution of *τ* to extract the effective unbinding rate constant *k*_−1_ (Fig. 2d; Supplementary Information 5). This SMT, along with single-cell total fluorescence quantitation, also enabled us to quantify the protein copy number, from which protein concentration in the cell can be determined (Supplementary Information 1.2.3).

**Fig. 2.**
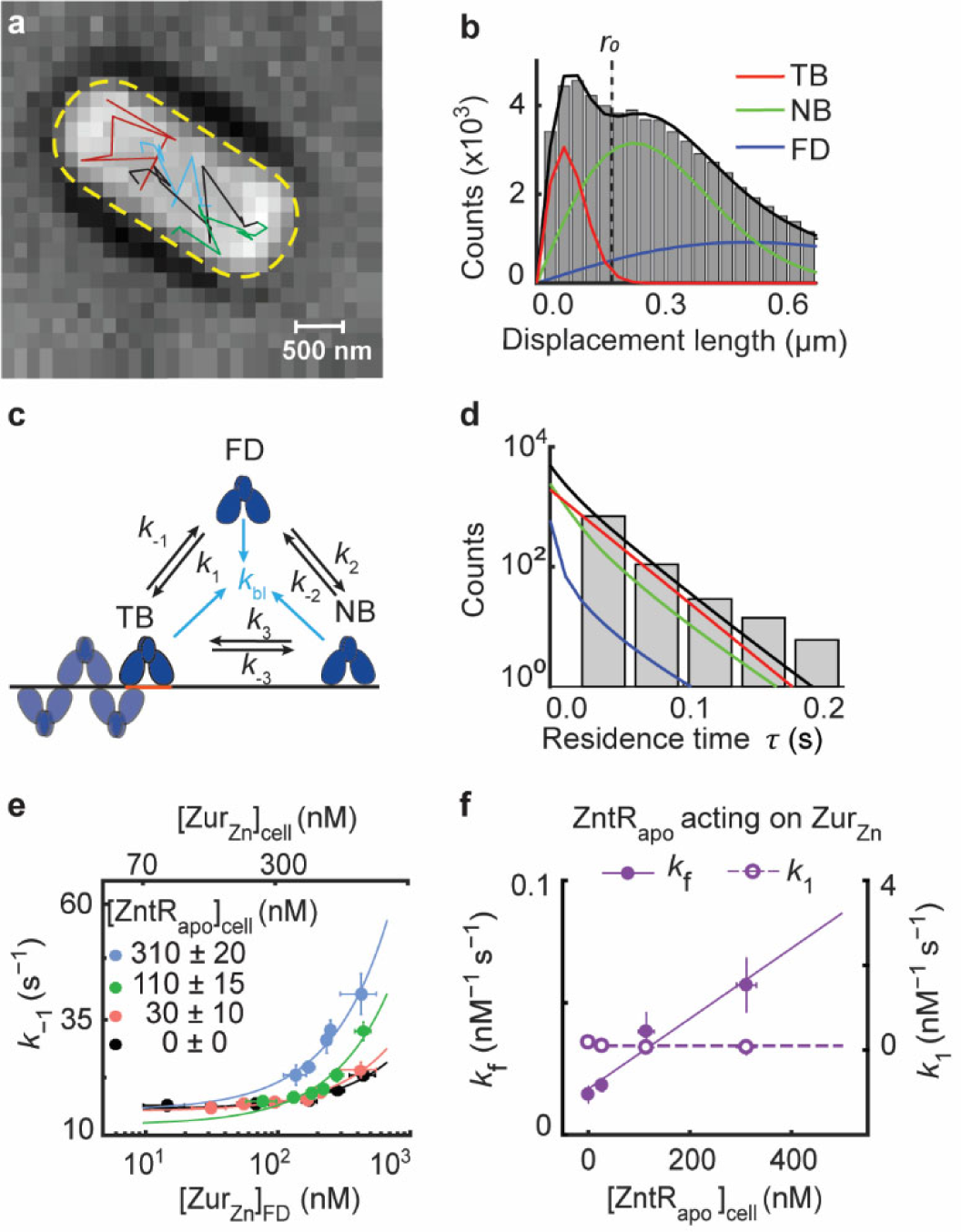
Single-molecule tracking shows that ZntR_apo_ enhances the kinetics of Zur_Zn_ unbinding from DNA in cells. **a,** Exemplary single-molecule 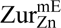 tracking trajectories (colored lines) overlaid on the bright-field transmission image of a live *E. coli* cell from a strain expressing both Zur^mE^ and 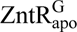 from plasmids grown in the presence of 20 µM Zn^2+^. Yellow dashed line: cell contour. **b,** Exemplary distribution of displacement length *r* per time lapse (40 ms) for 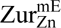 at a total cellular concentration of ∼500 nM from ∼4000 cells from a strain expressing both Zur^mE^ and 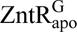 from plasmids grown in the presence of 20 µM Zn^2+^. Solid lines: the overall fit (black), and the resolved TB, NB, and FD diffusion states (Supplementary Information 4). Vertical dashed line: the displacement threshold *r*_0_ below which >99.5% TB state is included. **c**, 3-state kinetic model for Zur-DNA interactions in cell. *k*’s: rate constants. **d**, Histogram of microscopic residence time *τ* of 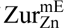 in cells having ∼250 nM Zur^mE^ and ∼30 nM 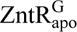. Black line: fit with Supplementary Eq. S11; the deconvoluted contributions from the three diffusion states are in red, green and blue as color-coded in B. **e**, Dependence of the effective unbinding rate constant *k*_−1_ of 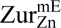 on its own concentration and at different 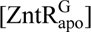 in the cell. Lines: fits with Equation (1) including 1^st^ and 3^rd^ terms only. **f**, The facilitated unbinding rate constant *k*_f_ and the binding rate constant *k*_1_ of 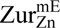 vs. cellular 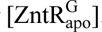. Solid line: linear fit; dashed line: horizontal line fit. Error bars in (e-f) are SEM.

For ZntR_apo_, we used the C115S mutation to remove its metal binding, making it permanently apo^36^. We further tagged ZntR_apo_ with super-folder GFP (i.e., 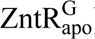) in an inducible plasmid on top of Δ*zntR* in the chromosome. This GFP tagging maintains ZntR’s function (Supplementary Information 3) and allows for spectral separation from the red fluorescent form of 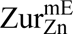 and for quantification in the cell.

By sorting individual cells into groups of similar cellular 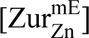 and 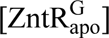 concentrations and analyzing each group separately (Supplementary Fig. 7), we overcame large cell-to-cell protein expression heterogeneity and determined the apparent unbinding rate constant *k*_−1_ from DNA for 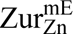 as a function of its cellular concentration and at different cellular 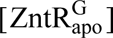 (Fig. 2e). In the Δ*zntR* strain with no 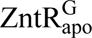 present, *k*_−1_ of 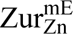 increases with increasing free (and total) 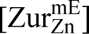 in the cell (Fig. 2e, black), reflecting its facilitated unbinding, as we reported previously (Fig. 5f, steps 1, 2, and 3)^22^, where the slope is the facilitated unbinding rate constant *k*_f_ (Equation 1). The impeded unbinding for Zur_Zn_ occurs at lower than accessible concentrations in the cell and is unobservable here^22^. Strikingly, with increasing 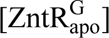, *k*_−1_ of 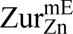 progressively increases (Fig. 2e), and its slope, *k_f_*, increases linearly with 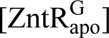 (Fig. 2f, solid symbols), suggesting that ZntR_apo_ can enhance the facilitated unbinding of Zur_Zn_.

The kinetic model in Fig. 2c also allowed for analyzing the fractional populations of Zur’s three states, leading to extraction of other kinetic parameters (Extended Data Table 1; Supplementary Information 5.2)^22^, including 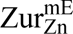’s binding rate constant *k*_1_ to tight binding sites. Interestingly, *k*_1_ shows no dependence on 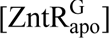 (Fig. 2f, open symbols). This independence indicates that ZntR_apo_ does not block Zur_Zn_ binding to operator sites and suggests that ZntR_apo_ does not bind on its own to the partial recognition sequences that overlap with Zur boxes on DNA, but instead requires Zur to be bound on DNA, possibly involving ZntR-Zur interactions.

### Zur_Zn_-DNA interactions are dynamic *in vitro*

The above results show that the unbinding of the Zn-uptake repressor Zur_Zn_ from DNA can be enhanced by the apo-repressor form of the Zn-efflux regulator ZntR_apo_ in the cell. Given the complexity of cellular environments, this enhancement may or may not be from direct actions of ZntR_apo_ on DNA-bound Zur_Zn_. To remove cellular complexity, we examined whether and how ZntR_apo_ can directly affect Zur_Zn_ unbinding from its cognate DNA *in vitro*. Here we labeled a 31-bp DNA at one end with the FRET donor Cy3, immobilized it on a surface via biotin-neutravidin linkage at the other end, and examined its interaction with Cy5-labeled Zur_Zn_ (a homodimeric protein) in the surrounding solution using single-molecule FRET (smFRET) measurements (Fig. 3a; Extended Data Fig. 2b; Supplementary Information 1.3). The DNA sequence was from the bidirectional promoter of *E. coli*’s *znuCB/znuA* genes, encompassing the two-dyad Zur binding box and a potential partial ZntR recognition sequence (Fig. 3a, top). We also constructed a 22-bp DNA, in which one dyad is truncated to allow only one Zur dimer to bind (Fig. 3a, lower).

**Fig. 3.**
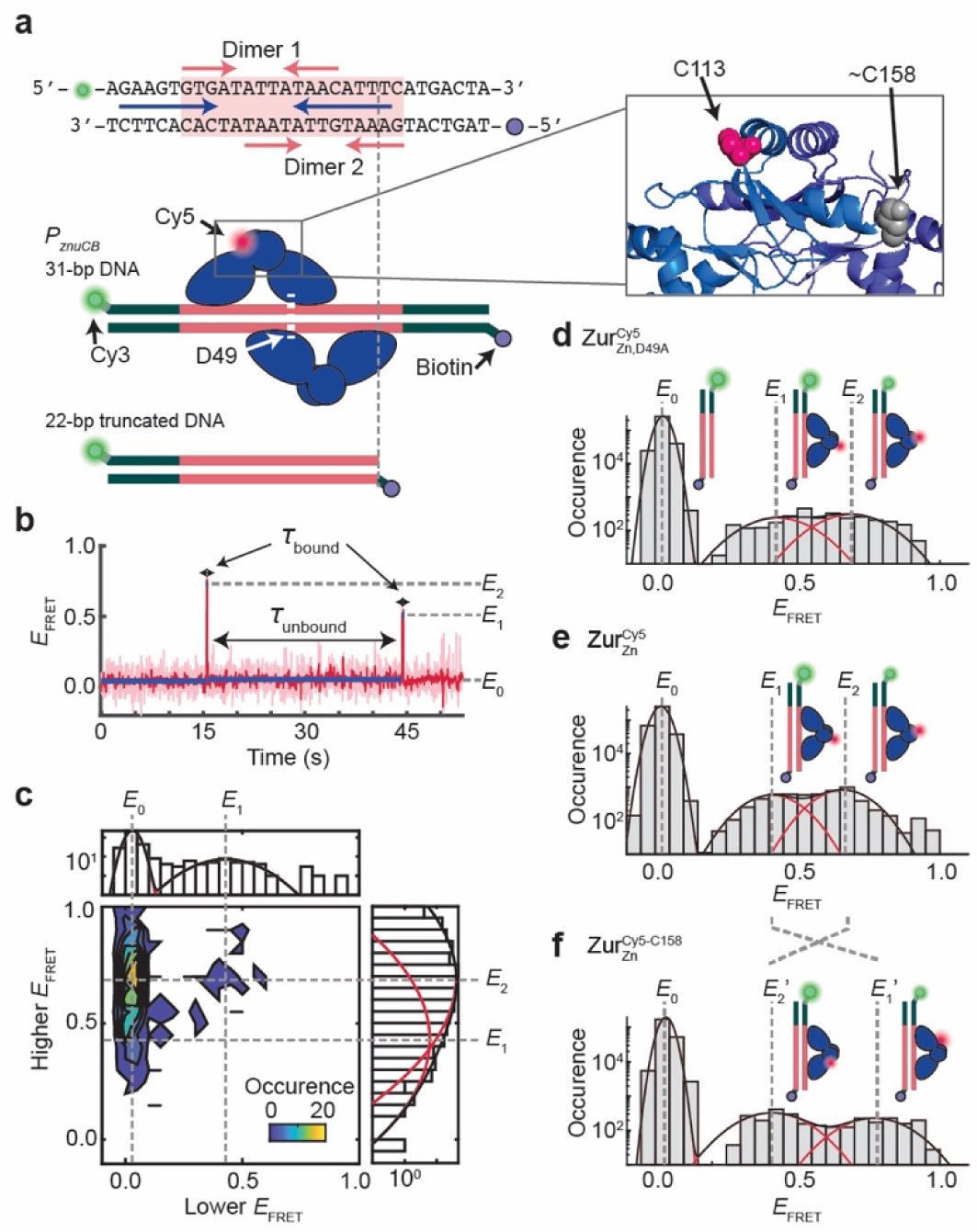
SmFRET measurements show dynamic Zur_Zn_-DNA interactions *in vitro*. **a,** Top: the DNA oligomer sequence from *E. coli znuCB* promoter and Cy3/biotin label positions for smFRET measurements. Pink shade: Zur box; pink arrows: two dyads for binding two Zur_Zn_ dimers; blue arrows: potential ZntR recognition dyad (Fig. 1b). Bottom: Two different-length DNA constructs. A single Cy5 is labeled at C113 or C158 on Zur. Zoom-in box: C113 and approximate C158 positions (shown on different monomers for clarity) in the dimeric Zur_Zn_’s crystal structure (PDB: 4MTD^32^; residues 153-171 are unresolved). **b,** Representative single-molecule *E*_FRET_ trajectory of an immobilized 22-bp truncated DNA^Cy3^ interacting with 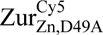 (Cy5 at C113) (4 nM). Pink line: raw data; red line: after non-linear filtering; blue line: mean value of each *E*_FRET_ state. Three *E*_FRET_ states (*E*_0_, *E*_1_, *E*_2_) are denoted. Dwell times on *E*_0_ state are designated as *τ*_unbound_ and those on higher states (*E*_1_ and *E*_2_) as *τ*_bound_. **c,** Two-dimensional histogram of the lower vs. higher *E*_FRET_ state values from single-molecule *E*_FRET_ trajectories, as in (b). Top/right: corresponding one-dimensional projections; red (black) lines: Gaussian resolved (overall) fits. **d,** Histogram of *E*_FRET_ trajectories, as in (b). **e,** Same as (d), but with 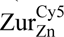 (Cy5 at C113) (4 nM). **f,** Same as (d), but with 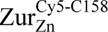 (4 nM). Cartoons in (d-f) show free DNA and DNA-bound Zur_Zn_ in two binding orientations differentiated by the Cy5-label. The FRET donor (green sphere) and acceptor (red sphere) are drawn on DNA and Zur at their approximate locations. All histograms are compiled from >300 *E*_FRET_ trajectories; bin size = 0.05.

First, we studied this truncated 22-bp DNA^Cy3^ interacting with the mutant 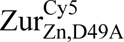; this D49A mutation eliminates the key inter-dimer salt-bridge interactions, resulting in dominantly single dimer-DNA interactions at up to 10 nM Zur_Zn_ concentrations^32^. The single Cy5 label is at C113 of one monomer of the dimeric protein (unless otherwise noted), and C113 is a non-conserved surface-exposed natural cysteine distant from Zur’s DNA-binding domains (Fig. 3a, inset; Extended Data Fig. 3; Supplementary Information 6.1). At 4 nM 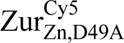, single-molecule *E*_FRET_ trajectories show distinct states and their dynamic transitions: some show three *E*_FRET_ states (Fig. 3b), but most show two states during the limited observation time before label photobleaching (Supplementary Fig. 21). We pooled hundreds of such trajectories and examined the two-dimensional histogram of lower vs. higher *E*_FRET_ values: three states are clearly resolved at *E*_FRET_ ∼0.03, 0.43, and 0.69, denoted as *E*_0_, *E*_1_, and *E*_2_, respectively (Fig. 3c). The same three states also fit the one-dimensional *E*_FRET_ histogram (Fig. 3d): the lowest *E*_FRET_ state (*E*_0_) is assigned as the free DNA state, which dominates the distribution expectedly. The *E*_1_ and *E*_2_ states can be assigned as the two orientations of a single 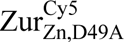 dimer bound to the truncated DNA^Cy3^, where the single Cy5-label breaks the symmetry (Fig. 3d, cartoons). Consistently, their relative populations are almost equal (slight difference could be due to the single Cy5 labeling).

We then studied 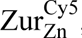, which contains the natural D49 residue for inter-dimer salt-bridge interactions, interacting with the truncated DNA^Cy3^. Expectedly, 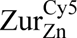 behaves similarly as 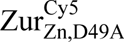, showing the same *E*_FRET_ states (Fig. 3e), because the truncated DNA^Cy3^ only allows for one Zur dimer binding regardless whether or not Zur can form inter-dimer interactions. To further improve data analysis reliability, we globally fitted the results of 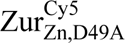 and 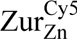 interacting with the truncated DNA^Cy3^ to obtain *E*_1_ and *E*_2_ values at ∼0.44 and ∼0.65, respectively (Supplementary Information 7), which agree with predictions from the Zur_Zn_-DNA complex structure (Supplementary Information 6.2; Supplementary Table 7).

To further confirm the assignment of *E*_1_ and *E*_2_ states, we moved the Cy5 from C113 to C158, another non-conserved surface-exposed natural cysteine (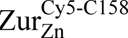; Fig. 3a, inset). Based on the Zur_Zn_-DNA complex structure, this C158 position is closer to the FRET donor in the complex for the *E*_1_ orientation but further for the *E*_2_ orientation (Extended Data Fig. 4a-c; Supplementary Information 6.2). Indeed, *E*_1_ increases to ∼0.77 (*E*_1_′ state) and *E*_2_ decreases to ∼0.41 (*E*_2_′ state), while maintaining the same population ratio (Fig. 3e-f).

### ZntR_apo_ enhances facilitated unbinding of Zur_Zn_ from DNA *in vitro*: a ‘through-DNA’ mechanism

Next, we used the 31-bp DNA^Cy3^, which contains the full two-dyad Zur box to maximally bind two Zur_Zn_ dimers and also encodes the potential ZntR recognition sequence (Fig. 3a). For 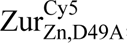, the salt-bridge mutant that has weakened inter-dimer interactions, the *E*_FRET_ histogram can be fitted by four protein-bound states (Supplementary Information 7), with approximately equal populations, besides the free DNA state *E*_0_ at 1 nM protein concentration (Fig. 4a). Two of them, *E*_1_ and *E*_2_, are the same as those in 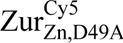’s interaction with the truncated DNA^Cy3^ (Fig. 3d) and are thus assigned similarly as the two orientations of a single 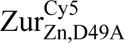 dimer bound at the dyad proximal to Cy3. The other two (*E*_3_ and *E*_4_) have lower *E*_FRET_ values (∼0.27 and ∼0.43) and can be assigned as the two orientations of a single 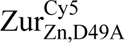 dimer bound at the dyad distal to Cy3 (Fig. 4a, cartoons). All *E*_FRET_ values agree with predictions from the Zur_Zn_-DNA structure (Supplementary Information 6.2; Supplementary Table 7). Upon increasing 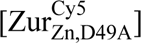 to 4 nM, a new *E*_FRET_ state appeared at ∼0.8 (*E*_7_; Fig. 4b), attributable to two 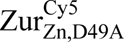 dimers concurrently bound to the 31-bp DNA, now possible due to increased protein concentration. Consistently, when we swapped out 75% of the 4 nM 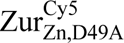 to its unlabeled form, the *E*_FRET_ ∼0.8 peak disappeared (Extended Data Fig. 5).

**Fig. 4.**
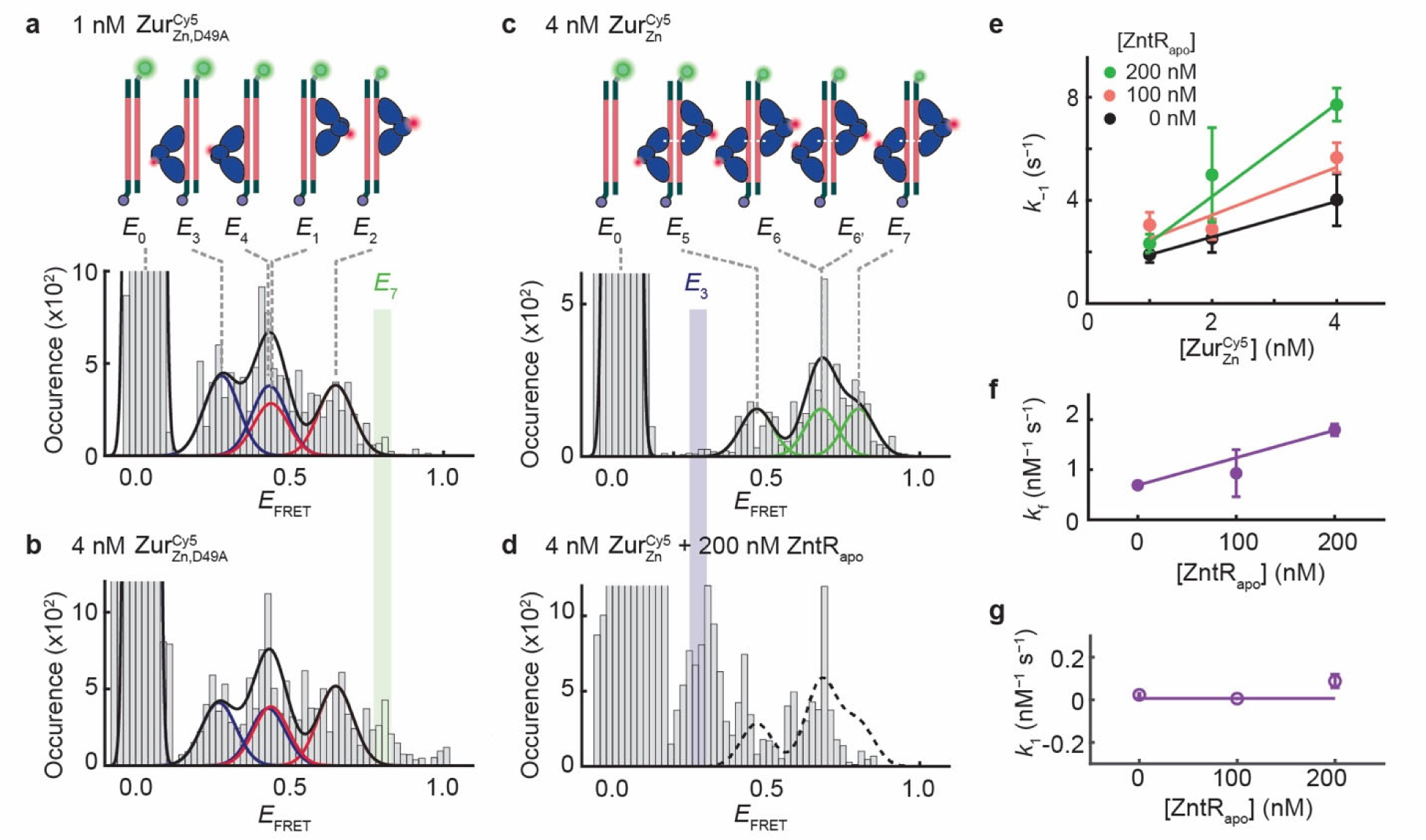
ZntR_apo_ enhances facilitated unbinding of Zur_Zn_ from DNA *in vitro*. **a,** Histogram of *E*_FRET_ trajectories of immobilized 31-bp DNA^Cy3^ interacting with 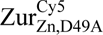 (1 nM). Colored lines: Gaussian resolved fits, where each color corresponds to one 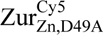 dimer at one of the two dyads of Zur box in two orientations, as shown by the inset cartoons. Black line: overall fit. Bin size = 0.02. **b,** Same as (a), but with 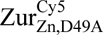 at 4 nM. *E*_7_ state (green-shaded) only appears when two 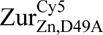 dimers are bound to DNA. **c,** Same as (a), but with 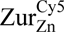 (4 nM). Cartoons show DNA-bound 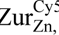 with four different combinations of two dimer-bound form. **d,** Same as (c) but with an introduction of unlabeled ZntR_apo_ (200 nM). *E*_3_ state (blue-shaded) only appears when a single Zur_Zn_ dimer is bound to DNA. **e,** Apparent unbinding rate constant *k*_−1_ of 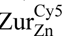 on the 31-bp DNA vs. its own concentration at 0, 100, 200 nM of ZntR_apo_. Lines: linear fits; the slope is the facilitated unbinding rate constant *k*_f_ for 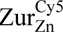. **f,** ZntR_apo_-concentration-dependent *k*_f_ for 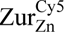 extracted from (e). Line: linear fit. **g,** The binding rate constant *k*_1_ for 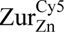 to 31-bps DNA shows no significant dependence on [ZntR_apo_]. Line: horizontal line fit. All error bars are 95% confidence intervals from fits.

Then, we studied the 31-bp DNA^Cy3^ in interacting with 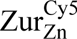. At 4 nM, 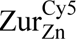 is expected to bind as two dimers because of inter-dimer salt-bridge interactions^32^. Consistently, at *E*_FRET_ ∼0.27, the lowest *E*_FRET_ state for a single dimer-bound form (i.e., *E*_3_ in Fig. 4a-b) and where the two-dimer-bound form should not have a FRET signal (Supplementary Information 6.2; Supplementary Table 7), no population was observed (Fig. 4c). The higher *E*_FRET_ states can be resolved into three states with a 1:2:1 population ratio at *E*_FRET_ ∼ 0.47, 0.68, and 0.80, denoted as *E*_5_, *E*_6_ (*E*_6_′), and *E*_7_, respectively, and assigned as the four different combinations of two-dimer-bound form, each dimer carrying a single Cy5 label (Fig. 4c, cartoons). Note that the *E*_6_ and *E*_6_′ states are unresolved from each other in the *E*_FRET_ histogram. All these *E*_FRET_ values also agree with the predictions from Zur_Zn_-DNA complex structure (Supplementary Information 6.2; Supplementary Table 7). Moreover, the *E*_7_ state is at the same position as the additional *E*_FRET_ ∼0.8 peak in Fig. 4b, when the salt-bridge mutant 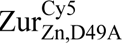 is at a higher concentration, supporting it being from the two-dimer-bound form.

We next added ZntR_apo_ (i.e., ZntR_C115S_ mutant) to probe whether it can enhance 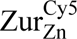 unbinding from the 31-bp DNA^Cy3^. Strikingly, *E*_7_, a two-dimer-bound state that does not overlap with any of the single-dimer-bound states, is substantially depopulated, while the *E*_3_ state, unique to single-dimer-bound form, appears (Fig. 4d; Extended Data Fig. 6g-l). Both observations indicate that ZntR_apo_ disrupts Zur_Zn_-DNA interactions. Moreover, the states at higher *E*_FRET_ values (i.e., *E*_FRET_ ∼ 0.5-0.7) are more depopulated than those at lower *E*_FRET_ values (i.e., *E*_FRET_ ∼ 0.3-0.5), which is even clearer at lower 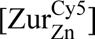 (Supplementary Information 8). Therefore, the 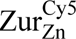 dimer bound at the proximal dyad to Cy3 is preferentially disrupted by ZntR_apo_, consistent with the fact that the partial ZntR_apo_ recognition sequence on the 31-bp DNA^Cy3^ is proximal to Cy3 (Fig. 3a).

We further analyzed the residence times *τ*_bound_ and *τ*_unbound_ from the *E*_FRET_ trajectories to extract Zur^Cy5^-DNA^Cy3^ interaction kinetics (Fig. 3b; Extended Data Table 2; Supplementary Fig. 23-Supplementary Fig. 24). The unbinding rate constant *k*_−1_ for 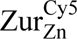 on the 31-bp DNA clearly shows a linear dependence on its own concentration (Fig. 4e, black), reflecting its facilitated unbinding that was observed in cells (Fig. 2e), where the slope is the facilitated unbinding rate constant *k*_f_^22^. Moreover, with increasing [ZntR_apo_], *k*_f_ increases (Fig. 4e-f), as observed in cells (Fig. 2f, solid symbols), indicating that ZntR_apo_ directly enhances the facilitated unbinding of Zur_Zn_ from DNA. Meanwhile, the binding rate constant *k*_1_ of 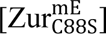 is independent of ZntR_apo_ (Fig. 4g), suggesting that ZntR_apo_ alone does not bind directly to this 31-bp DNA, but needs incumbent Zur_Zn_ on DNA, similarly as observed in cells (Fig. 2f, open symbols). These results further support that possible ZntR-Zur interactions are important for ZntR_apo_-enhanced Zur_Zn_ unbinding from DNA.

Altogether, the *in vitro* experiments demonstrate that ZntR_apo_ enhances Zur’s facilitated unbinding from DNA through its direct actions on Zur-DNA complex, enabled by the overlapping Zur and ZntR recognition sequences, where both protein-DNA and protein-protein interactions are important. We postulate that this ‘through-DNA’ mechanism possibly occurs via ZntR_apo_ acting directly on the Zur-DNA ternary complex to form a heteromeric ZntR-Zur-DNA quaternary complex (Fig. 5f, step 5).

**Fig. 5.**
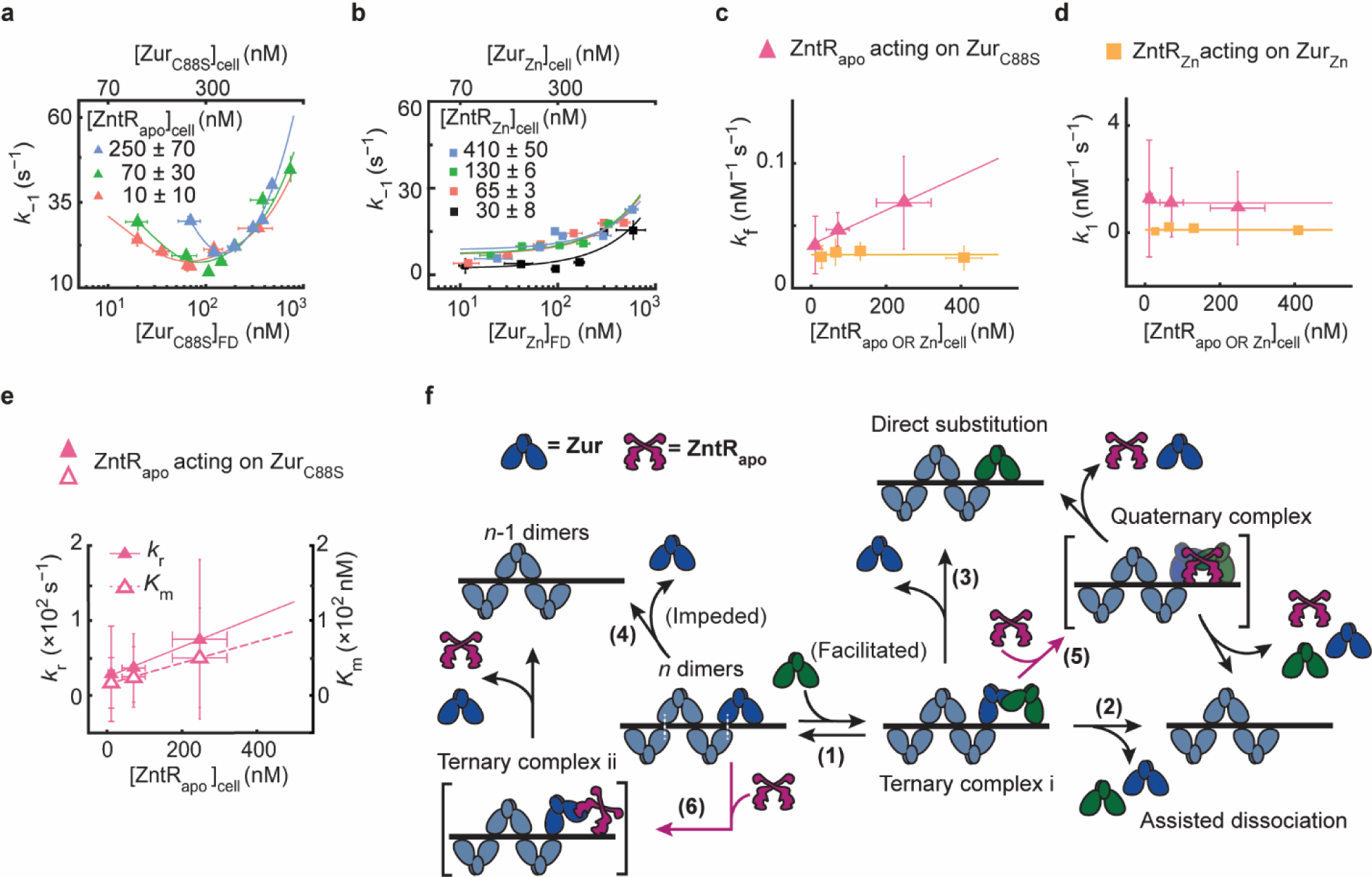
ZntR effects on Zur-DNA interactions. **a,** Dependence of the effective unbinding rate constant *k*_−1_ of 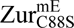 on its own concentration and at different 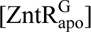 in the cell. Lines: fits with Eq. (1). **b,** Same as (a), but for *k*_−1_ of 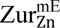 and at different 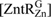 in the cell. Lines: fits with Eq (1) including 1^st^ and 3^rd^ terms only. **c,** The facilitated unbinding rate constant *k*_f_ of 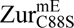 vs. cellular 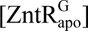 (magenta) and of 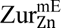 vs. the cellular 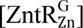 (yellow). Lines: linear (magenta) and horizontal line (yellow) fits. **d**, Same as (c) but for the binding rate constant *k*_1_. Lines: horizontal line fits. **e,** The impeded unbinding rate constant *k*_r_ and the effective oligomer dissociation constant *K*_m_ of 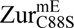 vs. cellular 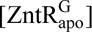. Lines: linear fits. Error bars in (a-e) are SEM. **f,** Mechanistic model for ZntR_apo_-dependent Zur unbinding kinetics. Starting with oligomerized Zur (dark and light blue) at a tight-binding site on DNA, the unbinding of an incumbent Zur protein (dark blue) can be facilitated by a freely diffusing Zur (dark green) through the formation of a ternary complex i (step 1), leading to assisted dissociation (step 2) or direct substitution (step 3); this facilitated unbinding of Zur can be enhanced by ZntR_apo_ through the formation of a heteromeric quaternary complex (step 5). The oligomer-induced impedance of Zur unbinding (step 4) can be weakened by ZntR_apo_ through the formation of a heteromeric ternary complex ii (step 6), leading to faster Zur unbinding as well. White dashed lines on the ‘*n* dimers’ denote salt bridge interactions between Zur dimers.

### ZntR_apo_ enhances unbinding of Zur non-repressor form as well

Having shown that ZntR_apo_ directly acts on Zur_Zn_-DNA complex to enhance Zur_Zn_ unbinding from its recognition sites, we continued to examine whether ZntR_apo_ can act on the DNA-bound non-repressor form of Zur, as these two coexist in the cell under Zn deficient (and replete) conditions (Fig. 1a). To ensure Zur being in the non-repressor form in the cell, we examined the Zur_C88S_ mutant whose Cys88 at the regulatory Zn-binding site was mutated to make it a constitutive non-repressor^1,32^ and which binds tightly to DNA but at unidentified sites distinct from Zur boxes at its regulon promoters^22^. We again sorted individual cells into groups of similar 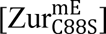 and 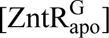 and analyze the groups separately.

At any cellular 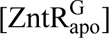, the apparent unbinding rate constant *k*_−1_ of 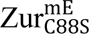 shows the biphasic unbinding behavior (Fig. 5a): initially decreases with increasing cellular 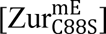 (i.e., impeded unbinding), reaches a minimum, and then increases toward higher concentrations (i.e., facilitated unbinding), as we previously discovered and described by Equation 1^22^. Strikingly, with increasing 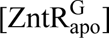, the biphasic behavior of *k*_−1_ shifts toward the upper-right of the plot (Fig. 5a). The extracted facilitated unbinding rate constant *k*_f_ of 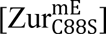 increases (Fig. 5c, magenta), similarly as observed for Zur_Zn_ above, and consistent with ZntR_apo_ acting on DNA-bound Zur non-repressor form, possibly through the same heteromeric quaternary complex (Fig. 5f, step 5).

The impeded unbinding rate constant *k*_r_ of 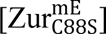 also increases with increasing 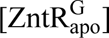 (Fig. 5e, solid symbols); so does *K*_m_, the effective dissociation constant of Zur oligomerization on DNA (Fig. 5e, open symbols), which correlates with the minimum position of *k*_−1_ in Fig. 5a. Both changes of *k*_r_ and *K*_m_ indicate that ZntR_apo_ can act on oligomerized Zur_C88S_ on DNA, weakening its oligomerization and diminishing its impedance on unbinding. We attribute this weakening to ZntR_apo_ directly interacting with oligomerized Zur_C88S_, for example via a possible heteromeric ternary complex (Fig. 5f, step 6). Since the non-repressor Zur_C88S_ binds to sequences outside Zur’s regulon promoters, ZntR_apo_’s effects on Zur_C88S_-DNA interaction suggest that ZntR recognition sites must exist at other places on the chromosome as well. Indeed, we discovered previously >80 potential ZntR recognition sites across the *E. coli* chromosome^23^.

The binding rate constant *k*_1_ of 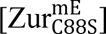 shows no dependence on 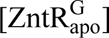 (Fig. 5d, magenta), suggesting that ZntR_apo_ does not block Zur_C88S_ binding to DNA and corroborating that ZntR_apo_’s interaction with the Zur-DNA complex involves ZntR-Zur interactions besides ZntR-DNA interactions (Fig. 5f, steps 5 and 6).

Combining the results on Zur_Zn_ and Zur_C88S_, ZntR_apo_ enhances the unbinding of Zur from DNA, regardless of whether Zur binds tightly to Zur boxes or other sequences, in a ‘through-DNA’ mechanism. Using the kinetic model in Fig. 5f, we derived the relation between Zur’s unbinding rate constant *k*_−1_ and the concentrations of Zur and ZntR_apo_ (Supplementary Eq. S39; Supplementary Information 9). Supplementary Eq. S39 satisfactorily describes the experimental data, further supporting the ‘through-DNA’ mechanism for ZntR_apo_ acting on DNA-bound Zur.

### ZntR_Zn_ has no effect on repressor Zur_Zn_ unbinding from DNA

Having shown that ZntR_apo_ can enhance the unbinding of both the repressor and non-repressor forms of Zur from DNA, we pondered about the metallated ZntR_Zn_, which is the activator for Zn efflux. As ZntR_Zn_ and Zur_non-repressor_ do not coexist in the cell under physiological conditions (Fig. 1a), we examined ZntR_Zn_’s effect on the unbinding of Zur_Zn_, which coexist in the cell under Zn excess conditions (i.e., in the presence of 100 μM Zn^2+^ in the media^22,36^; Fig. 1a, right). At a given 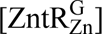 in the cell, the apparent unbinding rate constant *k*_−1_ of 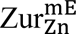 expectedly shows facilitated unbinding (Fig. 5b). More importantly, changing ZntR_Zn_’s cellular concentration has no discernible effect on the facilitated unbinding rate constant *k*_f_ of 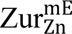, in contrast to that of

ZntR_apo_ (Fig. 5c, yellow vs. magenta). Therefore, ZntR’s interaction with Zur-DNA complex only applies to ZntR’s apo-repressor form and not its holo-activator form. This difference could come from that apo and holo ZntR have different conformations^21^, which may lead to their different interactions with Zur-DNA complex. Consistently, the binding rate constant *k*_1_ of 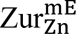 shows no dependence on 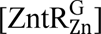, either (Fig. 5d, yellow).

## Discussion

In bacteria’s natural habitats, the availability of essential micronutrients like Zn^2+^ can fluctuate substantially. The ability of the efflux regulator ZntR_apo_ in enhancing the unbinding of both the repressor and non-repressor forms of the uptake regulator Zur from DNA could have functional significance in facilitating the switching in Zn uptake regulation when an *E. coli* cell encounters changing Zn environments. Starting from Zn replete conditions where Zur_Zn_ repressor and ZntR_apo_ coexist in the cell (Fig. 1a, center), if the cell encounters a Zn deficient environment, the enhancement of Zur_Zn_ unbinding from its regulon promoters by free ZntR_apo_ would facilitate the derepression of Zn uptake genes. On the other hand, starting from Zn deficient conditions where Zur’s non-repressor form coexists with ZntR_apo_ in the cell (Fig. 1a, left), if the cell moves into Zn replete or excess conditions, ZntR_apo_-enhanced Zur unbinding from non-operator sites would facilitate Zur release from DNA for binding to operator sites, upon Zn-metallation, to repress Zn uptake. Within ZntR’s physiological concentration range in the cell, ZntR_apo_ can enhance the apparent unbinding rate constant of Zur_Zn_ by ∼130% and that of Zur_C88S_ (a non-repressor) by ∼50% (Extended Data Fig. 7; Supplementary Information 10). Therefore, such facilitation by ZntR_apo_ could have significant kinetic effects in transcription regulation by Zur.

The through-DNA actions of an efflux regulator on its corresponding uptake regulator, or vice versa, likely extend beyond *E. coli* to other bacterial species and other metal ion homeostasis (Supplementary Information 2). We find that partial recognition sequences of ZntR or its homolog also exist around Zur boxes in other bacteria, such as in *S. typhimurium* and *P. aeruginosa* (Fig. 1b; Extended Data Fig. 1a). Such sequence overlaps also occur to other Zn efflux-uptake regulator pairs, including the CzrA-Zur pair in *B. subtilis* and the SczA-AdcR pair in *P. aeruginosa* (Extended Data Fig. 1b-c), as well as to bacterial efflux-uptake regulator pairs for other metals like Fe and Ni (Extended Data Fig. 1d-e). Moreover, opposite to the pattern of partial efflux regulator recognition sequences near uptake regulator boxes, Helmann et al. identified two potential Fur (Fe uptake regulator) recognition sites around a known PerR (Fe efflux regulator) binding box^37^. Similar through-DNA actions between regulator pairs could even exist in yeast, a higher organism, as partial efflux regulator recognition sequences can be found near promoter binding boxes of uptake regulators for iron homeostasis (Extended Data Fig. 1f). Altogether, these observations of sequence overlaps on DNA suggest a broad relevance of the through-DNA mechanism between uptake and efflux regulators of many metals, across a range of regulatory proteins, and through different levels of organisms, which may constitute another mechanistic paradigm for metal regulation in biology.

## Data availability

All data are available in the main text or the Supplementary Information. Raw data supporting the findings of this study are available upon request.

## Code availability

MATLAB codes are included in the Supplementary Software 1.

## Acknowledgements

This research is supported by NIH Grant GM109993. We thank Dr. Bing Fu of Cornell University for discussions.

## Author contributions

U.K.C. designed and performed live-cell imaging experiments, constructed strains, expressed and purified proteins, performed biochemical experiments, wrote MATLAB codes, analyzed data, and wrote the manuscript; Y.P. designed and performed *in vitro* smFRET experiments, expressed and purified proteins, performed biochemical experiments, wrote MATLAB codes, analyzed data, and wrote the manuscript; K.S. purified and characterized Zur proteins, performed early smFRET and biochemical experiments; W.J. wrote MATLAB codes for smFRET image analysis; C.P.J. constructed Zur expression plasmids, designed mutant Zur variants, and purified Zur proteins; D.H.F. contributed to plasmid constructions; P.C. directed research and wrote the manuscript.

## Ethics declarations

The authors declare no competing interest.

## Methods

### Construction of strains for *in vivo* and *in vitro* experiments

For live cell single-molecule imaging and tracking of Zur as a function of Zur and ZntR concentrations in *E. coli* BW25113 cells, the proteins Zur and ZntR were tagged genetically with photo-convertible protein mEos3.2 and sfGFP, respectively, to separate them spectrally. The *zur* and *zntR* genes were either tagged chromosomally or the tagged genes were encoded into a plasmid rather than in the chromosome, to access a broad range of protein concentrations.

For *in vitro* smFRET measurement, site-directed mutagenesis was used to make Zur variants that contain a uniquely labelable cysteine in each monomer to label Zur with the FRET acceptor Cy5. All Zur variants and ZntR(C115S) mutant were cloned in a pET3a vector, and the proteins were expressed in *E. coli* (BL21 DE3) cells. See details in Supplementary Information 1.

### Protein purification and DNA labeling for *in vitro* smFRET

All Zur variants were purified as previously described^32^. Briefly, the proteins are overexpressed in cells by isopropyl-beta-D-thiogalactopyranoside (IPTG) and then lysed with lysozyme in lysis buffer, followed by freeze and thaw cycle and sonication. The proteins were collected by centrifugation, then the protein was purified by a series of columns. Protein purity was confirmed by SDS-PAGE, quantified using UV measurement at 280 nm, and protein identity was confirmed by mass spectrometry. After then, Cy5 FRET acceptor was labeled at the targeted cysteine in protein via maleimide chemistry. The mono-labeled fraction was purified using an anion exchange column. For ZntR(C115S) mutant, the protein is expressed in the same way as Zur using IPTG, and purified as previously described^21,36^. Briefly, the supernatant was collected after centrifugation and the proteins were precipitated out with 45% saturated (NH_4_)_2_SO_4_ overnight. The precipitated proteins were resuspended in Tris buffer and purified via a series of columns. Protein purity was confirmed by SDS-PAGE, quantified using Bradford assay. Protein identity was confirmed by mass spectrometry. For Cy3 labeling to DNA, the Cy3 and biotin-tagged DNA oligomeric strands were purchased from Integrated DNA Technologies (IDT, Coralville, IA) and dissolved in Buffer, and annealed together. Two types of double-strand DNA (dsDNA) constructs were used. The sequences of both constructs were from the *znuCB* gene promoter and contain the specific two-dyad sequence recognized by two Zur dimers and the complementary. The other construct is truncated DNA that has only one dyad sequence and its complementary. See details in Supplementary Information 1.3.

### Sample preparation, experimental procedure, and data processing for live cell single-molecule tracking and protein quantification

The *E. coli* cells were grown in LB medium overnight and later diluted in minimal media with vitamins, amino acids and glucose. Cells were grown to an OD600 of 0.3 and l-arabinose was used to induce plasmid expression when applicable. Zn^2+^ was used for Zinc stress to a final concentration of 20uM or 100uM. The cells were then washed with the same minimal media, pelleted and added onto an agarose gel pad in a glass slide which was then sandwiched by a glass coverslip pretreated with gold nano-particles as position markers, and sealed with epoxy-glue.

For single-molecule tracking and protein quantification for mEos3.2-tagged Zur was done as reported previously (Extended Data Fig. 2a)^22,23^. First, a 405 nm (1-100 W/cm^2^) laser was used to photo-convert a single mEos3.2 tagged Zur proteins from their green to red emissive forms. The photo-converted protein was then tracked with 561 nm laser (21 kW/cm^2^), with 4 ms exposure time and a time-lapse of 40 ms. This was done for several cycles, to obtain a single molecule tracking movie. To quantify the total Zur concentration in the cells, the 405 nm laser was used to photo-convert all the mEos3.2 proteins to their red fluorescent form. The 561 nm laser was then used to obtain the total red intensity of the cell. This step was repeated multiple times to photo-bleach all red mEos3.2 proteins. After this step, the total ZntR concentration in the cell was quantified using the 488 nm (7 kW/cm^2^) laser to obtain the intensity of all the green sfGFP protein tagged ZntR in the cell.

We used custom MATLAB codes, iQPALM^23^, to analyze the single molecule tracking movies. Here, the candidate single-molecule spots were selected within defined cell boundaries using a two-dimensional Gaussian fitting and for each determined spot, the intensity of the single mEos3.2 was extracted. The total Zur copy number in the cells was obtained by dividing the whole cell mEos3.2 red intensity by the single molecule intensity. The total cellular ZntR concentration was also determined in a similar way from the whole cell sfGFP green intensity and the single sfGFP intensity determined from separate experiments. See Supplementary Information 1.2 for details.

### Resolution and extraction of effective Zur diffusion states in the cells

By determining the centroid position of each single molecule spot using the two-dimensional Gaussian fitting as described above and in Supplementary Information 1.2, we were able to extract the position trajectory of the single molecules, as done previously^22,23^. From these trajectories, we could also obtain the displacement lengths, *r*, of individual mE-tagged Zur proteins.

Further the cells were sorted by their Zur and ZntR concentrations, into similar concentration ranges to overcome large cell-to-cell heterogeneity, and each individual concentration group was analyzed.

The distributions of the displacement lengths at different Zur and ZntR concentrations, were globally fitted with a probability distribution function, using linear combinations of three Brownian diffusion states, assuming a quasi-static approximation^22,23^. From the fitting results, we could extract the effective diffusion coefficients (*D*’s) and their corresponding fractional population (*A*’s). The assignment of the three diffusive states, corresponding to three Zur populations, cytoplasmic diffusion (FD), Tight binding to DNA (TB) or non-specific binding to DNA (NB) were previously reported and rationalized^22^. See Supplementary Information 4 for details.

### Sample preparation, imaging and data analysis for *in vitro* smFRET studies

To immobilize DNA for in vitro studies, quartz slides were first amine-functionalized, followed by coating with biotinylated-polyethylene glycol (PEG) polymers. The biotinylated terminal group forms biotin-neutravidin linkages for immobilizing biotinylated DNA molecules (Extended Data Fig. 2b)^38,39^. Coverslips were also amine-functionalized and coated with PEG polymers. A microfluidic channel was formed by double-sided tape sandwiched between a quartz slide and a borosilicate cover slip. After then, neutravidin, Cy3-labeled biotinylated DNA solution flowed through the channel for immobilization. Then, the Cy5-labeled Zur solution containing an oxygen scavenging system^40^ in the same buffer, and if applicable, containing ZntR_apo_, was flowed for fluorescence imaging.

The single-molecule fluorescence experiments were performed using a prism-type total internal reflection microscope based on an Olympus IX71 inverted microscope, similarly as we previously reported^39,41,42^. The immobilized Cy3-labeled DNA was excited by a continuous-wave circularly polarized 532-nm laser (CrystaLaser, GCL-025-L-0.5%) on the sample. The fluorescence of both Cy3 and Cy5 was collected by a 60× NA 1.2 water-immersion objective and split by a dichroic mirror into two channels using a Dual-View system (Optical Insights). Each channel of fluorescence was further filtered (HQ580-60m or HQ660LP) and projected onto one-half of the imaging area of an EMCCD camera (Andor Ixon DV887) controlled by Andor IQ software. All image analysis was done by custom-written codes in MATLAB (Supplementary Software S1). Individual Cy3 and Cy5 fluorescence intensity trajectories were extracted and the FRET efficiency (*E*_FRET_) was computed as an approximation using the relationship: *I*_Cy5_/(*I*_Cy5_+*I*_Cy3_), where *I*_Cy3_ and *I*_Cy5_ are the fluorescence intensities. In order to obtain higher resolution *E*_FRET_ histograms, a forward-backward non-linear (fnbl) filter was used to reduce the noise in the fluorescence trajectories (Supplementary Fig. 4)^43,44^ and thresholded to distinguish *E*_FRET_ states. *E*_FRET_ value of each state was taken from the original *E*_FRET_ trajectories to avoid value changes by fbnl filtering. See details in Supplementary Information 1.3.

## Supplementary Information

Materials and Methods Supplementary Text Supplementary Figs. 1 to 24

Supplementary Tables 1 to 7

Supplementary References 1 to 84

Supplementary Software 1

## Extended Data Figures

**Extended Data Fig. 1.**
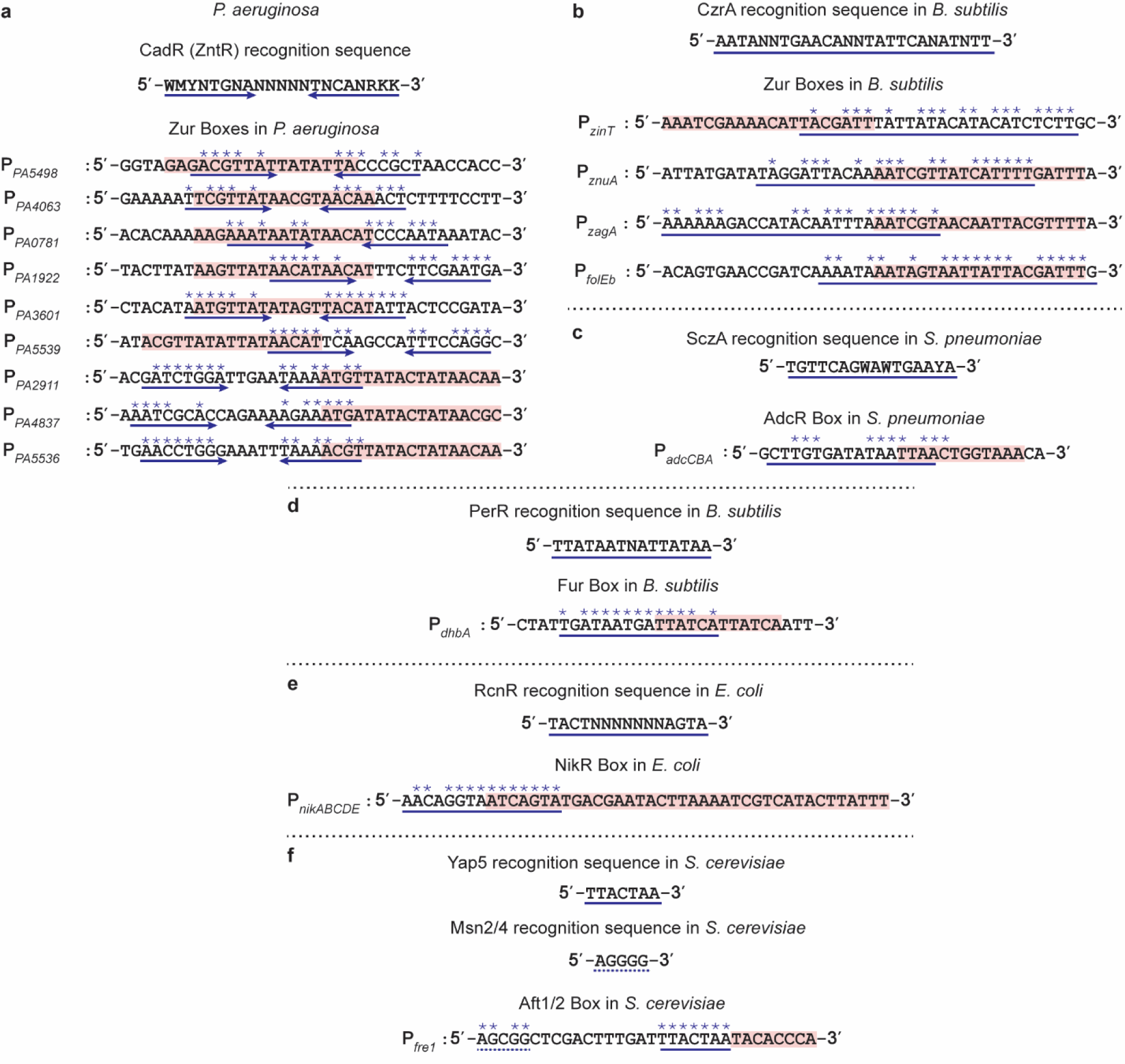
Potential efflux regulator recognition sequences exist around uptake regulator binding boxes (Supplementary Information 2). **a,** Top, Partial recognition sequences of CadR, a ZntR-homolog; bottom, existence of partial CadR recognition sequences around Zur boxes in regulon promoters in *P. aeruginosa.* **b**, Top, the recognition consensus sequence of CzrA, a Zn-efflux regulator; bottom, existence of a partial CzrA recognition sequence around binding box of the Zn-uptake regular Zur in *B. subtilis*. **c**, Top, the recognition consensus sequence of SczA, an Zn-efflux regulator; bottom, existence of a partial SczA recognition sequence around a binding box of AdcR, a Zn-uptake regulator in *S. pneumoniae*. **d**, Top, the recognition consensus sequence of PerR, a Fe-efflux regulator; bottom, existence of a partial PerR recognition sequence around a binding box of Fur, a Fe-uptake regulator in *B. subtilis*. **e**, Top, the recognition consensus sequence of RcnR, a Ni-efflux regulator; bottom, existence of a partial RcnR recognition sequence around a binding box of NikR, a Ni-uptake regulator, in *E. coli*. **f**, Top and middle, the recognition consensus sequence Yap5 (solid blue underline) and that of Msn2/4 (dashed blue underline), both Fe-efflux regulators in *S. cerevisiae*; bottom, existence of a partial Yap5 and Msn2/4 recognition sequence around a binding box of Aft1/2, a Fe-uptake regulator in *S. cerevisiae*. The uptake regulator binding boxes have been highlighted in pink and the potential efflux regulator recognition sequences have been indicated by the blue underline, with each asterisk representing a match with their known consensus recognition sequence. All sequences were obtained from the NCBI genetic database.

**Extended Data Fig. 2.**
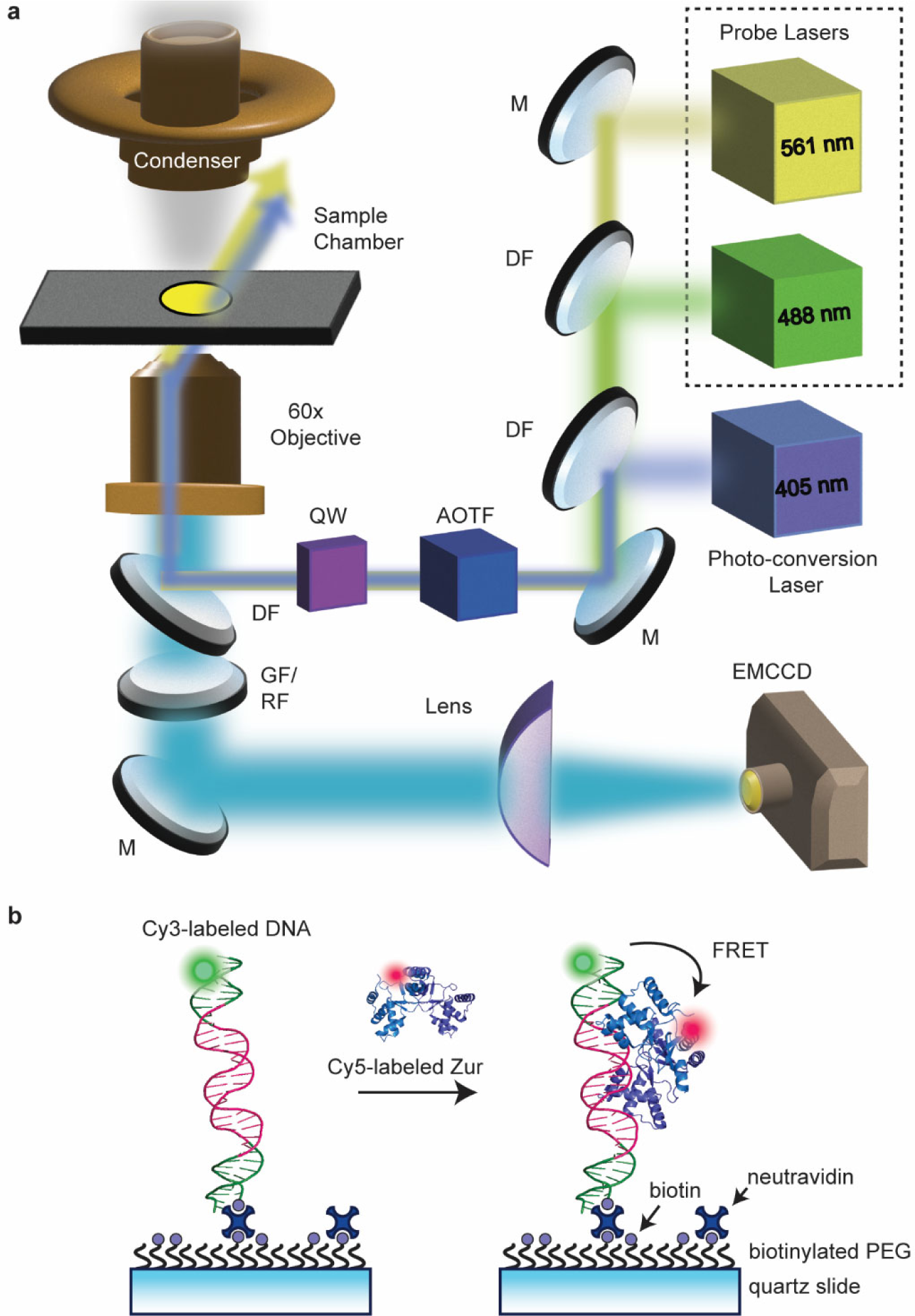
Schematics of PALM microscope setup and experimental scheme of surface immobilization of Cy3-labeled DNA for smFRET measurements. **a,** PALM microscope setup for *in vivo* single molecule tracking and stroboscopic imaging. (M: Mirror; DF: Dichroic Filter; GF/RF: Green Filter/ Red Filter; AOTF: Acoustic Optical Tunable Filter; QW: Quarter Waveplate; b, Scheme of smFRET measurements in vitro. Cy5-labeled Zur is supplied in a continuously flowing solution. Upon Zur binding to DNA, FRET occurs from the donor Cy3 (green sphere) to the acceptor (red sphere). Here, a homo-dimer Zur bound to 31-bp DNA is shown and the Cy5 location corresponds approximately to that in Zur^Cy5^ (Cy5 at C113).

**Extended Data Fig. 3.**
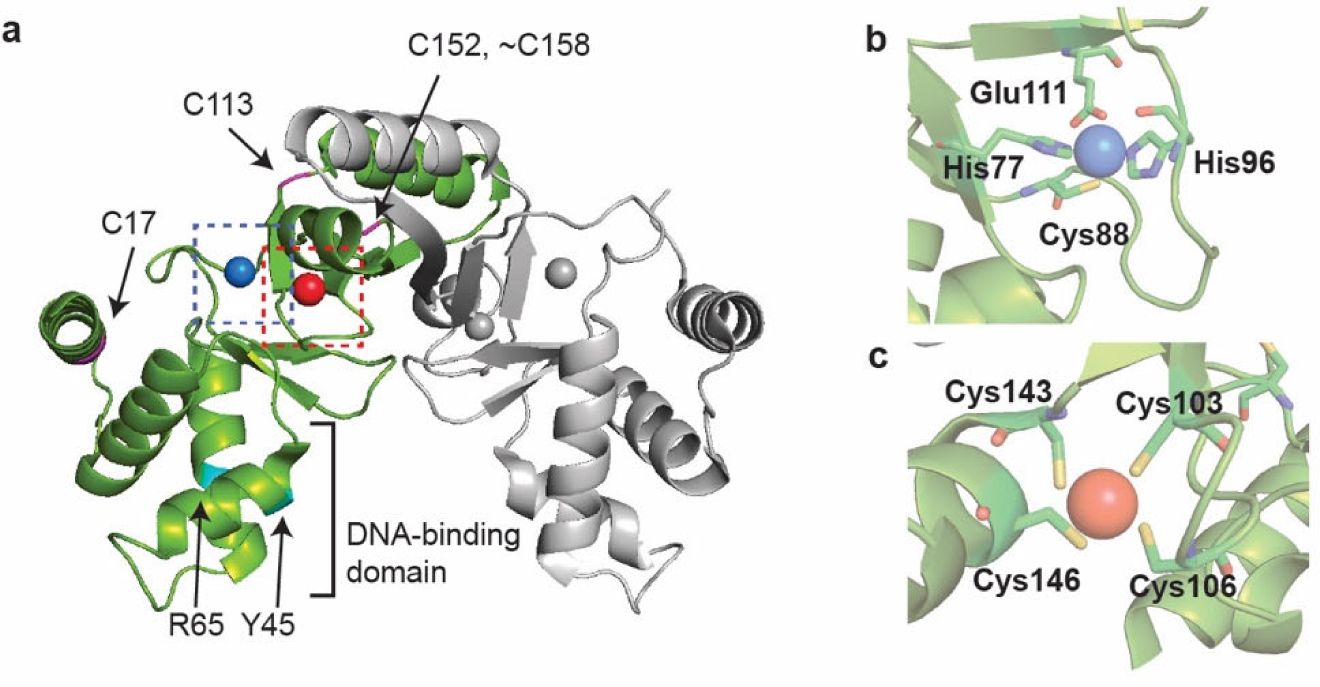
Fluorescent probe location on Zur based on Zur structure (Supplementary Information 6.1). **a,** Crystal structure of the homo-dimeric *E. coli* Zur (PDB: 4MTD^33^), where the two Zur monomers are colored green and gray, and two zinc ions are shown in blue and red spheres (four Zn^2+^ total per dimer). The positions of the four potential labeling sites, which are non-conserved cysteine residues (C17, C113, C152, C158), are colored in magenta and indicated by arrows (C158 location is unresolved in the structure and is denoted approximately together with the C152 location). DNA-binding domain is denoted on the green monomer, and the specific residues that make hydrogen bonds to the DNA bases (Y45, R65) are colored in cyan. In one variant, we used the natural C113 as the Cy5 labeling site, which is far away from Zur’s DNA binding domain, and thus labeling at this position is expected to not interfere with Zur’s DNA binding; the other three non-conserved cysteines (C17, C152, C158) were mutated to serine; we, refer this labeled Zur variant (Cy5 at C113) as Zur^Cy5^. **b-c,** Zoomed-in image of one cysteine (C88) at the regulatory site (b) and four cysteines (C103, C106, C143, C146) at the structural site (c).

**Extended Data Fig. 4.**
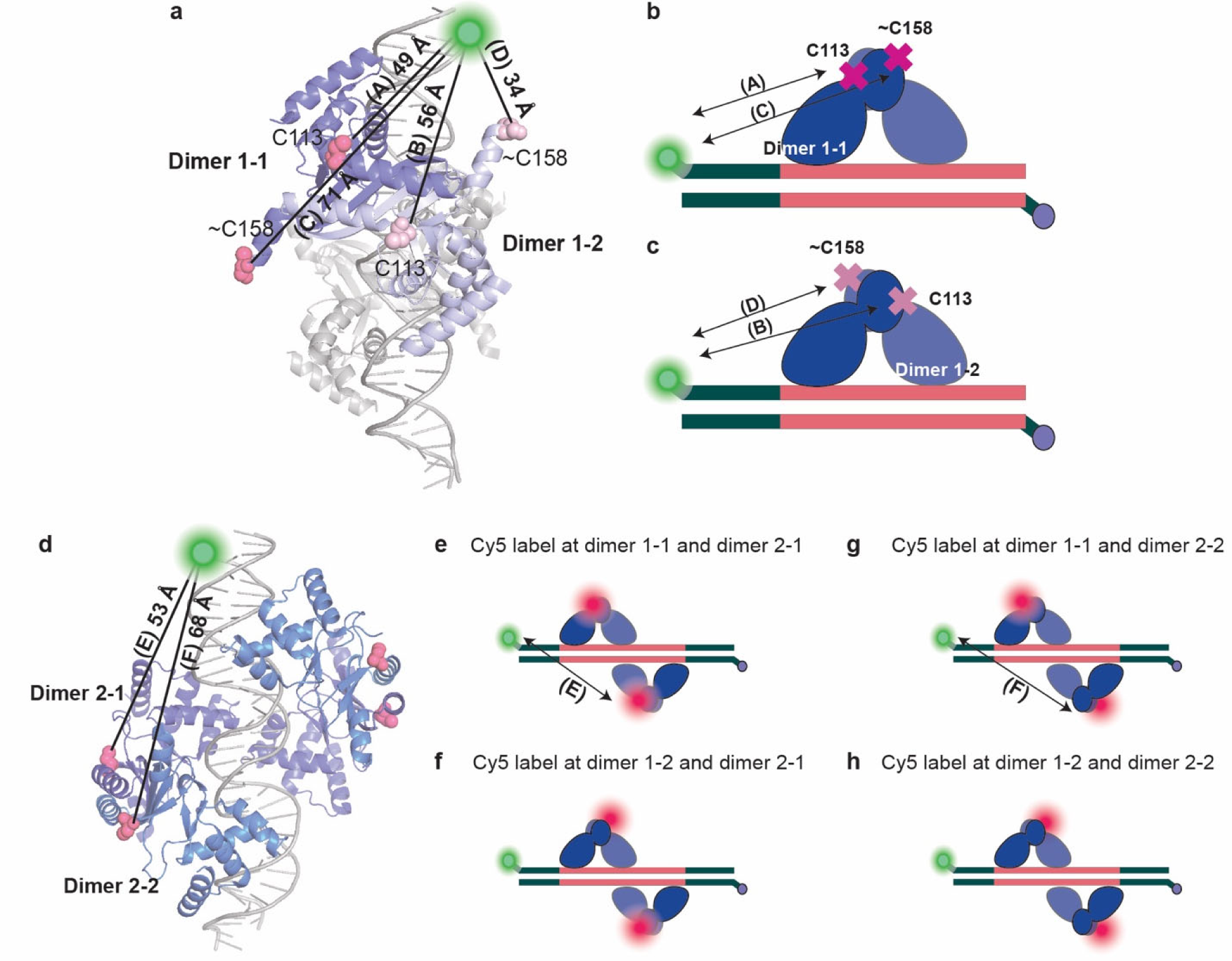
Prediction of *E*_FRET_ value based on Zur-DNA complex structure (Supplementary Information 6.2). **a,** Crystal structure of two holo Zur dimers bound on DNA (PDB: 4MTD^33^), where one dimer Zur is colored purple (Dimer 1-1) and light purple (Dimer 1-2) for the two monomers, and the other dimer of Zur and DNA are colored gray. Two labeling positions (C113 and ∼C158) are shown in pink and light pink spheres for Dimer 1-1 and Dimer 1-2, respectively. **b,** A cartoon showing that one homodimeric Zur is bound to 22-bp truncated DNA with labeling positions colored as pink crosses for Dimer 1-1. The Cy3−Cy5 anchor-to-anchor distances for each labeling position are (A) 49 Å and (C) 71 Å, and the corresponding *E_FRET_* values are ∼0.64 and ∼0.16, respectively. **c,** A cartoon showing that the same homodimeric Zur is bound to 22-bp truncated DNA as shown in (b) with labeling positions colored as light pink crosses for Dimer 1-2. The Cy3−Cy5 anchor-to-anchor distances for each labeling position are (B) 56 Å and (D) 34 Å, and the corresponding *E_FRET_* values are ∼0.45 and ∼0.94, respectively. **d,** Crystal structure of two holo Zur dimer bound to DNA, where two dimers of Zur are colored purple and blue, respectively. Cy5 labeling positions (C113) are shown in pink spheres. Cy3−Cy5 anchor-to-anchor distances for Dimer 2 are indicated for the two monomers as (E) 53 Å and (F) 68 Å, and the corresponding *E_FRET_* values are ∼0.53 and ∼0.20, respectively. **e-h,** Cartoons showing DNA-bound two 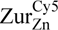 dimers, each at one of the two dyads in two labeling orientations. See details in Supplementary Information 6.2.

**Extended Data Fig. 5.**
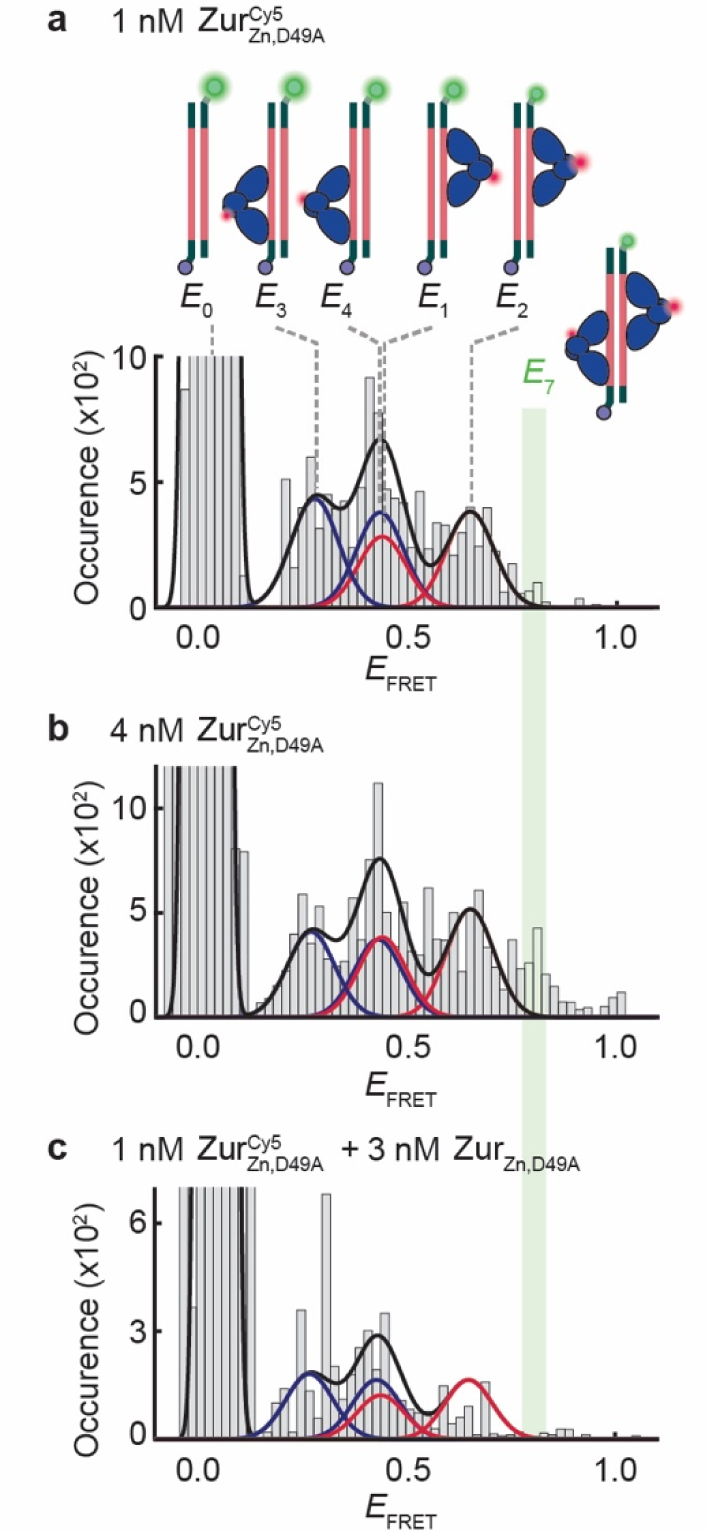
A *E*_FRET_ peak for two 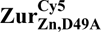 dimers bound to the 31-bp DNA (*E*_7_ at ∼ 0.8) disappears when 75% of the 4 nM 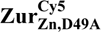 are swapped out to its unlabeled form, supporting that that this *E*_7_ states results from a two-dimer-bound state on DNA. **a,** Histogram of *E*_FRET_ trajectories of immobilized 31-bp DNA^Cy3^ interacting with 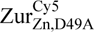 (1 nM). Each color corresponds to a 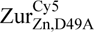 dimer at one of the two dyads of Zur binding box in two orientations (Red: *E*_1_ and *E*_2_ states at the proximal dyad site; blue: *E*_3_ and *E*_4_ states at the distal dyad site). Cartoon shows DNA-bound 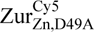 at two binding sites in two orientations on DNA. Salt-bridge mutation (D49A) eliminates key inter-dimer interactions. b, Same as (a), but with 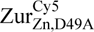 at 4 nM. *E*_7_ state is shaded in green, which only appears when two 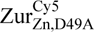 dimers are bound to the 31-bp DNA. c, Same as (b), but 75% of 4 nM 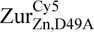 is swapped out into its unlabeled form, Zur_Zn,D49A_; here the *E*_7_ peak disappeared, supporting it was originally from two 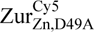 dimer bound state. (a) and (b) are the same figures as Fig. 4a and b in the main text.

**Extended Data Fig. 6.**
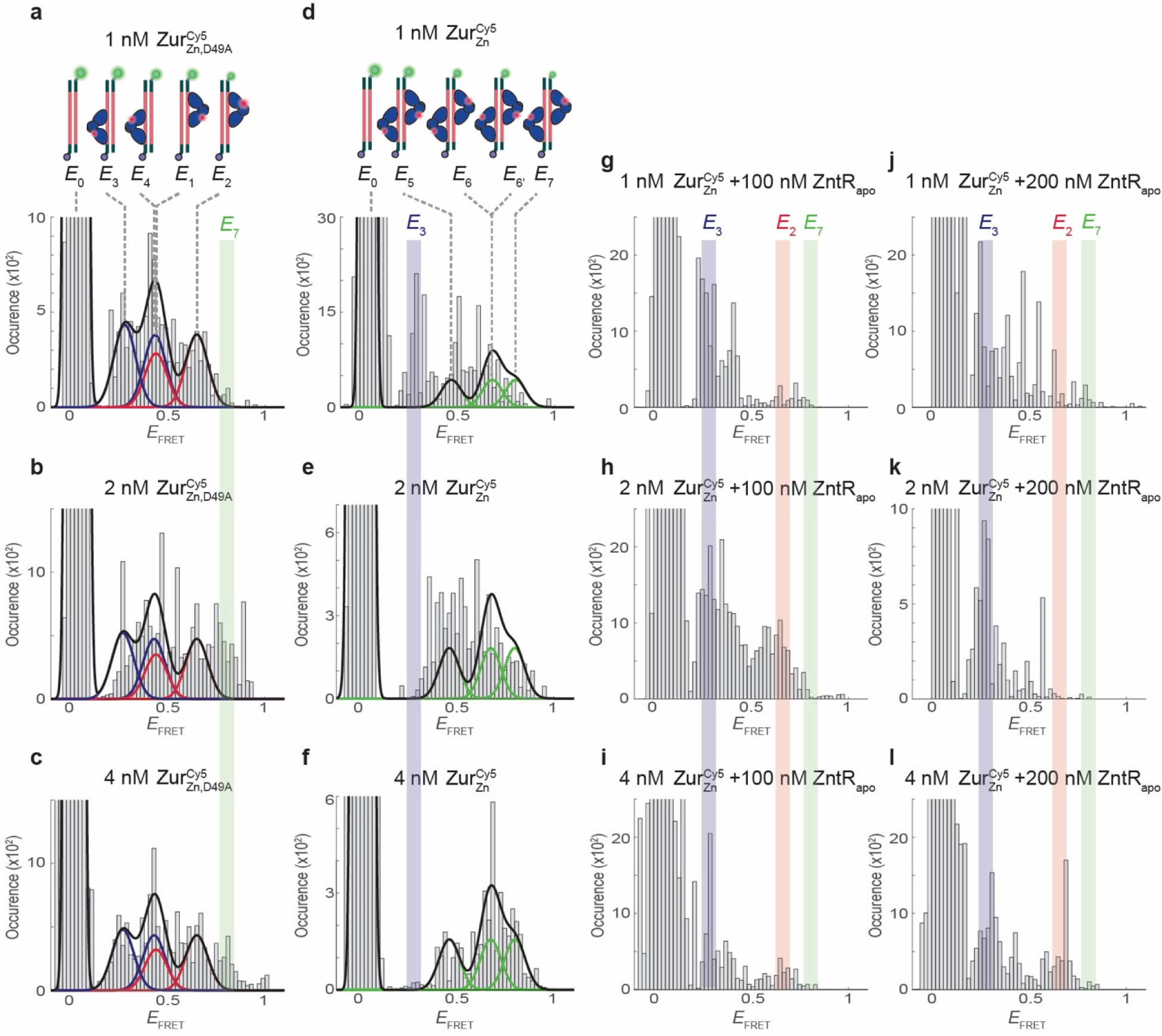
ZntR_apo_ preferentially disrupts Zur_Zn_ binding at the dyad proximal to the Cy3 labeling position on DNA (Supplementary Information 8). Histograms of *E*_FRET_ trajectories of immobilized 31-bp DNA^Cy3^ interacting with different Cy5-labeled Zur constructs in the absence or presence of ZntR_apo_. **a-c,** *E*_FRET_ histograms of 1 to 4 nM 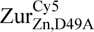 interacting with 31-bp DNA^Cy3^. Blue/red lines are Gaussian-resolved protein-bound states; each color corresponds to the two orientations of one 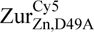 dimer at one of the two dyads of Zur binding box (red: *E*_1_ and *E*_2_ states at ∼0.44 and ∼0.65, respectively; blue: *E*_3_ and *E*_4_ states at ∼0.27 and 0.43, respectively, as assigned in Fig. 4a in the main text). Black line: overall fits. **d-f,** *E*_FRET_ histograms of 1 to 4 nM 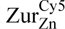 interacting with the 31-bp DNA^Cy3^ (green: *E*_5_, *E*_6_ (*E*_6’_), *E*_7_ states at ∼0.47, ∼0.68, ∼0.80, respectively, as assigned in Fig. 4c in the main text). **g-l,** Same as (d-f), but in the presence of 100 nM (g-i) and 200 nM (j-l) ZntR_apo_. *E*_7_ state position (∼0.8) is denoted as a green shade; it is an indicator of the two-dimer-bound form of Zur on DNA. *E*_3_ state position (∼0.3) is denoted as a blue shade; it is an indicator of the one-dimer-bound form of Zur binding at the dyad distal to the Cy3 labeling position on DNA. *E*_2_ state position (∼0.7) is denoted as a red shade; it is an indicator of one-dimer-bound form of Zur binding at the dyad proximal to the Cy3 labeling position on DNA. Here panel a, c, f, l are the same figures as Fig. 4a, b, c, d in the main text, respectively. See details in Supplementary Information 8.

**Extended Data Fig. 7.**
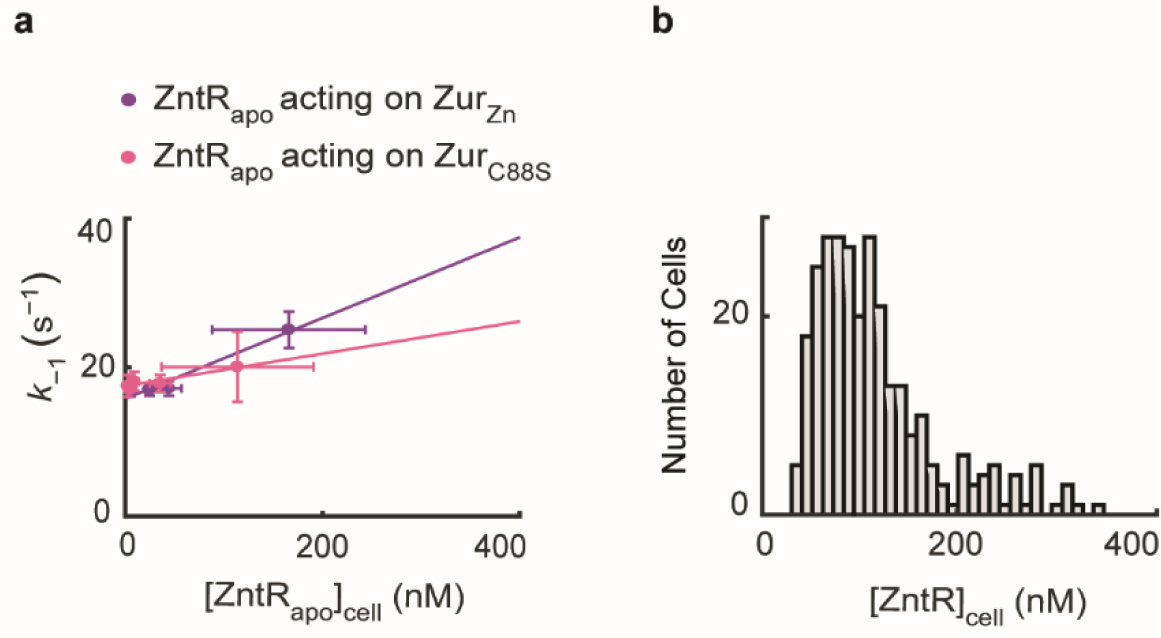
Enhancement of Zur’s apparent unbinding by ZntR_*apo*_ under physiological concentration ranges (Supplementary Information 10). **a**, Dependence of the apparent unbinding rate constant *k*_-_ of Zur_Zn_ (purple) and Zur_C88S_ (magenta) on [ZntR_apo_] in the cell. **b**, Physiological [ZntR] distribution in the cell, obtained by imaging a strain where the ZntR was tagged by mE at the chromosomal locus (Supplementary Table 3).

**Extended Data Table 1.** Summary of extracted kinetic and thermodynamic parameters from live cell imaging experiments.

| | $[\text{ZntR}_{\text{apo}}](\text{nM})$ | $k_0^{\text{off}} (\text{s}^{-1})$ | $k_f (\text{nM}^{-1} \text{s}^{-1})$ | $k_r (\text{s}^{-1})$ | $K_m (\text{nM})$ | $k_1 (\text{nM}^{-1} \text{s}^{-1})^*$ | $K_{D1} (\text{nM})^*$ | $[\text{P}]_{\text{FD}}^{\text{min}} (\text{nM})$ |
| --- | --- | --- | --- | --- | --- | --- | --- | --- |
| Zur<sub>Zn</sub> | 0±0 | 16±1 | 0.016±0.004 |  |  | 0.85±0.16 | 19±4 |  |
|  | 27±8 | 15±1 | 0.019±0.002 | n.o.† | n.o.† | 0.24±0.02 | 62±6 | n.o.† |
|  | 114±15 | 12±2 | 0.041±0.007 |  |  | 0.06±0.01 | 207±58 |  |
|  | 316±19 | 15±3 | 0.059±0.011 |  |  | 0.07±0.38 | 229±1287 |  |
| | | | $k_{\text{bl}} (\text{s}^{-1})$ | $k_{-2} (\text{s}^{-1})$ | $K_{D2} (\text{nM})^*$ | $K_{D3}^*$ | $[\text{D}_0]_{\text{TB}} (\text{nM})^*$ | $[\text{D}_0]_{\text{NB}} (\text{nM})^*$ |
| ZntR<sub>apo</sub> acting on Zur<sub>Zn</sub> |  |  | 225±3 | 7±0.4 | 550±210 | 0.043±0.003 | 966±124 | 1604±79 |
| | $[\text{ZntR}_{\text{apo}}](\text{nM})$ | $k_0^{\text{off}} (\text{s}^{-1})$ | $k_f (\text{nM}^{-1} \text{s}^{-1})$ | $k_r (\text{s}^{-1})$ | $K_m (\text{nM})$ | $k_1 (\text{nM}^{-1} \text{s}^{-1})^*$ | $K_{D1} (\text{nM})^*$ | $[\text{P}]_{\text{FD}}^{\text{min}} (\text{nM})$ |
| Zur<sub>C88S</sub> | 10±8 | 44±64 | 0.035±0.023 | 29±64 | 17±34 | 1.69±0.49 | 26±38 | 66±147 |
|  | 70±30 | 49±46 | 0.047±0.013 | 37±46 | 25±40 | 1.92±0.41 | 25±25 | 86±148 |
|  | 247±72 | 81±107 | 0.068±0.037 | 75±106 | 50±66 | 0.56±0.24 | 146±203 | 155±219 |
| | | | $k_{\text{bl}} (\text{s}^{-1})$ | $k_{-2} (\text{s}^{-1})$ | $K_{D2} (\text{nM})^*$ | $K_{D3}^*$ | $[\text{D}_0]_{\text{TB}} (\text{nM})^*$ | $[\text{D}_0]_{\text{NB}} (\text{nM})^*$ |
| ZntR<sub>apo</sub> acting on Zur<sub>C88S</sub> |  |  | 227±3 | 9.5±0.1 | 3644±638 | 0.003±0.001 | 391±113 | 10234±1010 |
| | $[\text{ZntR}_{\text{Zn}}] (\text{nM})$ | $k_0^{\text{off}} (\text{s}^{-1})$ | $k_f (\text{nM}^{-1} \text{s}^{-1})$ | $k_r (\text{s}^{-1})$ | $K_m (\text{nM})$ | $k_1 (\text{nM}^{-1} \text{s}^{-1})^*$ | $K_{D1} (\text{nM})^*$ | $[\text{P}]_{\text{FD}}^{\text{min}} (\text{nM})$ |
| Zur<sub>Zn</sub> | 27±7 | 2±3 | 0.025±0.009 |  |  | 0.03±0.02 | 76±96 |  |
|  | 64±3 | 7±3 | 0.029±0.011 | n.o.† | n.o.† | 0.44±0.05 | 16±6 | n.o.† |
|  | 131±5 | 7±1 | 0.029±0.007 |  |  | 0.25±0.03 | 29±6 |  |
|  | 407±48 | 9±3 | 0.024±0.009 |  |  | 0.21±0.02 | 41±13 |  |
| | | | $k_{\text{bl}} (\text{s}^{-1})$ | $k_{-2} (\text{s}^{-1})$ | $K_{D2} (\text{nM})^*$ | $K_{D3}^*$ | $[\text{D}_0]_{\text{TB}} (\text{nM})^*$ | $[\text{D}_0]_{\text{NB}} (\text{nM})^*$ |
| ZntR<sub>Zn</sub> acting on Zur<sub>Zn</sub> |  |  | 229±3 | 5.1±0.3 | 1747±1397 | 0.009±0.001 | 1175±216 | 3799±82 |

\*n<sub>0</sub> = 5 (in the fitting routine) †not observed

**Extended Data Table 2.**
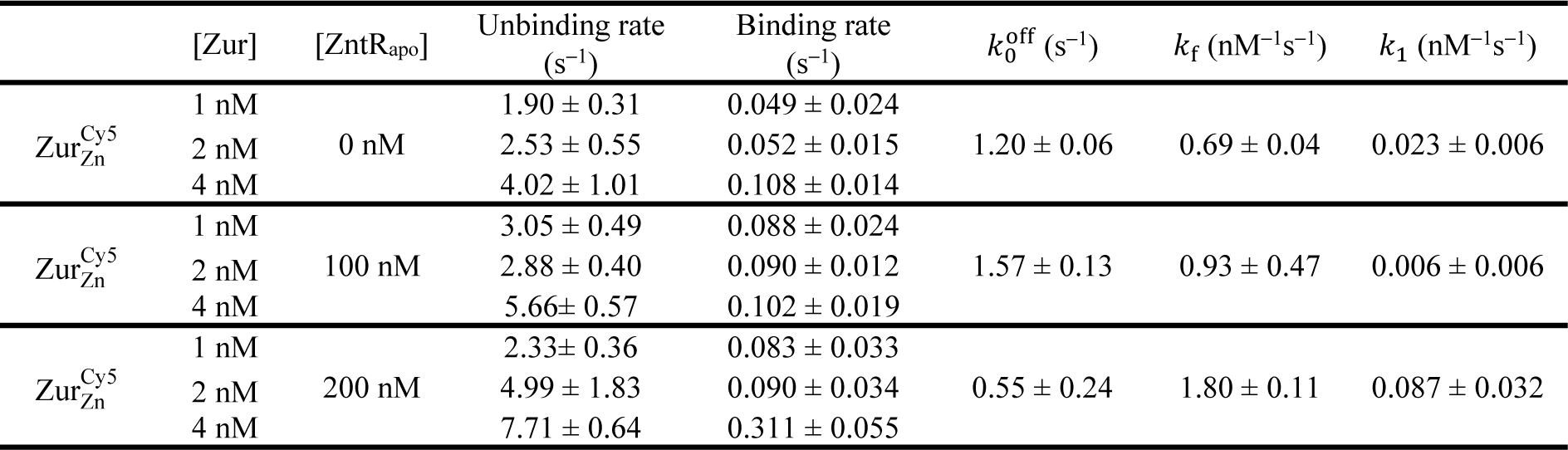
Kinetic parameters for Zur^Cy^^5^–31-bp DNA^Cy3^ interaction measured using *in vitro* smFRET.

