## Supplementary materials for "A ‘through-DNA’ mechanism for metal uptake-vs.-efflux regulation"

**Table of Contents**


**1 Materials and methods ... 3**

1.1 Construction of strains and plasmids for live cell studies ... 3

1.2 Live cell imaging sample preparation, method and data processing for single-molecule imaging, tracking, and protein quantification experimental procedure ... 6

1.3 Construction of strains, protein purification, DNA labeling, sample preparation, imaging and data analysis for in vitro smFRET studies ... 11

**2 Genome sequence analysis and identification of potential recognition sequences of metal efflux regulators (e.g., ZntR) at promoters that are regulated by metal uptake regulators (e.g., Zur), or vice versa, in E. coli, other bacteria, and yeast ... 14**

2.1 Potential partial ZntR recognition sequences around known Zur boxes in E. coli and other bacteria 14

2.2 Aside from the Zur-ZntR Zn uptake-efflux regulator pair, potential Zn efflux regulator recognition sequences are also found at promoters controlled by Zn uptake regulators of other families in bacteria. ... 14

2.3 This pattern of potential existence of partial efflux regulator recognition sequences around known uptake regulator binding box was also observed for regulator pairs involved in the homeostasis of other metals beyond Zn (for example: Fe and Ni) in bacteria. ... 15

2.4 Oppositely, uptake regulator recognition sequence is also found around known efflux regulator binding box in bacteria. ... 15

2.5 In yeast, a similar pattern of efflux regulator recognition sequence overlapping with known uptake regulator binding motifs can be found for iron homeostasis. ... 15

**3 Functionality and intactness of sfGFP-tagged ZntR in E. coli cells ... 15**

3.1 Western blot shows the intactness of ZntR-sfGFP fusion proteins ... 15

3.2 Ensemble fluorescence measurements show that the sfGFP-tagged ZntR is fluorescent ... 17

**4 Analysis of resolvable diffusion states of Zur in the cell and extraction of their effective diffusion coefficients (D) and fractional populations (A) ... 17**

4.1 The resolved three diffusion states of Zur in the cell were assigned as FD, NB, or TB based on their diffusion coefficients and rationales ... 17

4.2 The fractional populations of Zur's three diffusion states show expected dependence on cellular [Zur] ... 22

4.3 Bootstrap analysis shows statistical reliability of data ... 22

**5 Extraction of kinetic and thermodynamic parameters for Zur-DNA interactions in cells ... 23**

5.1 Extraction of $k_{-1}$, the apparent unbinding rate constant from the tight binding state ... 23

5.2 Extraction and summary of additional kinetics and thermodynamic parameters. ... 25

**6 Protein labeling design for single-molecule FRET measurements in vitro ... 26**

6.1 Selecting locations for fluorescent probe location on Zur based on Zur structure, and mutations to make ZntR permanent apo ... 26

6.2 Prediction of $E_{FRET}$ values based on Zur-DNA complex structure ... 27

**7 Procedures for Gaussian fitting to extract $E_{FRET}$ values from the $E_{FRET}$ histograms ... 28**

**8 $ZntR_{apo}$ preferentially disrupts $Zur_{Zn}$ binding at the dyad proximal to the Cy3 labeling position on DNA ... 30**

**9 A through-DNA mechanism for Zur-DNA-$ZntR_{apo}$ interactions and kinetic derivations ... 31**

9.1 Empirical kinetic equation for $ZntR_{apo}$-induced enhancement of Zur's facilitated unbinding and diminishment of Zur's impeded unbinding ... 32

9.2 Kinetic derivation and justification of the mechanistic model for $ZntR_{apo}$-dependent Zur unbinding from DNA ... 33

**10 Within the physiological concentration range of Zur and ZntR, ZntRapo can enhance the apparent unbinding rate constant of $Zur_{Zn}$ by ~130% and that of $Zur_{C88S}$ by ~50% ... 35**

**11 Additional data and figures ... 36**

**12 Supplementary References ... 41**

### 1 Materials and methods

#### 1.1 *Construction of strains and plasmids for live cell studies*

For all cloning and gene editing, the PCRs were performed using the AccuPrime Pfx DNA Polymerase Kit. The primers and enzymes were purchased from the Integrated DNA Technologies and New England Biolabs, respectively. PCR amplifications and digestion products were recovered using the Wizard SV Gel and PCR Clean-Up System (Promega). Plasmid extractions were performed using the QIAprep Spin Miniprep Kit (Qiagen). All primers, plasmids, and strains used are listed in Supplementary Table 1, Supplementary Table ***2*** and Supplementary Table ***3*** respectively.

##### 1.1.1 *Making electrocompetent cells for plasmid transformation or linear DNA homologous recombination*

Transformation of plasmids and linear DNA inserts into *Escherichia coli* BW25113 (CGSC# 7739 Keio Collection, Yale; genotype: (F-*Δ(araD-araB)567, ΔlacZ4787*(::rrnB-3), *λ-*, *rph-1*, *Δ(rhaD-rhaB)568*, *hsdR514*) cells was performed via electroporation. Electrocompetent *E. coli* cells were prepared in the SOB media [2% w/v Bacto Tryptone (Sigma-Aldrich, cat. #: T9410), 0.5 % w/v Bacto Yeast Extract (Sigma-Aldrich, cat. #:Y1625), 10 mM NaCl (Macron, 7581-12), 2.5 mM KCl (Fisher Scientific, P217-500), 10 mM $MgCl_2$ (Mallinckrodt, 5958-04), and 10 mM $MgSO_4$ (Fisher Scientific, M63-500) in nanopure sterile water] containing appropriate antibiotics [ampicillin (100 μg/mL), chloramphenicol (25 μg/mL), or kanamycin (30 μg/mL); USBiological]. In case of homologous recombination, an ampicillin resistant and temperature sensitive pSLTS plasmid was also introduced in the *E. coli* cells. 20 mM L-arabinose (SigmaAldrich, cat. #: A3256), which is a reagent that can induces the expression of the *bet*, *gam*, and *exo* λ-Red enzymes encoded in pSLTS for DNA homologous recombination[1], was used for culturing. The cells were centrifuged and washed twice with cold 10% glycerol (Macron, 5092-02) in nanopure water. The linear DNA inserts or plasmids were then electroporated (2.5 kV or 1.8 kV, using MicroPulser Electroporator; cat.#: 1652100, Bio-Rad) into the prepared electrocompetent cells, and then recovered in SOC medium [SOB medium + 20 mM glucose (Sigma-Aldrich, cat. #: G7528)]. After 4 hours incubation, the cells were plated onto LB-agar containing appropriate antibiotics and further incubated for 18 hours.

Chromosomal DNA insertions and plasmid transformations were verified by colony PCR screening using the Econo Taq DNA Polymerase Kit (Lucigen) and gene sequencing. The temperature sensitive pSLTS plasmid was removed by incubation at 42 °C for 18 hours after successful homologous recombinations and verified by ampicillin selection.

##### 1.1.2 *Construction of ΔzntR and $zur^{mE}$ΔzntR strains*

λ-Red homologous recombination was used to derive the Δ*zntR* (DZR; Supplementary Table 3) and *$zur^{mE}$*Δ*zntR* ($Zur^{mE}$-DZR; Supplementary Table 3) strains from *Escherichia coli* BW25113 and *$zur^{mE}$* strains respectively [2]. A linear DNA insert targeting the *zntR* gene in the chromosome was made using primers H1H2DZntrPT2SK-fp and DZntrH1H2PT2SK-rp (Supplementary Table 1) together with a template containing a kanamycin resistance gene cassette containing an I-SecI recognition site, for subsequent RecA recombination, obtained from the pT2SK plasmid[1]. The linear insert was introduced via electroporation into the electrocompetent BW25113 and *$zur^{mE}$* strains bearing a temperature sensitive pSLTS plasmid. The cells were recovered in 1 mL SOC medium, incubated at 30 °C and shaking at 250 rpm for 4 hours, and finally plated onto LB-agar plate containing both ampicillin (50 μg/mL) and kanamycin (15 μg/mL) for Δ*zntR*, and ampicillin (10 μg/mL), chloramphenicol (10 μg/mL) and kanamycin (15 μg/mL) for *$zur^{mE}$*Δ*zntR*, resulting in the strains DZR and $Zur^{mE}$-DZR (Supplementary Table 3). Deletions were further confirmed by colony PCR.

##### 1.1.3 *Construction of ΔzurΔzntR double deletion strain*

The Δ*zur*Δ*zntR* strain was derived from the DZR strain (Supplementary Table 3). First the kanamycin resistance cassette at the erstwhile *zntR* locus in DZR was eliminated via RecA recombination.

To induce I-SceI enzyme cleavage mediated scar-less elimination of the kanamycin resistance cassette, a sample of DZR overnight culture was diluted 1:50 in 10x PBS buffer; 200 µL was plated onto LB-agar plate containing anhydrotetracycline (aTc) (150 ng/mL; Acros Organics). To confirm the elimination of the kanamycin resistance cassette in the genome, 8 colonies from the aTc plate were tested for kanamycin sensitivity on LB-agar plate containing kanamycin (30 µg/mL). Cells from colonies that had kanamycin-sensitive phenotypes were chosen for DNA sequencing to confirm the presence of the desired genomic edit. A linear DNA insert targeting the *zur* gene in the chromosome was made using primers H1H2DZurPT2SK-fp and DZurH1H2PT2SK-rp (Supplementary Table 1) together with a template containing a kanamycin resistant cassette. Subsequent steps for homologous recombination and deletion of the *zur* gene were followed according to procedures described above. The pSLTS plasmid was removed from the strains by culturing the cells at 42 °C overnight. The strain thus obtained, DZ-DZR (Supplementary Table 3), lacked both *zur* and *zntR* genes.

##### 1.1.4 *Construction of* $zntR^G_{C115S}$ *and* $zntR^G$ *in L-arabinose inducible pBAD plasmids*

To spectrally separate Zur and ZntR in the cells, we tagged *zntR* and $zntR_{C115S}$ with super-folder GFP. To make the pBAD33 (chloramphenicol resistant) plasmid expressing $zntR_{C115S}$-*sfGFP*, the $zntR_{C115S}$ gene was first cloned out of the plasmid pBZR(C115S)-mEos3.2[3] using primers Sac1-EZntR-pB33-fp and ZntR_GFP_rp. The *sfGFP* gene was cloned from the sfGFP-pBAD plasmid[4] using primers GFP_fp and sf-GFP-rp. We used overlapping PCR with primer pairs Sac1-EZntR-pB33-fp and sf-GFP-rp-pst1 to tag the $zntR_{C115S}$ gene with *sfGFP*. After PCR amplification using AccuprimePfx DNA Polymerase, the linear $zntR_{C115S}$-*sfGFP* product was digested with SacI-HF and PstI-HF restriction enzymes and inserted into a similarly digested pBAD33 plasmid using quick ligase enzyme to generate the p33ZRG(C115S) plasmid. Another plasmid p24ZRG(C115S) was constructed using the pBAD24 vector backbone bearing the same gene insert for differential antibiotic selections. Next the *zntR* gene was copied out of the pBZntR-mEos3.2 plasmid[3] using primers Sac1-EZntR-pB33-fp and ZntR_GFP_rp. The linear $zntR$-*sfGFP* product was again obtained by overlapping PCR using primers zntRsfGFP_Sac1_Gib_fp and zntRsfGFP_Sac1_Gib_rp and was digested with SacI-HF enzyme and inserted into a digested pBAD33 plasmid using Gibson Assembly Mastermix (New England Biolabs) to generate the p33ZRG plasmid. The plasmids p24ZRG(C115S), p33ZRG(C115S) and p33ZRG (Supplementary Table 2) were then each transformed into *E. cloni 10G* chemically competent cells for propagation and miniprep. The constructs were subsequently confirmed by colony PCR and DNA sequencing.

Another version of the $Zur^{mE}$-DZR strain, where the *zur-mEos3.2* gene was encoded in a plasmid rather than in the chromosome, was also constructed, DZ-DZR-pZmE, via the electroporation of the pZur_mE plasmid into the DZ-DZR strain. Electroporation of the plasmid pApoZur_mE in the DZ-DZR strain led to the construction of a DZ-DZR-pZmEC88S strain. Subsequently, the p33ZRG(C115S) plasmid was transformed into the DZ-DZR, DZ-DZR-pZmE and DZ-DZR-pZmEC88S strains resulting in the DZ-DZR-pZRGC115S, DZ-DZR-pZmE-pZRGC115S and DZ-DZR-pZmEC88S-pZRGC115S strains. The p33ZRG plasmid was transformed into DZ-DZR-pZmE, resulting in the DZ-DZR-pZmE-ZRG strain. The p24ZRG(C115S) plasmid was transformed into DZR-$Zur^{mE}$, resulting in the DZR-$Zur^{mE}$-pZRGC115S strain (Supplementary Table 3).

**Supplementary Table 1 |** List of primers used in this study**.**

| Primer Name | Sequence (5′-3′) |
|---|---|
| 1. Sac1 EZntR-pB33-fp | AATTCGAGCTCAGGAGGAATTCACCATGTATCGCATTGGTGAGCT |
| 2. Pst1 EZntR-pB33-rp | TGCCTGCAGTTATTTATCATCATCATCTTTATAATCAGGACGACAACCACTC TTAACGCC |
| 3. EcoR1 – EzntR-fp | GGA GGAATT CACCATGTATCGCATTGGTGAGCT |
| 4. ZntR_GFP_rp | TCCTCGCCCTTGCTCACCATACAACCACTCTTAACGCCAC |
| 5. GFP_fp | ATGGTGAGCAAGGGCGAGGA |
| 6. sf-GFP-rp | CTTGTACAGCTCGTCCATGCC |

| | |
|---|---|
| 7. sf-GFP-rp-pst1 | GCATGCCTGCAGTTACTTGTACAGCTCGTCCA |
| 8. zntRsfGFP_Sac1_Gib_fp | TGGGCTAGCGAATTCGAGCTAGGAGGAATTCACCATGTATC |
| 9. zntRsfGFP_ Sac1_Gib_rp | GGATCCCCGGGTACCGAGCTTTACTTGTACAGCTCGTC |
| 10. H1H2DZntrPT2SK-fp | ATCAACGATAACTAGTGGAGTATGTTTTTTTGCACTGGCAATCTCAAGAGT GGCAGC |
| 11. DZntrH1H2PT2SK- rp | AGTGTAATCCTGCCAGTGCAAAAAAACATACTCCACTAGTTTACGCCCCGC CCTGC |
| 12. H1H2DZurPT2SK- fp | CTTAACCCCCACTTTGAGGTGCCCGGAGGGCGTACATCCTATCTCAAGAGT GGCAGC |
| 13. DZurH1H2PT2SK- rp | GACGTGTACAAGGATGTACGCCCTCCGGGCACCTCAAAGTTTACGCCCCGC CCTGC |
| 14. znuC220_up | CAGAAGCTGTATCTCGACACC |
| 15. znuC297_dn | TTCTTTATGTGTACCAGGGCG |
| 16. pET_T7_fp | TACGACTCACTATAGGGG |
| 17. pET_down_rp | CCAAGGGGTTATGCTAGT |
| 18. C17S_fd | GCAGGCTGAAAAAATCAGCGCGCAGCGTAATGTGC |
| 19. C17S_rc | GCACATTACGCTGCGCGCTGATTTTTTCAGCCTGC |
| 20. C152S_rc | ACTGTTCAGGATGACGACTCGCTTCCACTTCTACA |
| 21. C152S_fd | TGTAGAAGTGGAAGCGAGTCGTCATCCTGAACAGT |
| 22. C113S_fd | CGCAGTGAAAGAAGAGAGTGCAGAAGGCGTGGAAG |
| 23. C113S_rc | CTTCCACGCCTTCTGCACTCTCTTCTTTCACTGCG |
| 24. C158S_fd | TCGTCATCCTGAACAGAGCCAGCATGATCACTCTG |
| 25. C158S_rc | CAGAGTGATCATGCTGGCTCTGTTCAGGATGACGA |
| 26. EZurD49A-fp | ATGATCTGCTTGCTTTACTGCGCG |
| 27. EZurD49A-rp | CGCGCAGTAAAGCAAGCAGATCAT |
| 28. NdeI EZntR_fp_pET3a(5) | ATATACATATGTATCGCATTGGTGAGCTGGC |
| 29. BamHI EZntR_rp_pET3a(5) | CAGCCGGATCCTTATTTATCATCATCATCTTTATAATCAGGACGACAACCA CTCTTAACG |

**Supplementary Table 2 |** List of plasmids used or constructed in this study.

| Plasmid Name | Gene Insert | Resistance | Source |
|---|---|---|---|
| 1. pSLTS | bet, gam, exo recombinase enzymes, ISceI enzyme | Amp | [1] (Addgene plasmid 59386) |
| 2. pT2SK | kanamycin cassette, I-SecI cleavage site | Kan | [1] (Addgene plasmid 59383) |
| 3. sfGFP-pBAD | Superfolder Green fluorescent protein | Amp | [4] |
| 4. pBAD24 | L-arabinose inducible, Base Plasmid | Amp | [5] |
| 5. pBAD33 | L-arabinose inducible, Base Plasmid | Cam | [5] |
| 6. pBZntR-mEos3.2 | zntR-mEos3.2-FLAG | Amp | [3] |
| 7. pBZR(C115S)-mEos3.2 | zntR-C115S-mEos3.2-FLAG | Amp | [3] |
| 8. pApoZur_mE | zur-C88S- mEos3.2-FLAG | Amp | [2] |
| 9. pZur_mE | zur-mEos3.2-FLAG | Amp | [2] |
| 10. p24ZRG(C115S) | zntR-C115S-sfGFP | Amp | This Study |
| 11. p33ZRG(C115S) | zntR-C115S-sfGFP | Cam | This Study |
| 12. p33ZRG | zntR-sfGFP | Cam | This Study |
| 13. pET3a | T7 (IPTG inducible) | Amp | Novagen |
| 14. pZntRapo | ZntR(C115S) | Amp | [3] |
| 15. pZurC113 | Zur (C17S, C152S, C158S) | Amp | This Study |
| 16. pZurC113D49A | Zur (C17S, C152S, C158S, D49A) | Amp | This Study |
| 17. pZurC158 | Zur (C17S, C113S, C152S) | Amp | This Study |

**Supplementary Table 3 |** List of strains constructed in this study.

| Strains | Plasmids | Chromosomal modification | Source |
|---|---|---|---|
| 1. BW25113 | none | Base Strain | Keio collection |
| 2. ZRM3.2 | none | *zntR-mEos3.2* | [3] |
| 3. DZR | none | *ΔzntR* | This study |
| 4. Zur$^{mE}$-DZR | none | *zur-mEos3.2-FLAG, ΔzntR* | This study |

| | | | |
|---|---|---|---|
| 5. DZR-Zur$^{mE}$-pZRGC115S | p24ZRG(C115S) | *zur-mEos3.2-FLAG, ΔzntR* | This study |
| 6. DZ-DZR | none | *Δzur, ΔzntR* | This study |
| 7. DZ-DZR-pZmE | pZur_mE | *Δzur, ΔzntR* | This study |
| 8. DZ-DZR-pZmEC88S | pApoZur_mE | *Δzur, ΔzntR* | This study |
| 9. DZ-DZR-pZRGC115S | p33ZRG(C115S) | *Δzur, ΔzntR* | This study |
| 10. DZ-DZR-pZmE-pZRGC115S | pZur_mE, p33ZRG(C115S) | *Δzur, ΔzntR* | This study |
| 11. DZ-DZR-pZmEC88S-pZRGC115S | pApoZur_mE, p33ZRG(C115S) | *Δzur, ΔzntR* | This study |
| 12. DZ-DZR-pZmE-ZRG | pZur_mE, p33ZRG | *Δzur, ΔzntR* | This study |

**Supplementary Table 4 |** Abbreviations used in this study.

| **Abbreviation** | **Full form** |
|---|---|
| SMT | Single-molecule tracking |
| SCQPC | Single cell quantification of protein concentration |
| WT | Wild Type |
| Zur | Zinc Uptake Regulator |
| ZntR | Zinc Transport Regulator |
| mE | mEos 3.2 protein |
| sfGFP or G | Super-folder Green Fluorescent Protein |
| Amp | Ampicillin |
| Kan | Kanamycin |
| Cam | Chloramphenicol |
| PDF | Probability Distribution Function |
| CDF | Cumulative Distribution Function |
| FD | Freely Diffusing |
| NB | Non-specific Binding |
| TB | Tight Binding |
| PWDD | Pair-wise Distance Distribution |
| iqPALM | Image-base quantitative photo-activated localization microscopy |
| smFRET | Single-molecule Föster Resonance Energy Transfer |
| PALM | Photo-Activated Localization Microscopy |

#### 1.2 *Live cell imaging sample preparation, method and data processing for single-molecule imaging, tracking, and protein quantification experimental procedure*

##### 1.2.1 *Sample preparation for live cell imaging:*

A single *E. coli* cell colony was inoculated into and grown in LB medium for 18 h at 37 °C. This overnight culture was diluted 1:100 in M9 medium[3] supplemented with amino acids (GIBCO, cat. #: 11130051), vitamins (GIBCO, cat. #: 11120052), and 0.4% glycerol, and further grown to OD600 of 0.3. L-arabinose was added to induce plasmid expression for 0 - 20 mins when applicable. For Zn stress, $ZnSO_4$ was added into the media to a final concentration of 20 µM or 100 µM. 2 mL of the cell culture was pelleted via centrifugation and washed thrice with the same M9 media (supplemented with 0.4% glucose instead of glycerol), and was further incubated at 37 °C for 1 hour to help maturation of the fluorescent protein tags. The cells were then collected by centrifugation and added onto an agarose gel pad between a coverslip (Thermo Scientific Cat. #: 20848) pre-dispersed with 100 nm gold nanoparticles (Ted Pella, Inc., Cat. #: 15708-9) and a glass slide (VWR Lot #: 48300-37), and sealed with epoxy-glue.

##### 1.2.2 *Single-molecule tracking (SMT) and single cell quantification of protein concentration (SCQPC):*

SMT via stroboscopic imaging and SCQPC were performed as described previously, using a homebuilt PALM microscope based on Olympus IX71 (Extended Data Fig. 2a)[2,3]. For SMT, a short (20 ms) and low power (1-100 W/cm$^2$) 405-nm laser illumination was used to photoconvert a single mEos3.2 tagged protein from its native green fluorescent form to the red fluorescent form. 30 pulses of a 561-nm laser exposure (21 kW/cm$^2$), with 4 ms pulse duration and time-lapse $T_{tl}$ = 40 ms was used to excite this red fluorescence. The EMCCD exposure was synchronized with the 561-nm laser pulses, and this stroboscopic imaging allowed us to obtain diffraction limited images of both stationary and mobile single molecules. This process was repeated for 500 cycles for each cell to obtain a tracking movie.

After the SMT cycles, we perform the SCQPC part. Here the cells were illuminated with 405-nm laser (100 W/cm$^2$) for 2 mins to photoconvert all the remaining green mEos3.2 to their red form, the emission of which was excited by 561-nm laser illumination at the same power density for 2000 frames to obtain the whole cell fluorescence intensity of mEos3.2 and photobleach them. This step was repeated for a total of 3 cycles to ensure all mEos3.2 tagged proteins were photoconverted, imaged, and photobleached.

After all the fluorescence of the mEos3.2 in the cell was photobleached following the steps above, the total green fluorescence of the remaining sfGFP-tagged-ZntR was excited by a 488-nm laser (7 kW/cm$^2$), for 1000 frames to obtain the whole cell intensity.

##### 1.2.3 *Determination of total cellular Zur and ZntR copy numbers*

To obtain the total Zur copy number $N_{cell}$ in each cell, the whole cell mEos3.2 red emission obtained in the SCQPC step, was divided by the average intensity of a single mEos3.2 molecule in that cell obtained from the SMT steps[2,3]. The total copy number was estimated using the following Eq. S1:

$$N_{\text{cell}} = \frac{N_{\text{SMT}} + N_{\text{SCQPC}}}{\text{PE}_{\text{mEos3.2}} * \text{OS}_{\text{Zur}}} \qquad \text{Eq. S1}$$

where, $N_{SMT}$ and $N_{SCQPC}$, are the copy numbers obtained from the SMT and SCQPC, respectively. $PE_{mEos3.2}$ is the photoconversion efficiency of mEos3.2 protein (=0.42)[6,7] and $OS_{Zur}$ is the oligomerization state of Zur (homodimer, $OS_{Zur}$ = 2).

To determine the single-molecule intensity of mEos3.2 from the SMT step, a custom-written MATLAB software called iQPALM (Image-based Quantitative Photo-Activated Localization Microscopy) (Supplementary Table 4)[3] and Figshare software[8] was used to process the fluorescence images to determine the centroid location of the candidate red single mEos3.2 fluorescence spots. The cell boundary was first determined using the bright field optical transmission image. Furthermore, cells with length of 2.7 ± 0.9 μm were selected to decrease the possibility of picking dividing cells, which potentially contain more than one copy of chromosome (Supplementary Fig. 1, 2$^{nd}$ column). The cell boundaries in the region of interest (ROI) were then superimposed onto the corresponding fluorescence image to select candidates of single-molecule fluorescence within the cell boundaries, which were then determined by fitting the fluorescence spots with a two-dimensional Gaussian point spread function (PSF) in Eq. S2, as was described previously[2,3].

$$I(x,y) = A\exp\left[-\frac{(x-x_o)^2}{2{\sigma_x}^2} - \frac{(y-y_o)^2}{2{\sigma_y}^2}\right] + B \qquad \text{Eq. S2}$$

where, $I(x,y)$ is the fluorescence intensity at position $(x,y)$, and $A$, $B$, $(x_o, y_o)$, and $(\sigma_x, \sigma_y)$ are the amplitude, background, centroid position, and standard deviations of the Gaussian function.

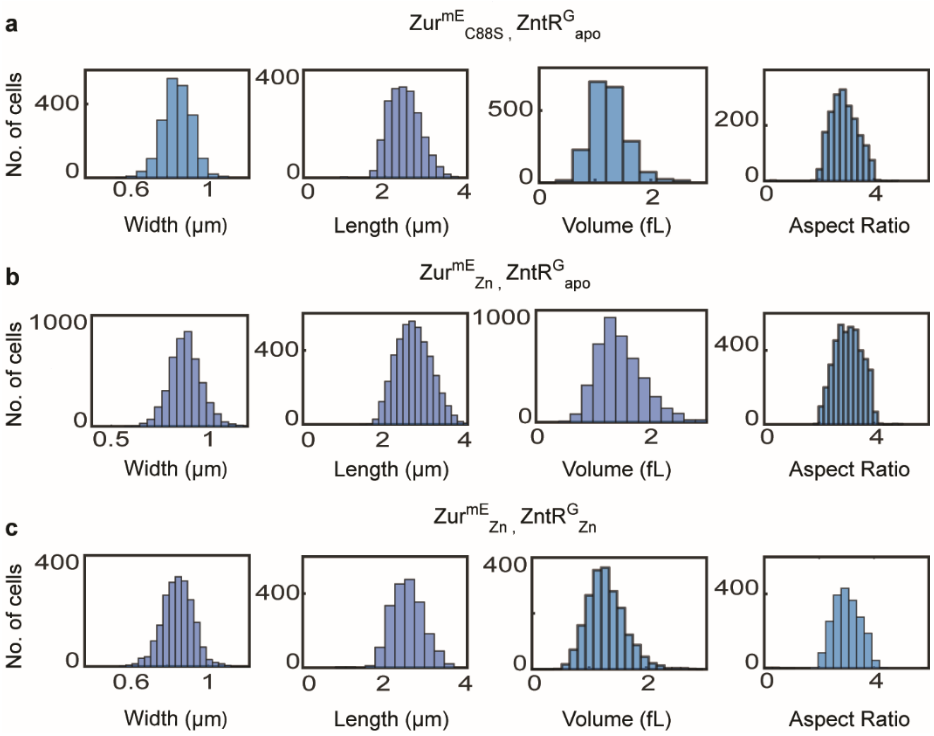


**Supplementary Fig. 1 | Exemplary distribution of cell width, length, volume, and aspect ratio. a**, DZ-DZR-pZmEC88S-pZRGC115S strain expressing Zur$^{mE}_{C88S}$ and ZntR$^{G}_{apo}$ (i.e., ZntR$^{G}_{C115S}$ ) in the cells, **b**, DZ-DZR-pZmE-pZRGC115S strain expressing Zur$^{mE}$ and ZntR$^{G}_{apo}$ in the cells, and **c**, DZ-DZR-pZmE-pZRG strain expressing Zur$^{mE}$ and ZntR$^{G}$ in the cell.

To estimate the total ZntR copy number, we need to obtain the single sfGFP intensity. Since sfGFP is not a photoconvertible or photoactivatable fluorescent protein, the single sfGFP intensity was determined separately as described: Firstly, a sample prepared from a strain containing only the ZntR$^{G}_{apo}$ (DZ-DZR-pZRGC115S; Supplementary Table 3) was illuminated using 488-nm laser excitation (2-7 kW/cm$^2$) with 4 ms exposure for 10000 frames. The candidate fitted spots were then filtered by their spot sizes; here the spot size is measured by the standard deviation of the fitted Gaussian PSF (Eq. S2), $\sigma_x$ (or $\sigma_y$), the theoretical value of which is around 110 nm on the basis of diffraction-limited resolution ($\frac{\lambda}{2\text{NA}}$; NA is the objective numerical aperture). Based on the distribution of $\sigma_x$, we rejected any spot with $\sigma_x$ smaller than 50 nm (too narrow for a reasonable single-molecule PSF) and any $\sigma_x$ greater than 350 (too wide for a clean single-molecule PSF) (Supplementary Fig. 2a)[3]. Further, the first 1200 frames of each cell were removed to decrease the contamination by fluorescence images of an ensemble of sfGFP in the cell (Supplementary Fig. 2b), as initially the cell contains many sfGFP-tagged ZntR$_{apo}$. From the remaining frames, the candidate single sfGFP spots were analyzed similarly as described above, to obtain the intensity of the fitted Gaussian function ($I(x,y)$) and the spot sizes described by ($\sigma_x$, $\sigma_y$) in the Gaussian function.

To decouple and determine the local power dependence of the single molecule intensity, experiments were done at three different power densities. Two-dimensional histograms of these filtered $\sigma_x$ and their corresponding intensity ($I$), which is the integrated volume of the fitted Gaussian function component in Eq. S2) resolved three populations (Supplementary Fig. 2c-e). These candidate populations were globally fitted with a three-component bivariate Gaussian function across the three different experimental power densities sharing the width ($\sigma_{x2}$), and peak position ($x_2$), of the 2$^{nd}$ component in Eq. S3 (Supplementary Fig. 2c-e, red dotted line) to further resolve the correct candidate spots and obtain their intensities:

$$z = \sum_{i=1}^{3} A_i e^{\frac{(x-x_i)^2}{2\sigma_{xi}{}^2} - \frac{(I-Ii)^2}{2\sigma_{Ii}{}^2}} \qquad \text{Eq. S3}$$

Eq. S3 comprises 3 components of a bivariate Gaussian function where the first and third components correspond to populations of $\sigma_x$ narrower and far greater than the theoretical value of a single-molecule PSF (~110 nm), respectively. The population of $\sigma_x$ lower than the theoretical value was determined to be false detections as this population was also observed in the wild type BW25113 strain that does not express any fluorescence tag (Supplementary Fig. 2f). The 2nd component in the fitting corresponds to a population of $\sigma_x$ (centered at 137 nm), similar to the expected PSF size, and thus is assigned as single sfGPF molecules in the cell. Due to high cell-to-cell heterogeneity of protein concentration, the total fluorescence of every cell decays to different extents after 1200 frames, at which there are still many cells that contain many sfGFPs and whose fluorescence is from the ensemble fluorescence. We assigned the population of $\sigma_x$ with mean value higher than 137 nm to such any remaining ensemble fluorescence.

The single-molecule population $\sigma_x$ (=137 nm) from the 2nd term (Supplementary Fig. 2c-e, red Gaussian fit) was selected out using thresholds (magenta lines) determined from the fitting (Supplementary Fig. 2c-e). For these selected single-molecule candidate spots, the corresponding intensities and the local power densities were determined (from the centroid position of the cell in the microscope field of view) and a calibration curve was formed. This single-molecule intensity and local power density calibration curve served to determine the corresponding single sfGFP intensity (Supplementary Fig. 2g).

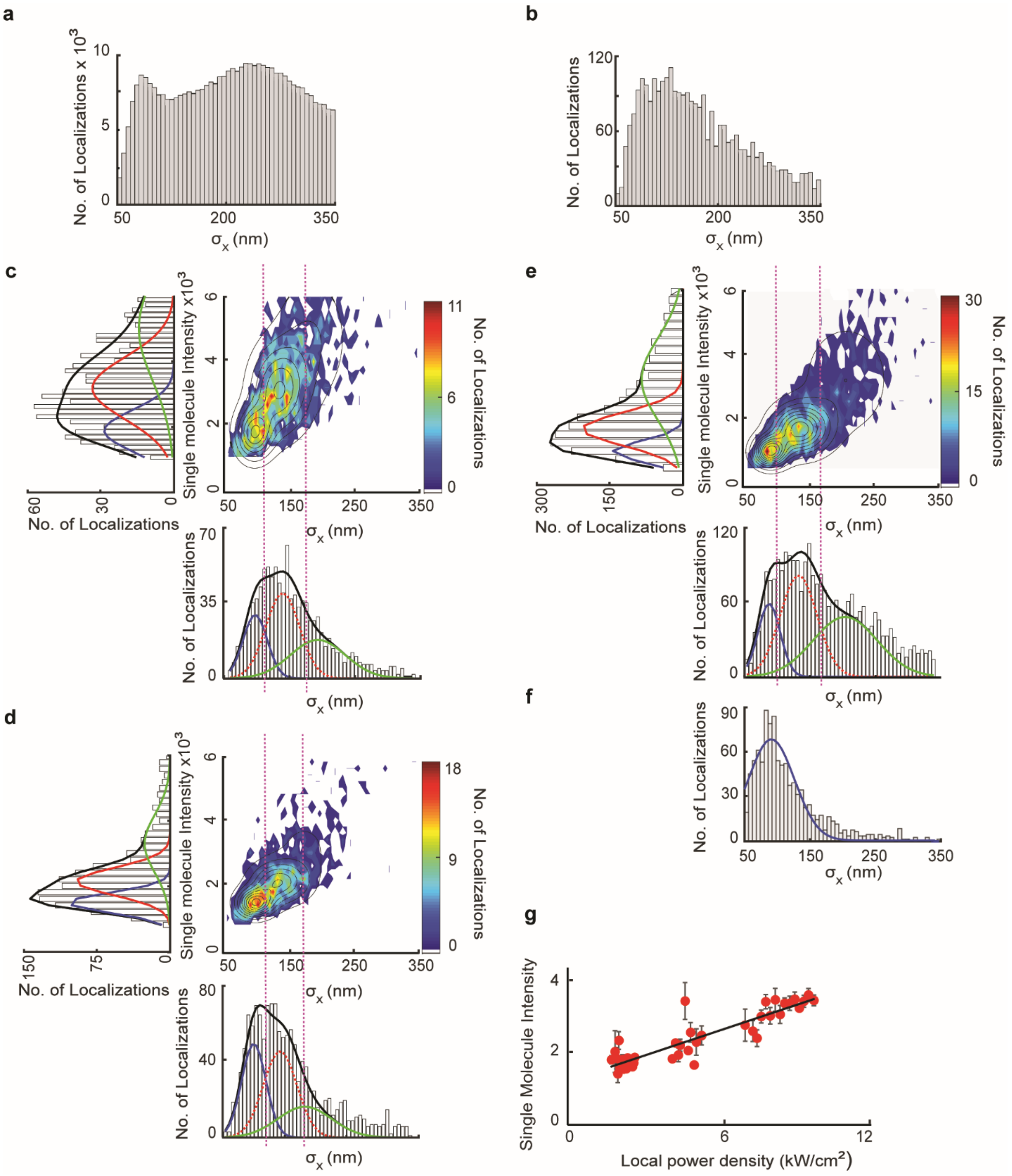


**Supplementary Fig. 2 | Determination of single-molecule intensity of sfGFP. a**, Distribution of filtered candidate fluorescence spot size, measured by the Gaussian function standard deviation σ in Eq. S2, 50 nm < $\sigma_x$ < 350 nm. **b**, Distribution of the candidate spot sizes, $\sigma_x$, after removal of first 1200 frames, resolves a single population. **c-e**, Two-dimensional distribution of the candidate spot sizes and intensities at three laser power densities 7 kW/cm$^2$ (c), 3.5 kW/cm$^2$ (d), and 2 kW/cm$^2$ (e). The corresponding one-dimensional projections of the $\sigma_x$ and intensity are plotted along the x and y axes, respectively. The black line is the total fit. The three colored lines (blue, red, and green) are the three Gaussian components of the fit, corresponding to the false detection, single molecules, and multi-molecule/ensemble populations, respectively. The dotted red line component represents the terms of the fit that were globally shared across the data of the three power densities. The vertical magenta dashed lines represent the thresholds applied to extract and isolate the middle population of correct candidates. **f**, Distribution of $\sigma_x$ from WT BW25113 after similar filtering, shows only one population with average lower than 100 nm (blue population), resulting from false detection in the green channel for sfGFP imaging. **g**, Single molecule intensity vs. local power density calibration curve.

Finally, the whole cell sfGFP intensity vs. time was fitted with a double exponential function and an offset (Supplementary Fig. 3) (Eq. S4):

$$y = Ae^{-k_1 x} + Be^{-k_2 x} + \mathrm{C} \qquad \text{Eq. S4}$$

where, the first component corresponds to the sfGFP fluorescence intensity and the 2$^{nd}$ to the cellular auto-fluorescence, and C is the background offset. Subtraction of the 2$^{nd}$ and 3$^{rd}$ terms of Eq. S4 from the fluorescence decay (Supplementary Fig. 3, blue curve) yields the total cellular sfGFP fluorescence, whose value at frame no. 1 is taken as the initial total cellular sfGFP fluorescence.

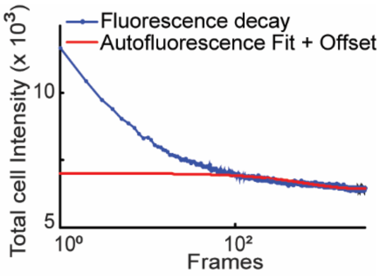


**Supplementary Fig. 3 |** An example of a whole cell sfGFP fluorescence photobleaching decay curve (blue) obtained by imaging a strain that expresses only $\mathrm{ZntR}^{\mathrm{G}}_{\mathrm{apo}}$ (DZ-DZR-pZRGC115S). The autofluorescence and offset are shown in red.

The apparent copy number $N_{\mathrm{SCQPC}}$ of the sfGFP in the cell was calculated by dividing this initial total sfGFP fluorescence intensity with the single sfGFP intensity determined from the calibration curve (Supplementary Fig. 2g), with the local power density calculated from the centroid position of the cell. The total copy number of sfGFP-tagged ZntR in the cell $N_{\mathrm{cell}}$ was estimated using Eq. S5:

$$N_{cell} = \frac{N_{SCQPC}}{OS_{ZntR}} \qquad \text{Eq. S5}$$

where $\mathrm{OS}_{\mathrm{ZntR}}$ (=2) corresponds to the oligomeric state of ZntR (a homodimer).

#### 1.3 *Construction of strains, protein purification, DNA labeling, sample preparation, imaging and data analysis for in vitro smFRET studies*

##### 1.3.1 *Mutagenesis, expression, purification, and fluorescence labeling of Zur variants*

To label Zur with the FRET acceptor Cy5, site-directed mutagenesis was used to make Zur variants that contain a unique labelable cysteine in each monomer (see Supplementary Information 6.1) on the choice of cysteine position, the removal of other natural cysteines, and protection of essential Zn-binding cysteines at structural and regulatory sites). For example, the Zur variant, $\mathrm{Zur}^{\mathrm{Cy5}}$, was created, expressed, purified, labeled and further purified to obtain the single Cy5-labeled form at the specific cysteine at position C113. All Zur variants were cloned in a pET3a vector and the sequence was confirmed. The proteins were expressed in *E. coli* (BL21 DE3) cells and purified as previously described[9]. Briefly, the cells were grown until $OD_{600}$ ~ 0.6 before 0.4 mM isopropyl-beta-D-thiogalactopyranoside (IPTG) and 0.2 mM $ZnSO_4$ were added. After an additional 3 hours growth at 37 °C, cells were harvested by centrifugation and then lysed with lysozyme in lysis buffer (20 mM Tris, 300 mM NaCl, 50 mM Dithiothreitol (DTT), and 10% glycerol at pH 8.0). The cells were further disrupted with three cycles of freeze and thaw followed by sonication. The proteins were collected by centrifugation and the pellet was suspended in denaturing buffer (20 mM Tris, 6 M Urea, 100 mM Dithiothreitol (DTT) at pH 8.0). After 30 min of denaturing at 4 °C, the solution was centrifuged to remove insolubles, and the soluble proteins in the supernatant were collected and then added drop by drop into the refolding buffer (20 mM Tris, 100 μM $ZnSO_4$, 5 mM Dithiothreitol (DTT) at pH 8.0) to refold the protein. Then, the protein was purified by anion exchange column (HiTrap Q HP, GE Healthcare). The collected fractions were further purified through a Heparin affinity column (16/10 Heparin FF, GE Healthcare), a gel filtration column (HILOAD 26/60 Superdex 200 PR, GE Healthcare), and an anion exchange column (Mono Q 5/50 GL, GE Healthcare). Protein purity was confirmed by SDS-PAGE, quantified using UV measurement at 280 nm, and stored at −80 °C in 50 mM Tris buffer with 50 mM NaCl, 10 nM $ZnSO_4$, 2 mM TCEP, and 10% glycerol at pH 8. Protein identity was confirmed by mass spectrometry (ESI-TOF, HPLC-ESI-MS/MS, UT Health San Antonio; Supplementary Fig. 18 later).

Cy5 FRET acceptor was labeled at the targeted cysteine in protein via maleimide chemistry. Zur was present as the fully-metallated holo-dimer form in the presence of $ZnSO_4$ with excess TCEP reducing disulfide bond formation. The $Zn^{2+}$ ion first binds to the metal-binding cysteines of Zur, protecting these cysteines from dye labeling. Cy5-maleimide (Invitrogen) was added to the holo-Zur solution ([dye]:[Zur monomer] = 6:1) in 100 mM phosphate buffer solution at pH 7. The reaction mixture was kept on a shaker at 4 °C for ~18 hours and then quenched by adding excess beta-mercaptoethanol (BME). After incubating for additional 2 hours, the excess dye was removed through gel filtration (Superdex peptide 10/300 GL, GE Healthcare). Because Zur is a homodimer, a mixture of unlabeled, mono-labeled, and bi-labeled species were generated during the labeling reaction. The mono-labeled fraction was purified using an anion exchange column (Mono Q 5/50 GL) and has a dye:protein ratio of ~0.9:1. The extinction coefficient of 250,000 $M^{-1}\,cm^{-1}$ at 650 nm was used for determining the Cy5 concentration. Similarly, the extinction coefficient of 9,700 $M^{-1}\,cm^{-1}$ at 280 nm was used for determining the Zur concentration; this extinction coefficient was calibrated using the BCA protein quantification assay.

##### **1.3.2** *Mutagenesis, expression, purification for $ZntR_{apo}$ (i.e., $ZntR_{C115S}$ mutant)*

ZntR(C115S) mutant was created, expressed, and purified as previously described[10,11]. Briefly, the mutant was cloned in a pET3a vector and expressed in *E. coli* (BL21 DE3) cells. The cells were grown until $OD_{600}$ ~ 0.6 before 0.4 mM IPTG was added. After an additional 3 hour growth at 37 °C, cells were harvested by centrifugation and then lysed with lysozyme in lysis buffer (50 mM Tris, 2 mM EDTA, 5 mM DTT at pH 8.0). The cells were further disrupted with three cycles of freeze and thaw and followed by sonication. The supernatant was collected after centrifugation and the proteins were precipitated out with 45% saturated $(NH_4)_2SO_4$ overnight. The precipitated proteins were resuspended in Tris buffer (20 mM Tris, 5 mM DTT at pH 8.0) and purified via a desalting column (HiPrep 26/10 Desalting, Cytiva), a Heparin affinity column (16/10 Heparin FF, GE Healthcare), and a gel filtration column (HILOAD 26/60 Superdex 200 PR, GE Healthcare). For the Heparin affinity column, the protein was stored in a buffer at pH 6 to increase binding affinity to the column. Protein purity was confirmed by SDS-PAGE, quantified using Bradford assay, and stored at −80 °C in 50 mM pH 8.0 Tris buffer with 250 mM NaCl, 5 mM DTT, and 5% glycerol. Protein identity was confirmed by mass spectrometry (HPLC-ESI-MS/MS, UT Health San Antonio, Supplementary Fig. 19 later).

##### **1.3.3** *Fluorescence labeled DNA preparation*

The Cy3 and biotin tagged DNA oligomeric strands were purchased from Integrated DNA Technologies (IDT, Coralville, IA) and dissolved in Nuclease-Free Duplex Buffer (IDT, Coralville, IA) and annealed together. Two types of double-strand DNA (dsDNA) constructs were used. The sequences of both constructs were from the *znuCB* gene promoter and contain the specific two-dyad sequence recognized by two Zur dimers: 5′/Cy3/AGAAGTGTGATATTATAACATTTCATGACTA-3′ and the complementary 5′-/Biotin-TEG/TAGTCATGAAATGTTATAATATCACACTTCT-3′. The other construct is a truncated DNA that has only one dyad sequence: 5′/Cy3/AGAAGTGTGATATTATAACATT-3′, and the complementary 5′-/Biotin-TEG/AATGTTATAATATCACACTTCT-3′.

##### 1.3.4 *Functionalization of slide and immobilization of DNA for in vitro studies*

A microfluidic channel containing the sample was formed by double-sided tape sandwiched between a quartz slide (Technical Glass Products, Inc. (TGP) and a borosilicate cover slip (Thermo Scientific). Quartz slides were first amine-functionalized with Vectabond (Vector Laboratories), followed by coating with biotinylated-polyethylene glycol (PEG) polymers (50:1 ratio of PEG and Biotin-PEG, Nanocs, m-PEG-SPA-5000 and biotin-PEG-NHS-3400) to reduce nonspecific protein and DNA adsorption on the quartz surfaces, and the biotinylated terminal group forms biotin-neutravidin linkages for immobilizing biotinylated DNA molecules (Extended Data Fig. 2b)[12,13]. Coverslips were also amine-functionalized using Vectabond and then coated with PEG polymers (Nanocs, 100 mg/mL m-PEG-SPA-5000). Quartz surfaces were further blocked using 2 mL BSA (0.1 mg/ml) to minimize non-specific binding. The neutravidin (Invitrogen) was introduced as 500 µL of 0.2 mg/mL in buffer solution (20 mM Tris, 200 nM $ZnSO_4$, 2 mM $MgCl_2$, 1 mM $CaCl_2$, 100 mM potassium glutamate, 1 mM TCEP at pH 8.0) and incubated for 15 min. After washing out unbound neutravidin, 1 mL of 100 pM Cy3-labeled biotinylated DNA solution in buffer flowed through the channel for immobilization. Then, the Cy5-labeled Zur solution containing an oxygen scavenging system (0.1 mg/mL glucose oxidase, 0.025 mg/mL catalase, 4% glucose, and 1 mg/mL Trolox)[14] in the same buffer, and if applicable, containing $ZntR_{apo}$, was flowed in continuously at a rate of 20 µL/min for fluorescence imaging.

##### 1.3.5 *In vitro FRET experiments scheme*

Single-molecule FRET experimental design and surface immobilization to probe Zur−DNA interactions are shown in Extended Data Fig. 2b and described in the text (Supplementary Information 1.3.4), similarly as we studied other protein-DNA or protein-protein interactions[13,15,16]. Double-strand DNA, in which one end of one strand had a FRET donor Cy3 and the other end of the other strand had biotin, was immobilized to the surface via a neutravidin-biotin linkage. The homo-dimeric Zur was labeled with a single FRET acceptor Cy5 and flew through the microfluidic channel across the immobilized DNA. Upon Zur binding to DNA, fluorescence intensities of Cy3 and Cy5 changed due to FRET. By monitoring the fluorescence intensities of Cy3 and Cy5 simultaneously, we studied Zur−DNA interactions in real-time.

##### 1.3.6 *Single-molecule FRET experiments and data analysis*

The single-molecule fluorescence experiments were performed using a prism-type total internal reflection microscope based on an Olympus IX71 inverted microscope, similarly as we previously reported[13,15,16]. The immobilized Cy3-labeled DNA was excited by a continuous-wave circularly polarized 532-nm laser (CrystaLaser, GCL-025-L-0.5%) of ~7 mW focused onto an area of ~94 × 68 $\mu m^2$ on the sample. The fluorescence of both Cy3 and Cy5 was collected by a 60× NA 1.2 water-immersion objective and split by a dichroic mirror into two channels using a Dual-View system (Optical Insights). The HQ550LP filter was used to reject the excitation laser light and each channel of fluorescence was further filtered (HQ580-60m or HQ660LP) and projected onto one-half of the imaging area of an EMCCD camera (Andor Ixon DV887) controlled by Andor IQ software. The time resolution for all the single-molecule experiments was 25 ms. All image analysis was done by custom-written codes in MATLAB (Supplementary Software 1). Individual Cy3 and Cy5 fluorescence intensity trajectories for immobilized DNA molecules interacting with Zur proteins were extracted from the fluorescence movie recorded by the camera. The FRET efficiency ($E_{\mathrm{FRET}}$) was computed as an approximation using the relationship: $I_{\mathrm{Cy5}}/(I_{\mathrm{Cy5}}+I_{\mathrm{Cy3}})$, where $I_{\mathrm{Cy3}}$ and $I_{\mathrm{Cy5}}$ are the fluorescence intensities. Then FRET trajectories that showed Cy3-Cy5 anti-correlated intensity fluctuation followed by a single photobleaching step were identified. The $E_{\mathrm{FRET}}$ histograms were compiled from hundreds of trajectories at each condition. In order to obtain higher resolution $E_{\mathrm{FRET}}$ histograms, a forward-backward non-linear (fnbl) filter was used to reduce the noise in the fluorescence trajectories[17,18] and thresholded to distinguish $E_{\mathrm{FRET}}$ states. $E_{\mathrm{FRET}}$ value of each state was taken from the original $E_{\mathrm{FRET}}$ trajectories to avoid value changes by fbnl filtering (Supplementary Fig. 4).

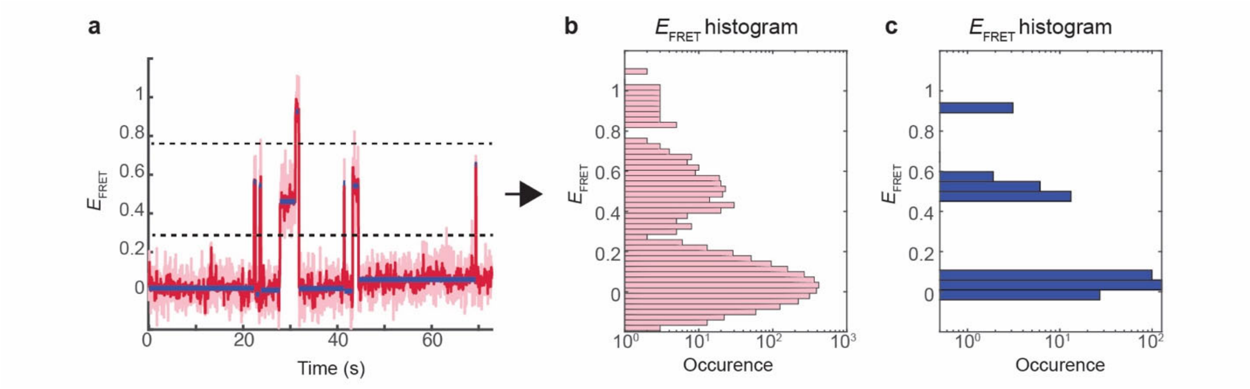


**Supplementary Fig. 4 | Construction of higher resolution $E_{FRET}$ histogram. a,** An example of $E_{FRET}$ trajectory before (pink lines) and after forward-backward nonlinear (fbnl) filtering (red lines) with two thresholds (horizontal dashed lines) to distinguish the three $E_{FRET}$ states. $E_{FRET}$ value for each state is the mean value of each state (blue lines) from the original trajectory before fbnl filtering. **b,** $E_{FRET}$ histogram from the original trajectory in (a). **c,** $E_{FRET}$ histogram re-constructed with $E_{FRET}$ values of individual states.

#### 2 Genome sequence analysis and identification of potential recognition sequences of metal efflux regulators (e.g., ZntR) at promoters that are regulated by metal uptake regulators (e.g., Zur), or vice versa, in *E. coli*, other bacteria, and yeast

In this section, we present sequence analysis to identify potential recognition sequences of metal efflux regulators (e.g., ZntR) at promoters that are regulated by metal uptake regulators (e.g., Zur), or vice versa, in *E. coli*, other bacteria, and yeast. Softwares SnapGene and ApE were used to search for a potential efflux regulator recognition sequence around the known uptake regulator binding boxes, or vice versa, in the promoter regions of its regulons.

##### 2.1 *Potential partial ZntR recognition sequences around known Zur boxes in E. coli and other bacteria*

The existence of sequences bearing some homology to known ZntR (or its homolog) recognition consensus sequence was observed around known Zur boxes in promoter regions in *E. coli, S. typhimurium,* and *P. aeruginosa*. In *E. coli*, these Zur regulon promoters include the promoters of genes *znuCB/znuA* ($Zn^{2+}$ uptake gene cluster), *zinT* (periplasmic $Zn^{2+}$ chaperone), *l31p* and *l33p* (a pair of ribosomal proteins), and *pliG* (a periplasmic lysozyme inhibitor) (Fig. 1b)[9]. In *S. typhimurium*, these include *zinT* and *znuABC* (Fig. 1b)[19,20].

In *P. aeruginosa*, the Zur regulon promoters include those of *PA5498* ($Zn^{2+}$ uptake gene cluster, including *znuA*), *PA0781, PA2911, PA4837 and PA1922* (putative TonB-dependent receptors), *PA5536* (*dksA2*, $Zn^{2+}$-independent global transcription regulator), *PA5539* (*folEB*, GTP cyclohydrolase), *PA4063* (periplasmic $Zn^{2+}$ binding protein), and *PA3600* (ribosomal proteins) (Extended Data Fig. 1a)[21]. In *P. aeruginosa*, Zn-efflux is regulated by CadR, a ZntR homolog of the MerR family that senses $Zn^{2+}$ and $Cd^{2+}$ as well; the recognition sequence of CadR is very similar to that of *E. coli* ZntR[22,23]. Extended Data Fig. 1a shows the Zur boxes (pink shade) in *P. aeruginosa* that are overlapping with potential CadR recognition sequences (blue arrows, where blue asterisks show matches with CadR recognition consensus sequence)[22].

##### 2.2 *Aside from the Zur-ZntR Zn uptake-efflux regulator pair, potential Zn efflux regulator recognition sequences are also found at promoters controlled by Zn uptake regulators of other families in bacteria.*

In *B. subtilis*, Zur regulates Zn uptake[24,25], while CzrA, a ArsR-family regulator, is a Zn-efflux regulator[26]. At the promoters of *zinT* ($Zn^{2+}$ trafficking protein), *znuABC* ($Zn^{2+}$ uptake gene cluster), *zagA*

($Zn^{2+}$ chaperone) and *folEB* (GTP cyclohydrolase), which are regulated by Zur[27,28], we also discovered potential recognition sequences for CzrA (Extended Data Fig. 1b).

In. *S. pneumoniae*, the MarR-family AdcR is another $Zn^{2+}$ uptake regulator[29], and SczA is another $Zn^{2+}$ efflux regulator of the TetR family[30]. At the promoter of *adcCBA* gene (a high affinity $Zn^{2+}$ importer) regulated by AdcR, there are also potential SczA recognition sequences (Extended Data Fig. 1c).

##### 2.3 *This pattern of potential existence of partial efflux regulator recognition sequences around known uptake regulator binding box was also observed for regulator pairs involved in the homeostasis of other metals beyond Zn (for example: Fe and Ni) in bacteria.*

In *B. subtilis*, Fur is a $Fe^{2+}$-uptake regulator while PerR is a $Fe^{2+}$-efflux regulator both belonging to the Fur family (i.e., same family as Zur)[31]. At the promoter of *dhbA* (gene involved in siderophore bacillibactin biosynthesis) regulated by Fur, we discovered potential PerR recognition sequence (Extended Data Fig. 1d).

In *E. coli*, NikR is a Ni-uptake regulator of the ribbon-helix-helix (RHH) family of DNA binding proteins[32], while RcnR is a Ni-efflux regulator of the CsoR family[33]. We discovered that at the promoter of gene *nikABCDE* (a $Ni^{2+}$ transport system) regulated by NikR, there are also potential recognition sequence of RcnR (Extended Data Fig. 1e).

##### 2.4 *Oppositely, uptake regulator recognition sequence is also found around known efflux regulator binding box in bacteria.*

Additionally, in *B. subtilis*, Helmann et al. observed the existence of two potential Fur (uptake regulator) binding sites around the known PerR (efflux regulator) binding box[34]; they also demonstrated the physiological importance of these two additional Fur boxes in iron efflux. It is worth noting that in this case, the recognition sequence of uptake regulator exists near efflux regular binding site, as opposed to the case of the recognition sequence of efflux regulator (e.g., ZntR) exists near the binding site of uptake regulator (e.g., Zur), which suggests a broader relevance of potential uptake-efflux regulator cross-actions on DNA.

##### 2.5 *In yeast, a similar pattern of efflux regulator recognition sequence overlapping with known uptake regulator binding motifs can be found for iron homeostasis.*

In yeast (*S. cerevisiae*), zinc homeostasis does not utilize pairs of efflux-uptake regulators and instead both uptake and efflux are controlled by one regulator; therefore the balance of cellular Zn concentration follows a different mechanism[35–37].

On the other hand, for iron homeostasis, yeast uses distinct transcription factors to tightly regulate iron uptake and efflux[38]. During conditions of iron deficiency, the two transcription factors, Aft1 and Aft2, bind to iron-regulatory promoter elements (FeREs) to activate the expression of genes involved in iron uptake, mobilization, and recycling, known as the iron regulons[38,39]. When cellular iron reaches toxic levels, the regulators Yap5, Msn2, and Msn4 activate the expression of Ccc1, which detoxifies excess cytosolic iron by importing it into the vacuole for mobilization during deficiency[38–40]. Here, we discovered that around the known Aft1/2 binding boxes on the promoter region of the iron regulon (e.g., *fre1*), there exist sequences that bear partial similarities with the known Yap5 and Msn2/4 binding motifs (Extended Data Fig. 1f).

#### 3 Functionality and intactness of sfGFP-tagged ZntR in *E. coli* cells

##### 3.1 *Western blot shows the intactness of ZntR-sfGFP fusion proteins*

We previously showed that mEos3.2 tagged Zur is a functional regulator[2]. The fusion tag remained largely intact in the cell (i.e., no discernible cleavage for the tagged Zur). We previously also showed that ZntR, tagged with mEos3.2 (a GFP variant), is also functional in the cell, but the mEos3.2 tag there has some cleavage in the cell[3].

For sfGFP tagged ZntR that we use in this study, we performed western blot to check its intactness in the cells, focusing on the ZntR(C115S)-sfGFP fusion as the representative (i.e., $ZntR^{G}_{apo}$).

An anti-GFP antibody was used for immunoblotting. The DZ-DZR-pZRGC115S strain which could express $ZntR^{G}_{apo}$ from the pBAD33 plasmid inducible by L-arabinose, and a negative control strain DZ-DZR-pBAD33 containing the parent pBAD33 vector without insert were cultured overnight (18 hours) in 6 mL LB with appropriate antibiotics. A dilution (1:100) of the overnight culture was done in 5 mL M9 medium with amino acids (8% v/v 50x GIBCO), vitamins (4% 100x GIBCO), glycerol (0.4%) and the samples were further grown to an OD600 of 0.4. L-arabinose was then added to a final concentration of 1 mM into the appropriate cultures which were further incubated for 30 min to induce the plasmid expression. 1 mL aliquots of the resulting cell cultures were collected by centrifugation and washed with the same M9. The cell pellets were re-suspended in 95 µL 2X SDS lysis buffer (BIORAD 2X Laemmli sample buffer), 2.5 µL BME (2-mercaptoethanol; Sigma-Aldrich) and 2.5 µL protein inhibitor cocktail (Promega). The lysed samples were run in SDSPAGE with ECL Plex fluorescent rainbow protein molecular weight markers (GE Healthcare Life Science) in 1X Running buffer, and then transferred onto the Hybond-LEP PVDF membrane (GE Healthcare Life Sciences). 4% Amersham ECL Prime blocking reagent (GE Healthcare Life Sciences) in PBS-T (0.1% Tween-20, Sigma-Aldrich) wash buffer was used to block the transferred membrane while shaking at RT for 2 hours. The membrane was washed with PBS-T and incubated with rabbit-derived antiGFP primary antibody (1:10,000 dilution, Rockland Immunochemical) for 18 hours at 4 ℃. The membrane was rinsed with PBS-T 4 times and PBS buffer 3 times. The goat-derived Horseradish Peroxidase-conjugated Fab fragment anti-rabbit antibody (1:20,000 dilution, Rockland Immunochemical) was used as the secondary antibody, which could be probed with SuperSignal West Femto Maximum Sensitivity Substrate (Fisher Scientific). BioRad ChemiDoc MP Imaging System was used to detect peroxidase activity.

A dominant band from L-arabinose induced $ZntR^{G}_{apo}$ was observed at MW ~ 43 kDa (that is, MW of ZntR + MW of sfGFP), and no discernable band was observed at MW ~ 27 kDa, which is expected to be the MW of sfGFP (Supplementary Fig. 5). Therefore, $ZntR^{G}_{apo}$ is intact in the cell.

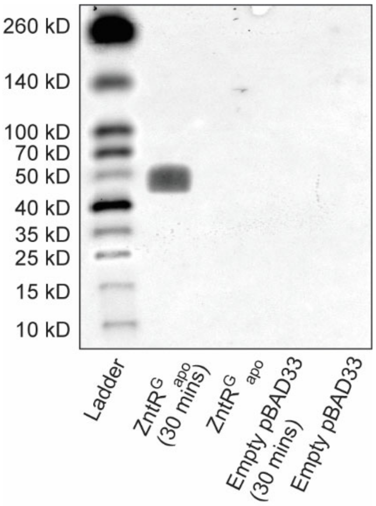


**Supplementary Fig. 5 |** Western blot demonstrated that $ZntR^{G}_{apo}$ is intact in the cell. Only $ZntR^{G}_{apo}$ expressed from a pBAD33 plasmid induced with L-arabinose was detectable (2nd column). In the negative controls, which are un-induced $ZntR^{G}_{apo}$ encoded in the same pBAD33 plasmid (3rd column), the empty pBAD33 (without the $ZntR^{G}_{apo}$ insert) induced with L-arabinose (4th column)

and un-induced (5th column), no band was observed. The expected size of $\mathrm{ZntR}^{\mathrm{G}}_{\mathrm{apo}}$ is ~ 43 kDa, and no cleavage product was observed (2nd column).

##### 3.2 *Ensemble fluorescence measurements show that the sfGFP-tagged ZntR is fluorescent*

To test whether the sfGFP tag on ZntR is fluorescent inside cells, we performed bulk fluorescence measurements, using $\mathrm{ZntR}^{\mathrm{G}}_{\mathrm{apo}}$ as the representative. Strain DZ-DZR-pZRGC115S, which harbors the zntR-C115S-sfGFP gene encoded in a pBAD33 plasmid (Supplementary Table 2), was cultured in 6 mL LB with chloramphenicol (25 μg/mL), kanamycin (30 μg/mL) and 5 mM L-arabinose in 37 °C, shaking at 250 rpm, for 18 hours. The cells were then centrifuged, and the pellet was washed and re-suspended in PBS buffer (pH 7.4) for fluorescence measurements (Agilent Eclipse fluorometer). The emission spectrum of the green-fluorescent sfGFP was collected using 465 nm excitation; its excitation spectrum was obtained by monitoring the emission at 530 nm. Supplementary Fig. 6 shows the emission and excitation spectra of $\mathrm{ZntR}^{\mathrm{G}}_{\mathrm{apo}}$ in the cells. The spectra closely match those expected for the sfGFP[4]. This shows that the sfGFP component of the fusion gene expressed inside the cell is fluorescent.

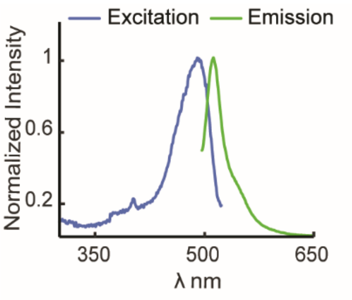


**Supplementary Fig. 6 |** The excitation (blue; emission detected at 530 nm) and emission (green; emission excited at 465 nm) spectra of cells expressing $\mathrm{ZntR}^{\mathrm{G}}_{\mathrm{apo}}$ suspended in a PBS buffer.

#### 4 Analysis of resolvable diffusion states of Zur in the cell and extraction of their effective diffusion coefficients (*D*) and fractional populations (*A*)

##### 4.1 *The resolved three diffusion states of Zur in the cell were assigned as FD, NB, or TB based on their diffusion coefficients and rationales*

The distribution of the displacement lengths, *r*, obtained from tracking individual mE-tagged Zur proteins in two-dimension was fitted with a scaled probability distribution function ($\mathrm{PDF_{2D}}$) (Fig. 2b, Eq. S6) or a cumulative distribution function ($\mathrm{CDF_{2D}}$) (Eq. S7), using Brownian diffusion model to determine the number of resolvable diffusion states and their respective diffusion coefficients and fractional populations, as we previously did in studying Zur-DNA interactions[2] and ZntR-DNA interactions in *E. coli* cell[3]:

$$\mathrm{PDF_{2D}}\,(r) = N \sum_i A_i \left(\frac{r}{2D_i T_{tl}} \left(\exp\left(-\frac{r^2}{4D_i T_{tl}}\right)\right)\right) \qquad \text{Eq. S6}$$

$$\mathrm{CDF_{2D}}\,(r) = \sum_i A_i \left(1 - \exp\left(-\frac{r^2}{4D_i T_{tl}}\right)\right) \qquad \text{Eq. S7}$$

Here $N$ is a scaling constant; $D_\mathrm{i}$ is the effective diffusion coefficient of state $i$ whose fractional population is $A_\mathrm{i}$, and $\sum A_i = 1$. $T_\mathrm{tl}$ is the time lapse (40 ms) in our time-lapse stroboscopic imaging. A linear combination of three Brownian diffusion states in the CDF was used here, assuming a quasi-static approximation, as we did previously because our measurement time resolution (40 ms) is faster than the interconversions between the states[2,3].

Only the first displacement of each experimentally obtained tracking trajectory was used to avoid biasing towards longer trajectories. Since the experimental PDF of displacement length requires a choice of bin size, fitting the CDF was preferred, and fitted parameters can be used to generate the corresponding fit for the PDF of displacement lengths.

From our single-cell protein quantitation of $Zur^{mE}$ and $ZntR^{G}_{apo}$ (or $ZntR^{G}$), we first sorted individual cells by their [$ZntR^{G}_{apo}$] (or [$ZntR^{G}$]) concentrations into groups of similar protein concentrations (e.g., horizontal dashed lines in Supplementary Fig. 7). Then within each group of cells having similar ZntR concentration, they are further sorted by their [$Zur^{mE}$] concentrations into groups having similar protein concentrations (e.g., vertical dashed lines in Supplementary Fig. 7).

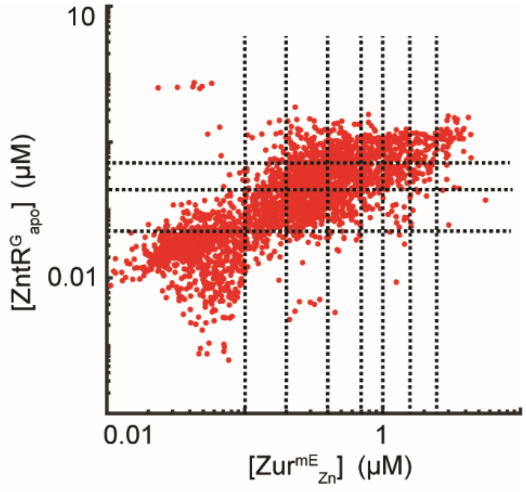


**Supplementary Fig. 7 |** Example of a two-dimensional scatter plot of [$Zur^{mE}_{Zn}$] vs [$Zur^{G}_{apo}$] of ~4300 cells for the DZ-DZR-pZmE-pZRGC115S strain expressing both proteins from plasmids; each red dot represents one cell. The horizontal dashed black lines divide cells into four different [$ZntR^{G}_{apo}$] groups, each of which was further divided into 6-8 [$Zur^{mE}$] groups by the vertical dashed black lines. The division lines were chosen to ensure that each concentration group in general have several thousands of single-molecule tracking trajectories of $Zur^{mE}_{Zn}$.

A global CDF fit was performed across these Zur concentration groups sharing the $D_i$'s because the diffusion coefficient is more likely an intrinsic property of each diffusion state and expected to be independent of protein concentration. Their respective fractional populations ($A_i$) are allowed to differ among different cell groups. Analyzing the residuals of the fit, minimally three diffusion states were required for the fitting the results of Zur, as we observed earlier[2].

Supplementary Fig. 8 shows exemplary CDF fits across different $Zur_{Zn}$ concentration groups at [$ZntR_{apo}$] = 27 ± 8 nM (error bar here is the standard deviation among the individual cells in the group). The effective diffusion coefficients of the states, $D_i$'s, and their respective fractional populations, $A_i$'s, across different concentrations of Zur and ZntR are summarized in Supplementary Table 5 and Supplementary Table ***6***.

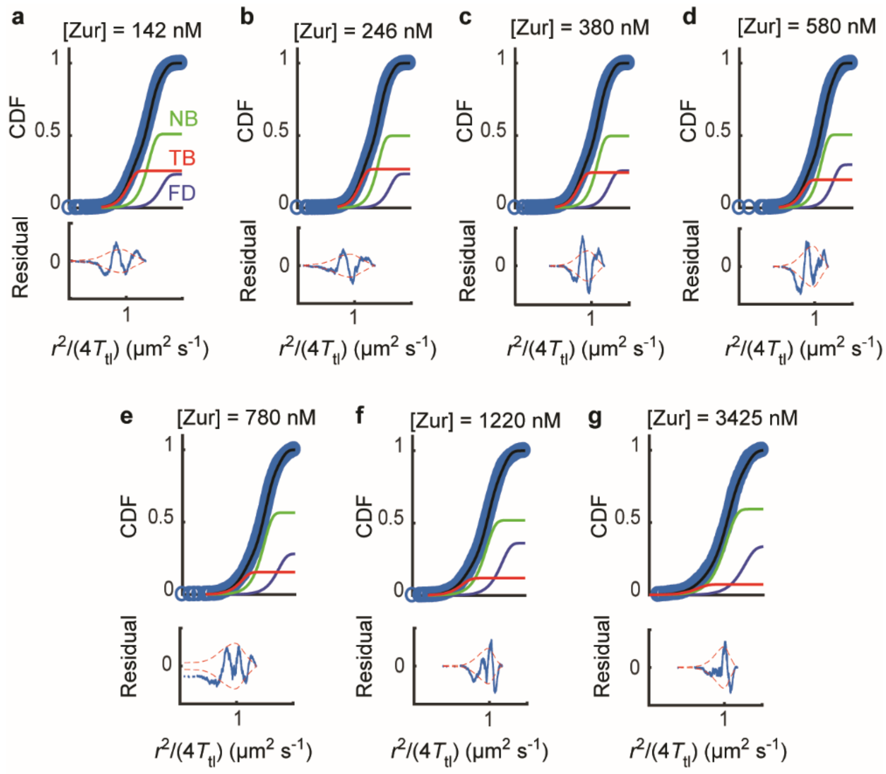


**Supplementary Fig. 8 | Extraction of Zur$^{mE}$ diffusion coefficients and fractional populations.** (**a-g**) Top: exemplary global CDF fits for cells with [ZntR$^{G}_{apo}$] = 27 ± 8 nM across different Zur$^{mE}_{Zn}$ concentration groups. The red, green, and blue curves represent the resolved TB, NB, and FD states. The black lines are the overall fits of the CDFs. Bottom: the residuals in blue are shown with the 95% confidence bounds in red.

The assignments of the three resolved states of Zur in the *E. coli* were reported, rationalized, and justified in our previous study[2]. The fastest diffusion state of Zur$^{mE}$ has an effective diffusion coefficient $D_{FD} \approx 5$ μm/s$^2$ and was assigned to those Zur proteins freely diffusing in the cytosol (i.e., FD state); the slowest diffusion state has an effective diffusion coefficient $D_{TB}$ = 0.04 μm/s$^2$ and was assigned as Zur proteins tightly bound to DNA (e.g., at a Zur box) (i.e., TB state). The slight motion of TB state reflects chromosome dynamics and our experimental localization uncertainties of ~12 nm[3,41–45]. The intermediate diffusion state has an effective diffusion coefficient $D_{NB}$ = 0.4 μm/s$^2$ and was assigned to Zur non-specifically binding to DNA (i.e., NB state).

The values obtained for the diffusion coefficients are similar to what were previously observed for Zur[2], other metal-sensing regulators[3] and other transcription factors in *E. coli*[45–48]. It is worth noting that these are effective diffusion coefficients, not intrinsic diffusion coefficients, as the effective values are influenced by the cell-confinement effects and the experimental time resolution (i.e., time lapse between images). The difference between effective and intrinsic diffusion coefficients are minimal for the TB and NB states as they are either quasi-stationary or slow, and the difference for FD state is larger, but FD state is always the fastest among the three states, as we evaluated previously[2,3].

**Supplementary Table 5 |** Summary of all the extracted Zur$^{mE}$ diffusion coefficients of the three different states.

| Proteins in the Strain | $D_{FD}$ (μm$^2$ s$^{-1}$) | $D_{NB}$ (μm$^2$ s$^{-1}$) | $D_{TB}$ (μm$^2$ s$^{-1}$) |
|---|---|---|---|
| $Zur_{Zn}$ , $ZntR_{apo}$ | 6.24±0.62 | 0.939±0.057 | 0.0532±0.0074 |
| $Zur_{C88S}$ , $ZntR_{apo}$ | 5.46±0.59 | 0.723±0.14 | 0.0411±0.0024 |
| $Zur_{Zn}$ , $ZntR_{Zn}$ | 5.04±0.23 | 0.903±0.04 | 0.0426±0.0042 |

**Supplementary Table 6 |** Summary of all the extracted populations of the three different $Zur^{mE}$ diffusion states.

| [$ZntR_{apo}$] (nM) | [$Zur_{Zn}$] (nM) | $A_{FD}$ (%) | $A_{NB}$ (%) | $A_{TB}$ (%) |
|---|---|---|---|---|
| 0±0 | 99±34 | 15.5±0.3 | 52.1±1 | 32.4±0.6 |
| | 174±15 | 18.8±0.4 | 53.3±1.1 | 27.9±0.6 |
| | 313±84 | 22.7±0.5 | 50.2±1 | 27.1±0.5 |
| | 633±81 | 28.7±0.6 | 49.5±1 | 21.7±0.4 |
| | 998±117 | 29.6±0.6 | 54±1.1 | 16.4±0.3 |
| | 1644±363 | 28.3±0.6 | 57.3±1.1 | 14.4±0.3 |
| | 3573±873 | 31.2±0.6 | 60.3±1.2 | 8.5±0.2 |
| 27±8 | 142±41 | 23.1±0.5 | 51.2±1 | 25.7±0.5 |
| | 246±29 | 23.5±0.5 | 49.7±1 | 26.7±0.5 |
| | 380±53 | 25.9±0.5 | 49.6±1 | 24.4±0.5 |
| | 582±59 | 30±0.6 | 50.6±1 | 19.5±0.4 |
| | 779±59 | 28.1±0.6 | 56.4±1.1 | 15.5±0.3 |
| | 1217±376 | 36.4±0.7 | 51.8±1 | 11.9±0.2 |
| | 3425±954 | 33.6±0.7 | 59.2±1.2 | 7.2±0.1 |
| 114±15 | 366±95 | 22.1±0.4 | 55.4±1.1 | 22.5±0.4 |
| | 596±61 | 23.2±0.5 | 58.8±1.2 | 18.1±0.4 |
| | 800±58 | 23.6±0.5 | 60.3±1.2 | 16.1±0.3 |
| | 953±30 | 24.1±0.5 | 59.9±1.2 | 16±0.3 |
| | 1146±130 | 25.5±0.5 | 59.2±1.2 | 15.3±0.3 |
| | 1867±254 | 24.8±0.5 | 64.7±1.3 | 10.5±0.2 |
| | 3311±1010 | 30.4±0.6 | 62.2±1.2 | 7.5±0.1 |
| 316±19 | 531±115 | 31±0.6 | 50.5±1 | 18.5±0.4 |
| | 800±61 | 30.4±0.6 | 54.8±1.1 | 14.8±0.3 |
| | 950±29 | 27.3±0.5 | 58.5±1.2 | 14.2±0.3 |
| | 1164±135 | 29.7±0.6 | 57±1.1 | 13.3±0.3 |
| | 1986±259 | 30±0.6 | 58.4±1.2 | 11.5±0.2 |
| | 3373±797 | 24.5±0.5 | 64.3±1.3 | 11.2±0.2 |

| [$ZntR_{apo}$] (nM) | [$Zur_{C88S}$] (nM) | $A_{FD}$ (%) | $A_{NB}$ (%) | $A_{TB}$ (%) |
|---|---|---|---|---|
| 10±8 | 89±9 | 22.4±0.4 | 32.6±0.7 | 45±0.9 |
| | 155±29 | 22.7±0.5 | 46.7±0.9 | 30.6±0.6 |
| | 219±6 | 29.6±0.6 | 41.8±0.8 | 28.6±0.6 |
| | 262±21 | 26.8±0.5 | 43.4±0.9 | 29.8±0.6 |
| | 398±86 | 30.6±0.6 | 45.3±0.9 | 24.1±0.5 |
| | 975±370 | 35.9±0.7 | 56.3±1.1 | 7.8±0.2 |
| | 107±27 | 18.9±0.4 | 46.9±0.9 | 34.2±0.7 |
| | 260±74 | 24.1±0.5 | 53.5±1.1 | 22.4±0.4 |
| | 447±28 | 23.6±0.5 | 54.5±1.1 | 21.9±0.4 |

| | | | | |
|---|---|---|---|---|
| 70±30 | 599±58 | 24.2±0.5 | 62.1±1.2 | 13.6±0.3 |
| | 802±51 | 25±0.5 | 63.9±1.3 | 11.1±0.2 |
| | 1364±353 | 28.4±0.6 | 64.3±1.3 | 7.4±0.1 |
| | 2351±96 | 31.5±0.6 | 61.7±1.2 | 6.8±0.1 |
| | 3054±477 | 32.1±0.6 | 65.3±1.3 | 2.6±0.1 |
| | 464±118 | 15.1±0.3 | 67.1±1.3 | 17.8±0.4 |
| | 675±47 | 18±0.4 | 66.8±1.3 | 15.2±0.3 |
| | 993±136 | 19.6±0.4 | 66.7±1.3 | 13.7±0.3 |
| 247±72 | 1424±145 | 21.4±0.4 | 70.9±1.4 | 7.8±0.2 |
| | 1802±55 | 21.2±0.4 | 69.8±1.4 | 9±0.2 |
| | 2145±170 | 22.2±0.4 | 69.9±1.4 | 7.9±0.2 |
| | 3269±616 | 25.1±0.5 | 68.6±1.4 | 6.3±0.1 |

| [$ZntR_{Zn}$] (nM) | [$Zur_{Zn}$] (nM) | $A_{FD}$ (%) | $A_{NB}$ (%) | $A_{TB}$ (%) |
|---|---|---|---|---|
| | 78±11 | 14.6±0.3 | 33.7±0.7 | 51.8±1 |
| | 173±55 | 24.1±0.5 | 34.2±0.7 | 41.7±0.8 |
| 27±7 | 345±30 | 27.5±0.6 | 30.2±0.6 | 42.3±0.8 |
| | 491±56 | 34±0.7 | 36.4±0.7 | 29.6±0.6 |
| | 698±56 | 42.4±0.8 | 23.4±0.5 | 34.2±0.7 |
| | 1187±391 | 49.9±1 | 32.1±0.6 | 18±0.4 |
| | 136±40 | 8.9±0.2 | 29.3±0.6 | 61.8±1.2 |
| | 249±26 | 12.3±0.2 | 26.4±0.5 | 61.3±1.2 |
| | 379±56 | 17.5±0.3 | 31.8±0.6 | 50.7±1 |
| 64±3 | 709±150 | 20.6±0.4 | 38.9±0.8 | 40.5±0.8 |
| | 1198±140 | 24.6±0.5 | 53.5±1.1 | 22±0.4 |
| | 1794±271 | 26.6±0.5 | 51.3±1 | 22.1±0.4 |
| | 3171±664 | 25.2±0.5 | 64.8±1.3 | 10±0.2 |
| | 149±35 | 14±0.3 | 19.3±0.4 | 66.7±1.3 |
| | 251±29 | 16.9±0.3 | 29±0.6 | 54.1±1.1 |
| | 368±43 | 17.7±0.4 | 28.3±0.6 | 53.9±1.1 |
| 131±5 | 517±42 | 20.3±0.4 | 38±0.8 | 41.7±0.8 |
| | 788±117 | 23.4±0.5 | 45.7±0.9 | 31±0.6 |
| | 1237±151 | 26.9±0.5 | 53.2±1.1 | 19.9±0.4 |
| | 1952±275 | 31.5±0.6 | 49.6±1 | 18.8±0.4 |
| | 3366±643 | 33±0.7 | 58.1±1.2 | 8.9±0.2 |
| | 178±74 | 13.5±0.3 | 28±0.6 | 58.5±1.2 |
| | 399±57 | 16.4±0.3 | 32.2±0.6 | 51.4±1 |
| | 543±29 | 17.2±0.3 | 30.9±0.6 | 51.9±1 |
| 407±48 | 690±58 | 16.3±0.3 | 39.4±0.8 | 44.3±0.9 |
| | 1123±221 | 26±0.5 | 43.8±0.9 | 30.2±0.6 |

| | | | |
|---|---|---|---|
| 1945±285 | 29.6±0.6 | 53.9±1.1 | 16.5±0.3 |
| 3664±771 | 30.7±0.6 | 57.9±1.2 | 11.4±0.2 |

##### 4.2 *The fractional populations of Zur's three diffusion states show expected dependence on cellular [Zur]*

For all the conditions it was expectedly observed that the $A_{FD}$ increases and $A_{TB}$ decreases with increase in cellular [Zur]. Supplementary Fig. 9 shows an exemplary plot of fractional populations of the three diffusion states vs. $Zur^{mE}$ concentration for the DZ-DZR-pZmE-pZRGC115S strain expressing both the $ZntR^{G}_{apo}$ and $Zur^{mE}$ proteins. These trends of $A_{FD}$ and $A_{TB}$ are expected because there are only a limited number of chromosomal binding sites for Zur and as [Zur] increases, there is more competition between the proteins for a limited number of tight binding sites, so each protein molecule spends larger fraction of its time in the free diffusion state than in the tight binding state.

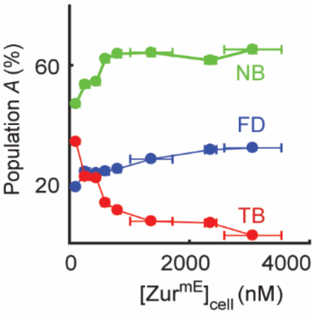


**Supplementary Fig. 9 |** Example plot of Zur protein concentration dependence of fractional populations of the TB, NB, and FD states, from the DZ-DZR-pZmE-pZRGC115S strain expressing both the $ZntR^{G}_{apo}$ and $Zur^{mE}$ proteins from plasmids (Supplementary Table 3). The data here did not sort the cells by [$ZntR^{G}_{apo}$] in the cell.

##### 4.3 *Bootstrap analysis shows statistical reliability of data*

A bootstrap analysis was performed by sampling randomly 50%, 75%, 85%, and 95% of the displacements lengths, to show that the extracted results from analyzing the CDF of the displacement lengths *r* are statistically reliable. The ratios of these extracted diffusion coefficients (*D*'s) and fractional populations (*A*'s) over the results in Supplementary Table 5 andSupplementary Table ***6*** clearly show that the extracted results from random sampling are all within 3% of the final results (Supplementary Fig. 10). This indicates that with just 50% of the experimental data we collected, we can determine diffusion coefficients and the corresponding fractional populations reliably, supporting the statistical saturation of our data.

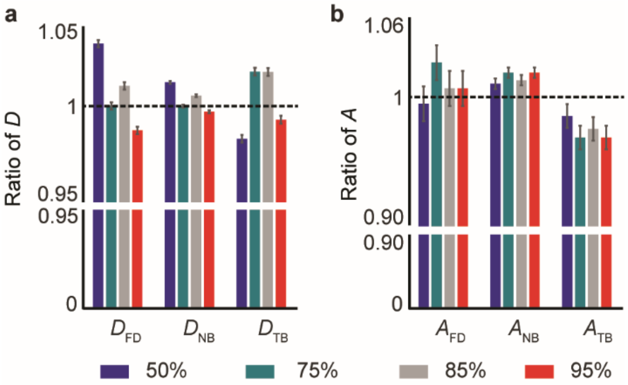


**Supplementary Fig. 10 | Bootstrap analysis show that our experimental data are statistically saturated**. Ratio of **a**, D's and **b**, A's by analyzing the CDF of a random subset of displacement lengths (i.e., 50%, 75%, 85%, and 95% of data) to the final results

(100% of the data) in Supplementary Table 5 and Supplementary Table **6**. The ratios are all within the range of ~0.97 to ~1.03, demonstrating that the results are within 3% of the final results and hence statistically saturated.

### 5 Extraction of kinetic and thermodynamic parameters for Zur-DNA interactions in cells

#### 5.1 *Extraction of $k_{-1}$, the apparent unbinding rate constant from the tight binding state*

The quantitative analysis of Zur's unbinding kinetics from the TB sites on the chromosome follows our previous work[2], where the formulation and derivation of the kinetic model and its validation are described in detail. Here we briefly summarize the treatment.

Zur proteins that are bound to chromosomal sites tightly are almost immobile, giving rise to the TB state resolved (Supplementary Fig. 8; Fig. 2b) in the analysis of their displacement distributions with an effective diffusion coefficient $D_{TB}$ = 0.04 μm$^2$ s$^{-1}$ and correspondingly small displacement length $r$ between adjacent images (Supplementary Fig. 11b). Thresholding the displacement $r$-vs-time trajectories with an upper limit $r_0$ (= 200 nm), which includes >99.5% of the TB state (Fig. 2b; Supplementary Fig. 11a) yields the microscopic residence times $\tau$ that are dominated by Zur tightly bound on DNA[2,3]. A residence time starts upon transition from larger values of $r$ to below $r_0$ and terminates upon transitions above $r_0$ (or by photobleaching/blinking of the fluorescent mE tag) (Supplementary Fig. 11a). The distribution of microscopic residence times $\tau$ extracted from many displacement-vs-time trajectories can be analyzed to determine the apparent unbinding rate constant $k_{-1}$ of Zur from the TB sites, using the 3-state kinetic model in Fig. 2c[2,3], while taking into account the photobleaching/blinking kinetics of the mE tag.

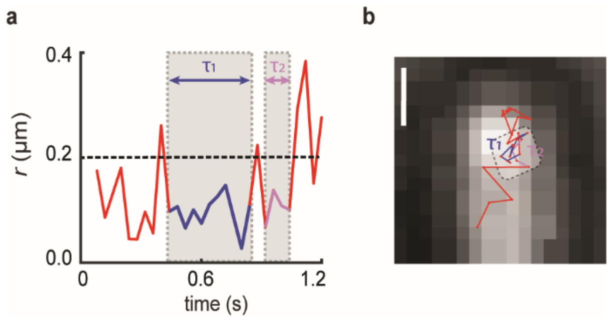


**Supplementary Fig. 11 | Extraction of Zur's microscopic residence time on tight-binding sites on DNA. a**, Exemplary displacement length $r$-vs-time trajectory of a single Zur$^{mE}$ in an *E. coli* cell. The dashed horizontal line represents the $r_0$ threshold (~200 nm), below which >99.5% of TB state are included (vertical dashed line in Fig**.** 2b). A microscopic residence time $\tau$ begins when the displacement goes below $r_0$ and it ends when it goes above $r_0$ or when the tag photobleaches or photoblinks. Here in this example, $\tau_1$ and $\tau_2$ represent two residence times, denoted by the gray shade and blue/pink trajectory segments. **b**, The position-vs-time trajectory shows the residence sites corresponding to the two residence times denoted by the blue/pink trajectory segments in (a). The scale bar is 200 nm.

Experimentally, the microscopic residence time $\tau$ could also be terminated by a photobleaching/blinking event of the mE tag. The photobleaching/blinking rate constant $k_{bl}$ can be determined from analyzing the distribution of the fluorescence on-times (Supplementary Fig. 12b) extracted from the corresponding fluorescence intensity trajectories of single-molecule tracking trajectories (Supplementary Fig. 12a). The distribution of on-time was fitted with Eq. S8, as we previously reported[2,3].

$$f_{on}(t) = N \exp\left(-k_{bl}\frac{T_{int}}{T_{tl}}t\right) \qquad \text{Eq. S8}$$

Since we used time-lapse stroboscopic imaging, the apparent photobleaching/blinking rate constant from the fluorescence on-time distribution has been corrected by the ratio of the laser integration time $T_{int}$ (= 4 ms) and the time-lapse $T_{tl}$ (= 40 ms)[2,3]. $N$ is a normalization constant. The extracted $k_{bl}$, 230 ± 10 $s^{-1}$, is consistent with the reported values under similar 561 nm excitations of the mE tag[2,3,49,50].

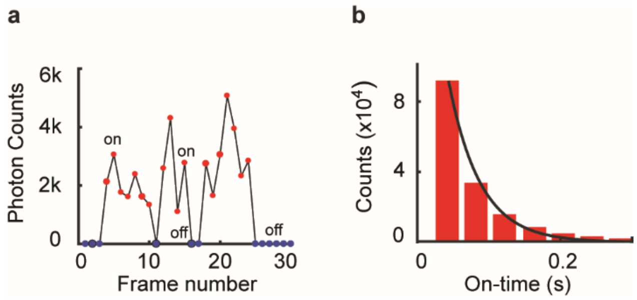


**Supplementary Fig. 12 | Determination of the Photobleaching and blinking rate constant $k_{bl}$. a**, Exemplary fluorescence-vs-time trajectory of a mEos3.2 tagged Zur protein. The red dots represent frames where mEos3.2 fluorescence is detected and assigned as fluorescence-on frames. The blue dots correspondingly represent the off-frames with no protein fluorescence detection. **b**, The distribution of the fluorescence on-times, fitted with Eq. S8 (black line), yields the $k_{bl}$ (photobleaching/blinking rate constant).

Since diffusion is a probabilistic process at the microscopic level, freely diffusing or non-specifically bound proteins, which are expected to have large displacements in general ($D_{FD}$ = 5 μm$^2$ s$^{-1}$ and $D_{NB}$ = 0.82 μm$^2$ s$^{-1}$), have finite probabilities to have small displacement lengths as well. These will thus contribute to the microscopic residence times thresholded by $r_0$. Thus, below $r_0$, where >99.5% TB states are included, the contributions of the FD and NB states are 4.9% and 26.3% of their populations, respectively (Fig. 2b), and these need to be deconvoluted. To separate the contributions of FD and NB states from TB in the distribution of microscopic residence time, a survival probability S($r_0$, t), which is the probability for a protein at the origin to survive within a circle of radius $r_0$ within time $t$, was calculated, as we described previously[2,3].

$$S(r_0, t) = \left[1 - \exp\left(-\frac{r_0^2}{4Dt}\right)\right] \exp(-k_{eff}t) \qquad \text{Eq. S9}$$

where $D$ is the diffusion constant of the protein and $k_{eff}$ is the summation of unbinding rate constants (applicable for the TB and the NB state only) and the effective photobleaching/blinking rate constant (i.e., $k_{bl}\frac{T_{int}}{T_{tl}}$). The overall survival probability for a Zur protein within $r_0$ is a linear combination of survival probabilities of each state weighted by its fractional population:

$$S_{all}(r_0,t) = A_{FD}S_{FD}(r_0,t) + A_{NB}S_{NB}(r_0,t) + A_{TB}S_{TB}(r_0,t) \qquad \text{Eq. S10}$$

Then, the respective probability distribution function of the thresholded residence time $\tau$, for the FD, NB, and TB states (i.e., $\varphi_{FD}(\tau)$, $\varphi_{NB}(\tau)$, and $\varphi_{TB}(\tau)$) can be obtained by taking a time-derivative of the survival probability (i.e., $\varphi(\tau) = -\frac{\partial S(t)}{\partial t}|_{t=\tau}$ ):

$$\varphi_{all}(\tau) = A_{FD}\varphi_{FD}(\tau) + A_{NB}\varphi_{NB}(\tau) + A_{TB}\varphi_{TB}(\tau) \qquad \text{Eq. S11}$$

$$\varphi_{FD}(\tau) = \left[\frac{r_0^2}{4D_{FD}\tau^2}\exp\left(-\frac{r_0^2}{4D_{FD}\tau}\right) + k_{eff}^{FD}\left(1 - \exp\left(-\frac{r_0^2}{4D_{FD}\tau}\right)\right)\right]\exp\left(-k_{eff}^{FD}\,\tau\right) \qquad \text{Eq. S12}$$

$$\varphi_{\mathrm{NB}}(\tau) = \left[\frac{r_0^2}{4D_{\mathrm{NB}}\tau^2}\exp\left(-\frac{r_0^2}{4D_{\mathrm{NB}}\tau}\right) + k_{\mathrm{eff}}^{\mathrm{NB}}\left(1-\exp\left(-\frac{r_0^2}{4D_{\mathrm{NB}}\tau}\right)\right)\right]\exp\left(-k_{\mathrm{eff}}^{\mathrm{NB}}\,\tau\right) \quad \text{Eq. S13}$$

$$\varphi_{\mathrm{TB}}(\tau) = k_{\mathrm{eff}}^{\mathrm{TB}}\exp\left(-k_{\mathrm{eff}}^{\mathrm{TB}}\,\tau\right) \quad \text{Eq. S14}$$

Here, $k_{\mathrm{eff}}^{\mathrm{FD}} = k_{\mathrm{bl}}\frac{T_{\mathrm{int}}}{T_{\mathrm{tl}}}$, $k_{\mathrm{eff}}^{\mathrm{NB}} = k_{\mathrm{bl}}\frac{T_{\mathrm{int}}}{T_{\mathrm{tl}}} + k_{-2}$ and $k_{\mathrm{eff}}^{\mathrm{TB}} = k_{\mathrm{bl}}\frac{T_{int}}{T_{\mathrm{tl}}} + k_{-1}$ (see Fig. 2c for definition of rate constants).

$k_{-2}$ is the unbinding rate constant from the NB sites. $k_{-2}$ was extracted from the highest cellular concentration regime by fitting the residence time distribution with Eq. S15, in which $A_{\mathrm{TB}}$ is <10% and the $A_{\mathrm{TB}}\varphi_{\mathrm{TB}}(\tau)$ term in Eq. S11 becomes negligible:

$$\varphi_{\mathrm{all}}(\tau) = A_{\mathrm{FD}}\varphi_{\mathrm{FD}}(\tau) + A_{\mathrm{NB}}\varphi_{\mathrm{NB}}(\tau) \quad \text{Eq. S15}$$

$k_{-1}$ was extracted by fitting the residence time distribution from all the other concentration groups with Eq. S11 with predetermined $D$'s, $A$'s, $k_{\mathrm{bl}}$, and $k_{-2}$. All determined rate constants are summarized in Extended Data Table 1. This method of extracting $k_{-1}$ from the residence time distribution was further validated by using simulation data of multistate diffusion processes, as described in detail in our previous study[3].

##### 5.2 *Extraction and summary of additional kinetics and thermodynamic parameters.*

The Zur protein in a cell dynamically interconvert between the three states (FD, NB, and TB) at a timescale of tens to hundreds of ms (i.e., the apparent unbinding rate constant from tight-binding sites, $k_{-1}$, and the non-specific sites, $k_{-2}$, are on the order of $10^0$ and $10^1$ $s^{-1}$, and the corresponding binding rates are on the same order as well[2]), much faster than the protein lifetime in the cell and our total imaging time (~30 min to 1 hour), during which the cellular protein concentration largely remains constant. We can therefore assume a quasi-equilibrium condition between these different states. By analyzing the relative populations of the three states, we can extract additional kinetic and thermodynamic parameters (Extended Data Table 1), for example, the binding rate constant ($k_1$), the binding affinity ($K_{\mathrm{D1}} = k_{\mathrm{o}}^{\mathrm{off}}/k_1$) (Fig. 2c), etc., as we showed previously[2].

Using the quasi-equilibrium approximation between the TB, NB and FD states, we have the following relationships between the ratios of $[\mathrm{PD}]_{\mathrm{TB}}$ and $[\mathrm{P}]_{\mathrm{FD}}$ ( Eq. S16), $[\mathrm{PD}]_{\mathrm{TB}}$ and $[\mathrm{PD}]_{\mathrm{NB}}$ (Eq. S17), and $[\mathrm{PD}]_{\mathrm{NB}}$ and $[\mathrm{PD}]_{\mathrm{FD}}$ (Eq. S18) as we derived previously[2], where $[\mathrm{PD}]_{\mathrm{TB}}$, $[\mathrm{P}]_{\mathrm{FD}}$, and $[\mathrm{PD}]_{\mathrm{NB}}$ are cellular protein concentrations of the TB, FD and NB states, which can be calculated from the fractional populations of $A_{\mathrm{TB}}$, $A_{\mathrm{FD}}$, and $A_{\mathrm{NB}}$ and the total cellular protein concentration.

$$\frac{[\mathrm{PD}]_{\mathrm{TB}}}{[\mathrm{P}]_{\mathrm{FD}}} = \frac{k_1[D_0]_{\mathrm{TB}}}{k_{-1}}\frac{\partial \ln F_{\mathrm{TB}\leftarrow\mathrm{FD}}(x_{\mathrm{TB}\leftarrow\mathrm{FD}})}{\partial x_{\mathrm{TB}\leftarrow\mathrm{FD}}} \quad \text{Eq. S16}$$

$$\frac{[\mathrm{PD}]_{\mathrm{NB}}}{[\mathrm{P}]_{\mathrm{FD}}} = \frac{[D_0]_{\mathrm{NB}}}{K_{D2} + [\mathrm{P}]_{\mathrm{FD}}} \quad \text{Eq. S17}$$

$$\frac{[\mathrm{PD}]_{\mathrm{TB}}}{[\mathrm{PD}]_{\mathrm{NB}}} = \frac{k_3[D_0]_{\mathrm{TB}}}{k_{-3}\left([D_0]_{\mathrm{NB}} - [\mathrm{PD}]_{\mathrm{NB}}\right)}\frac{\partial \ln F_{\mathrm{TB}\leftarrow\mathrm{NB}}(x_{\mathrm{TB}\leftarrow\mathrm{NB}})}{\partial x_{\mathrm{TB}\leftarrow\mathrm{NB}}} \quad \text{Eq. S18}$$

The expressions for: $x_{\text{TB}\leftarrow\text{FD}} = \frac{k_1[\text{P}]_{\text{FD}}}{k_{-1}}$, $x_{\text{TB}\leftarrow\text{NB}} = \frac{k_3[\text{PD}]_{\text{NB}}}{k_{-3}\,([D_0]_{\text{NB}}-[\text{PD}]_{\text{NB}})}$, $F_{\text{TB}\leftarrow\text{FD}}(x_{\text{TB}\leftarrow\text{FD}}) \equiv 1 + x_{\text{TB}\leftarrow\text{FD}} + x^2_{\text{TB}\leftarrow\text{FD}} + \cdots + x^{n0}_{\text{TB}\leftarrow\text{FD}} = \sum_{i=0}^{n0} x^i_{\text{TB}\leftarrow\text{FD}}$ and, $F_{\text{TB}\leftarrow\text{NB}}(x_{\text{TB}\leftarrow\text{NB}}) \equiv 1 + x_{\text{TB}\leftarrow\text{NB}} + x^2_{\text{TB}\leftarrow\text{NB}} + \cdots + x^{n0}_{\text{TB}\leftarrow\text{NB}} = \sum_{i=0}^{n0} x^i_{\text{TB}\leftarrow\text{NB}}$, were derived previously[2], where $k_i$ and $k_{-i}$ (i = 1, 2, 3) are the interconversion rate constants between the three diffusive states (Fig. 2c) and $[D_0]_{\text{TB}}$, and $[D_0]_{\text{NB}}$ are the cellular concentration of the TB and NB sites, respectively.

Using predetermined values of $k_0^{\text{off}}$, $k_{\text{f}}$, $k_{\text{r}}$, and $K_m$ from the analysis of $k_{-1}$ (Fig. 2 and Fig. 5) and a fixed value for the oligomerization number, $\text{n}_0$ (assumed to be 5 here), which is the maximum number of oligomers at a TB site (note we showed previously that for the range of values where $n_0 > 3$, the extracted kinetic parameters are not influenced significantly and approach asymptotic values within error bars[2]), we can fit $[\text{PD}]_{\text{TB}}$ / $[\text{P}]_{\text{FD}}$ vs $[\text{P}]_{\text{FD}}$ with equation Eq. S16 (Supplementary Fig. 13a) and obtain the binding rate constant ($k_1$) and the binding affinity $K_{\text{D1}}$ (= $k_0^{\text{off}}/k_1$). Further, by fitting $[\text{PD}]_{\text{NB}}$ / $[\text{P}]_{\text{FD}}$ vs $[\text{P}]_{\text{FD}}$ and $[\text{PD}]_{\text{TB}}$ / $[\text{PD}]_{\text{NB}}$ vs $[\text{PD}]_{\text{NB}}$ with equation Eq. S17 and Eq. S18 (Supplementary Fig. 13b and c), we can obtain $K_{\text{D2}}$ (= $k_{-2}/k_2$), $[D_0]_{\text{NB}}$, $K_{\text{D3}}$ (= $k_{-3}/k_3$), and $[D_0]_{\text{TB}}$, respectively (Extended Data Table 1)[2].

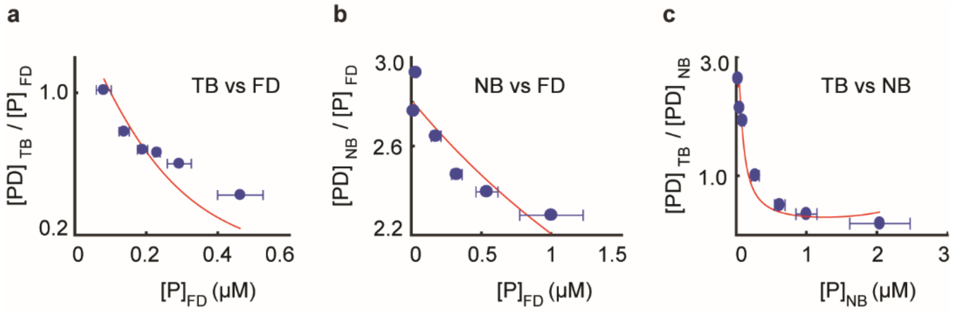


**Supplementary Fig. 13 | Example of relative population analysis for $\text{Zur}^{\text{mE}}_{\text{C88S}}$.** **a**, $[\text{PD}]_{\text{TB}}$ / $[\text{P}]_{\text{FD}}$ vs $[\text{P}]_{\text{FD}}$, **b**, $[\text{PD}]_{\text{NB}}$ / $[\text{P}]_{\text{FD}}$ vs $[\text{P}]_{\text{FD}}$, and **c**, $[\text{PD}]_{\text{TB}}$ / $[\text{PD}]_{\text{NB}}$ vs $[\text{P}]_{\text{NB}}$. The red lines are fits. All error bars are s.d.

#### 6 Protein labeling design for single-molecule FRET measurements *in vitro*

##### 6.1 *Selecting locations for fluorescent probe location on Zur based on Zur structure, and mutations to make ZntR permanent apo*

For the dimeric *E. coli* Zur, each monomer contains nine cysteines. Five of these nine cysteines are essential for Zur's Zn-binding properties at two Zn-binding sites: one cysteine at the regulatory Zn-binding site (C88, Extended Data Fig. 3b) and the other four at the structural Zn-binding site (C103, C106, C143, C146; Extended Data Fig. 3c)[9,51]. All these five essential cysteines can be protected by Zn from fluorophore labeling. The left four cysteines are non-conserved and are all exposed to the surface (C17, C113, C152, C158) on the basis of Zur's crystal structure[9]. In one variant, we used the natural C113 as the Cy5 labeling site, which is far away from Zur's DNA binding domain (Extended Data Fig. 3a), and thus labeling at this position is expected to not interfere with Zur's DNA binding; the other three non-conserved cysteines (C17, C152, C158) were mutated to serine; we, refer this labeled Zur variant (Cy5 at C113) as $\text{Zur}^{\text{Cy5}}$ in this study unless otherwise noted. Alternatively, when we used C158 as a labeling site, other three cysteine residues (C17, C113, C152) were mutated to serines; we refer to this labeled variant as $\text{Zur}^{\text{Cy5-C158}}$.

In wild-type *E. coli* ZntR, two $Zn^{2+}$ ions are bound to the protein through C114, C115, H119, C124, and C79 at each of the two Zn-binding sites in the dimeric protein (Supplementary Fig. 14)[11]. Mutating either of these residues deactivate ZntR[10]. We mutated C115 to serine, which was shown to make ZntR permanently apo and a constitutive repressor for Zn efflux genes in *E. coli*[10].

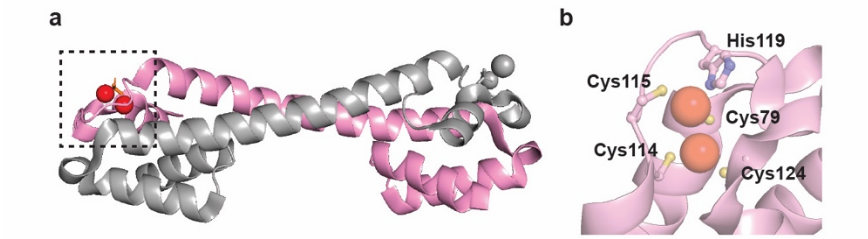


**Supplementary Fig. 14 | a,** Crystal structural of the homo-dimeric *E. coli* ZntR (PDB: 1Q08)[52], where the two ZntR monomers are colored pink and gray, and two zinc ions are shown in red spheres at one site. **b,** Zoomed-in image of zinc-binding residues at one site.

##### 6.2 *Prediction of $E_{FRET}$ values based on Zur-DNA complex structure*

For a single Zur$^{Cy5}$ dimer bound to 22-bp truncated DNA$^{Cy3}$, there are one FRET donor and one acceptor, forming a single FRET donor – acceptor system. In this system, $E_{\mathrm{FRET}}$ value can be estimated using Eq. S19[53], where $R_0$ is the Förster radius (5.4 nm for Cy3-Cy5 pair[54]), $R$ is the distance measured between two anchoring positions of the dyes, and $\tau_D$ is the lifetime of the donor Cy3.

$$E = \frac{k_{D\to A}}{k_{D\to A}+\frac{1}{\tau_D}},\ k_{D\to A} = \frac{1}{\tau_D}\left(\frac{R_0}{R}\right)^6 \qquad \text{Eq. S19}$$

For Zur$^{Cy5}$, Cy5 is attached to the surface-exposed C113 of one monomer of the dimeric Zur. This labeling scheme makes a singly labeled Zur homodimer asymmetric, giving rise to two different binding orientations on DNA. Based on the crystal structure of holo Zur-DNA complex and our label position (Extended Data Fig. 4a-c)[9], the Cy3−Cy5 anchor-to-anchor distances in a Zur$^{Cy5}$−DNA complex are about (A) 49 Å and (B) 56 Å for the two binding orientations (Extended Data Fig. 4b and c) The corresponding $E_{\mathrm{FRET}}$ values are ~0.64 and ~0.45, respectively, and should be resolvable in an experimental $E_{\mathrm{FRET}}$ histogram (e.g., Fig. 3d and e). Experimentally, we observed two $E_{\mathrm{FRET}}$ states at ~0.65 and ~0.44 (Fig. 3d and e), in agreement with these predictions.

Similarly, a single dimer Zur$^{Cy5\text{-}C158}$ binding to the truncated 22-bp DNA$^{Cy3}$ should also give rise to two different binding orientations with distances of (C) 71 Å and (D) 34 Å (Extended Data Fig. 4b and c), and the corresponding $E_{\mathrm{FRET}}$ values should be ~0.16 and ~0.94, respectively. The distance is measured between dye labeling anchor to C152 of Zur due to structurally unresolved C158. Thus, the observed value can be unmatched from the corresponding value but is expected to be farther away from Cy5 compared to C113 Experimentally, we observed two $E_{\mathrm{FRET}}$ states at ~0.41 and ~0.77 (Fig. 3f), in agreement with these predictions. All expected $E_{\mathrm{FRET}}$ values calculated by Eq. S19 are summarized in the 'Expected $E_{\mathrm{FRET}}$ (1)' column in Supplementary Table 7.

Alternatively, the FRET value can be predicted on the basis of our experimental calibration of observed $E_{\mathrm{FRET}}$ vs. Cy3-Cy5 distances, where Cy3-Cy5 are anchored on DNA structures with known inter-distances between the anchor points[16,55,56]. Experimental data were fitted empirically using Eq. S20, where $R_0$ is the Förster radius (5.4 nm for Cy3-Cy5[54]) and $R$ is the distance measured between two phosphate backbone atoms corresponding to the anchoring position of the dyes.

$$E = \frac{k_{D\to A}}{k_{D\to A}+\frac{1}{\tau_D}},\ k_{D\to A} = \frac{1}{\tau_D}\left(\frac{R_0\beta}{R+\alpha}\right)^6 \qquad \text{Eq. S20}$$

Here, $\alpha$ is the correction parameter for the additional distance due to the dye linker length and $\beta$ is the correction parameter for deviation in $R_0$ because of the linker flexibility and the orientation of the dyes[57].

Based on the fitting, $\alpha$ = 5.3 ± 0.6 nm and $\beta$ = 1.9 ± 0.1[16]. Note the $E_{\mathrm{FRET}}$ value is not corrected for the relative fluorescence detection efficiencies and quantum yields of the dyes[57,58]. Thus, the numerical values including $\alpha$ and $\beta$ should be treated as empirical fitting parameters, not be interpreted literally despite their physical connections. This empirical calibration curve provides a direct correlation between an experimental observable (apparent $E_{\mathrm{FRET}}$) and a distance quantity (anchor-to-anchor distance) that can be independently determined reliably using structural modeling. All expected $E_{\mathrm{FRET}}$ values calculated by Eq. S20 are summarized in the 'Expected $E_{\mathrm{FRET}}$ (2)' column in Supplementary Table 7.

For $\mathrm{Zur^{Cy5}}$ bound to 31-bp truncated $\mathrm{DNA^{Cy3}}$, which encodes the complete two-dyad Zur binding box, two homodimeric Zur can bind to the DNA simultaneously, which makes 1 FRET donor – 2 FRET acceptor system. Here, $E_{\mathrm{FRET}}$ value can be calculated using Eq. S21[59].

$$E = \frac{k_{D\to A1} + k_{D\to A2}}{k_{D\to A1} + k_{D\to A2} + \frac{1}{\tau_D}},\ k_{D\to A1} = \frac{1}{\tau_D}\left(\frac{R_0}{R_1}\right)^6,\ k_{D\to A2} = \frac{1}{\tau_D}\left(\frac{R_0}{R_2}\right)^6 \qquad \text{Eq. S21}$$

For the two Zur dimers, one is at the dyad sequence proximal to the Cy3 label on DNA (Dimer 1), the other at the dyad distal to the Cy3 label (Dimer 2). Both dimers, each carrying a Cy5 label that breaks the dimer symmetry, can each give rise to two different binding orientations. The two possible orientations for the proximal dimer are shown in Extended Data Fig. 4a. For the distal dimer, its two orientations give a Cy5-Cy3 distance of (E) 53 Å and (F) 68 Å, respectively, as shown in Extended Data Fig. 4d. Two Zur dimer binding with two binding orientations each make four combinations of two-dimer-bound form (Extended Data Fig. 4d-h).

Again, the FRET values for two FRET acceptor system can be predicted on the basis of our experimental calibration of observed $E_{\mathrm{FRET}}$ vs. Cy3-Cy5 distances as well as one FRET acceptor system. $R_0$ and $R_1$, $R_2$ are corrected with $\beta$ and $\alpha$, respectively based on Eq. S20. The expected FRET values and the experimentally observed FRET values are summarized in Supplementary Table 7; they show good agreements, especially in the ordering of $E_{\mathrm{FRET}}$ values among the different configurations.

**Supplementary Table 7 |** Expected $E_{\mathrm{FRET}}$ values calculated from structural model

| | Cy5 location | distance from Cy3 (Å) | Expected $E_{\mathrm{FRET}}$ (1) | Expected $E_{\mathrm{FRET}}$ (2) | Observed $E_{\mathrm{FRET}}$ |
|---|---|---|---|---|---|
| 1 FRET acceptor | Dimer1-1 (C113) | 49 | 0.64 | 0.51 | 0.65 |
| | Dimer1-2 (C113) | 56 | 0.45 | 0.41 | 0.44 |
| | Dimer2-1 (C113) | 53 | 0.53 | 0.45 | 0.43 |
| | Dimer2-2 (C113) | 68 | 0.20 | 0.27 | 0.27 |
| | Dimer1-1 (~C158) | 71 | 0.16 | 0.24 | 0.41 |
| | Dimer1-2 (~C158) | 34 | 0.94 | 0.73 | 0.77 |
| 2 FRET acceptor | Dimer1-1, Dimer 2-1 (C113) | 49, 53 | 0.74 | 0.65 | 0.80 |
| | Dimer1-1, Dimer 2-2 (C113) | 49, 68 | 0.67 | 0.58 | 0.68 |
| | Dimer1-2, Dimer 2-1 (C113) | 56, 53 | 0.66 | 0.60 | 0.68 |
| | Dimer1-2, Dimer 2-2 (C113) | 56, 68 | 0.51 | 0.52 | 0.47 |

#### 7 Procedures for Gaussian fitting to extract $E_{\mathrm{FRET}}$ values from the $E_{\mathrm{FRET}}$ histograms

As we described in Fig. 3d-e, in the main text, the binding of $\mathrm{Zur^{Cy5}_{Zn,\,D49A}}$ and $\mathrm{Zur^{Cy5}_{Zn}}$ onto the 22-bp truncated $\mathrm{DNA^{Cy3}}$ are expected to show the same $E_{\mathrm{FRET}}$ states because there is only one binding dyad site on the truncated DNA for Zur and the D49A mutation that removes the key inter-dimer salt-bridge interaction would not cause a significant difference between $Zur^{Cy5}_{Zn,\ D49A}$ and $Zur^{Cy5}_{Zn}$ (Fig. 3a, bottom). We extracted the $E_{FRET}$ values for the two binding orientations of $Zur^{Cy5}_{Zn,\ D49A}$ on the truncated 22-bp $DNA^{Cy3}$ via two-dimensional histogram analysis of lower vs. higher $E_{FRET}$ values observed in the $E_{FRET}$ trajectories (Fig. 3c; Supplementary Fig. 20a), which are 0.43 ± 0.13 and 0.69 ± 0.13. With the same two-dimensional histogram analysis for $Zur^{Cy5}_{Zn}$ on the truncated $DNA^{Cy3}$, $E_{FRET}$ values for its two binding orientations are 0.41 ± 0.08 and 0.66 ± 0.08 (Supplementary Fig. 20b), which, expectedly, are within error to, and thus the same as, those for $Zur^{Cy5}_{Zn,\ D49A}$. Therefore, to further improve data fitting reliability, we subsequently fitted two data sets globally with Gaussian functions sharing the peak positions (Supplementary Fig. 15a-b). Alternatively, we can also combine two data sets to have better statistics for fitting (Supplementary Fig. 15c). Both analyses gave the same three $E_{FRET}$ values, ~0.03, ~0.44, and ~0.65, for the free DNA state and the two binding orientations of the protein on the truncated $DNA^{Cy3}$.

We used these values to resolve $E_{FRET}$ states for $Zur^{Cy5}_{Zn,\ D49A}$ bindings on the 31-bp $DNA^{Cy3}$, which have the complete two dyads of the Zur binding box (Fig. 3a, middle). One dyad binding site is proximal to the Cy3 label on DNA and is the same as the one in the 22-bp truncated $DNA^{Cy3}$ (Fig. 3a, middle vs. bottom); so $Zur^{Cy5}_{Zn,\ D49A}$ binding to this proximal site is expected to show the same two $E_{FRET}$ states as those from binding on the truncated $DNA^{Cy3}$. To resolve the rest $E_{FRET}$ states, we fitted the $E_{FRET}$ histogram with five Gaussian functions including (Supplementary Fig. 15d): a peak for the free DNA state near zero $E_{FRET}$ value; two peaks whose positions and the amplitude ratio are taken from the interaction with the truncated $DNA^{Cy3}$; and two more peaks to account for the two orientations of $Zur^{Cy5}_{Zn,\ D49A}$ binding to the distal dyad site on the 31-bp $DNA^{Cy3}$ whose positions and amplitudes are floated; the widths of the four DNA bound peaks are shared. The fitted results gave the additional two $E_{FRET}$ values at ~0.43 and ~0.27, respectively (Supplementary Fig. 15d). All these $E_{FRET}$ values also agree with predictions from the $Zur_{Zn}$-DNA complex structure (Supplementary Table 7).

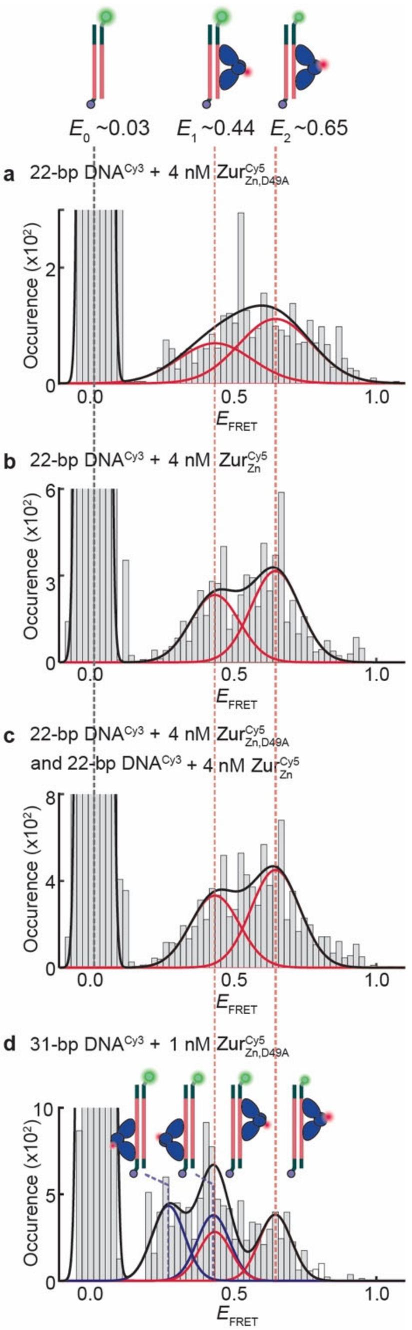


**Supplementary Fig. 15 | a,** Histograms of $E_{\mathrm{FRET}}$ trajectories of an immobilized 22-bp truncated DNA$^{\mathrm{Cy3}}$ interacting with $\mathrm{Zur}^{\mathrm{Cy5}}_{\mathrm{Zn,D49A}}$ (Cy5 at C113) (4 nM). **b,** Same as (a), but with $\mathrm{Zur}^{\mathrm{Cy5}}_{\mathrm{Zn}}$ (4 nM). Two histograms are globally fitted with Gaussian functions sharing the peak positions. **c,** Histograms of $E_{\mathrm{FRET}}$ trajectories of combined data of (a) and (b) and fitted with Gaussian functions. **d,** Same as (a), but with 31-bp DNA and 1 nM of $\mathrm{Zur}^{\mathrm{Cy5}}_{\mathrm{Zn,D49A}}$; red and blue lines: Gaussian resolved fits; black lines: overall fits. Red dashed lines indicate two peaks that are assigned for Zur bindings on the proximal dyad to Cy3 with two orientations. (d) is the same figure as Fig. 4a in the main text. Cartoons show free DNA and DNA-bound Zur in two binding orientations. The FRET donor (green sphere) and acceptor (red sphere) are drawn on DNA and Zur at their approximate locations. All histograms are compiled from >300 $E_{\mathrm{FRET}}$ trajectories; bin size = 0.02.

#### 8 ZntR$_{\mathrm{apo}}$ preferentially disrupts Zur$_{\mathrm{Zn}}$ binding at the dyad proximal to the Cy3 labeling position on DNA

When $Zur_{Zn}^{Cy5}$ interacts with 31-bp $DNA^{Cy3}$, a single $Zur_{Zn}^{Cy5}$ dimer can bind to either of the two dyads of Zur box on DNA and maximally two $Zur_{Zn}^{Cy5}$ dimers can bind to the DNA simultaneously.

At a lower concentration of $Zur_{Zn}^{Cy5}$ (e.g., 1 and 2 nM, Extended Data Fig. 6d-e), $E_3$ peak at ~0.2 is observed in $E_{FRET}$ histogram, indicating that one-dimer-bound form occurs, as $E_3$ is characteristic of the one-dimer bound form (Extended Data Fig. 6a). Meanwhile, when $Zur_{Zn}^{Cy5}$ concentration is increased to 4 nM (Fig. 4c; Extended Data Fig. 6f), $E_3$ peak is no longer observed, reflecting that $Zur_{Zn}^{Cy5}$ dominantly occupy both dyad recognition sites. For all $Zur_{Zn}^{Cy5}$ concentrations, upon introducing $ZntR_{apo}$, $E_{FRET}$ histogram shows a significant $E_3$ peak (~0.2) (Extended Data Fig. 6g-l), while $E_7$ peak (at ~0.8, which is characteristic of two-dimer bound form) almost disappears, indicating that $ZntR_{apo}$ disrupts $Zur_{Zn}$ interactions with DNA, leading to the dominance of one-dimer bound form.

Interestingly, among the two dyads that $Zur_{Zn}^{Cy5}$ binds, they are not equally populated in the presence of $ZntR_{apo}$, There are much higher population at a lower $E_{FRET}$ value ($E_3$), which corresponds to $Zur_{Zn}^{Cy5}$ binding at the dyad site distal to Cy3, than at a higher $E_{FRET}$ value ($E_2$), which corresponds to $Zur_{Zn}^{Cy5}$ binding at the dyad site proximal to Cy3 (Extended Data Fig. 6g-l). Therefore, $ZntR_{apo}$ preferentially disrupts $Zur_{Zn}^{Cy5}$ binding at the proximal dyad to the Cy3 label. From sequence analysis of potential ZntR recognition sequence at the *znuBC* promoter that the 31-bp DNA is based upon, the most probable ZntR binding site overlaps more significantly with the Zur-binding dyad proximal to the Cy3 label position (Fig. 1b). We, therefore, conclude that $ZntR_{apo}$ facilitates the unbinding of incumbent Zur through recognizing the most probably binding sequences.

#### 9 A through-DNA mechanism for Zur-DNA-$ZntR_{apo}$ interactions and kinetic derivations

Our previous single-molecule tracking studies of single cells showed that the apparent unbinding rate constant $k_{-1}$ of Zur from its tight-binding sites on DNA follows a biphasic, impeded-followed-by-facilitated unbinding behavior: it initially decreases with increasing cellular Zur concentration up to ~100 nM, reaching a minimum, and then increases at higher Zur concentrations[2]. The impeded unbinding results from Zur oligomerization at its tight-binding site on DNA, in which the salt-bridge interactions between Zur dimers contribute to its oligomerization, which in turn stabilizes Zur on DNA and slows down its unbinding kinetics (Supplementary Fig. 16, Step 4).

When the cellular Zur concentration further increases, the facilitated unbinding pathways becomes more competitive, in which a freely diffusing Zur in the cytoplasm can bind partially to a recognition site occupied by an incumbent Zur to form a ternary protein-DNA complex i (Supplementary Fig. 16, Step 1); this ternary complex is made possible by the bivalent interactions between the homodimeric protein and the dyad-symmetric recognition sequence, in which each of the two dimeric proteins binds to half of the dyad sequence on the DNA. The unstable nature of this ternary complex subsequently leads to either the falling-off of both proteins from DNA, a so-called assisted dissociation pathway (Supplementary Fig. 16, Step 2), or a direct substitution of the incumbent protein by the incoming protein (Supplementary Fig. 16, Step 3). Both these pathways lead to an increase in the unbinding rate of the incumbent protein when the concentration of the protein in the cell increases, giving rise to the facilitated unbinding behavior.

Besides the Fur-family metalloregulator Zur, we also discovered the facilitated unbinding for the MerR-family metalloregulators CueR and ZntR in cells[3], which also are dimeric proteins recognizing dyad symmetric sequences on DNA. For CueR, we further characterized this facilitated unbinding using *in vitro* single-molecule FRET experiments, which clearly showed the assisted dissociation and direct substitution pathways[13]. Such facilitated unbinding was also observed *in vitro* for a number of other DNA-binding proteins, including a sequence-nonspecific DNA-binding protein (e.g., nucleoid-associated proteins, NAP), a sequence-neutral single-stranded DNA-binding protein (e.g., Replication protein A, RPA), and DNA polymerases[13,60–77]. Moreover, facilitated dissociation was also observed between heterotypic proteins on DNA (e.g., the unbinding of a human linker histone H1.0 (H1) bound to a nucleosome facilitated by a histone chaperone prothymosin α and the unbinding of NF-κB facilitated by its specific inhibitor IκBα), where the 2nd protein seems to only interact with the DNA-bound protein and not the DNA[78–80]. Facilitated unbinding has also been investigated theoretically[81–84].

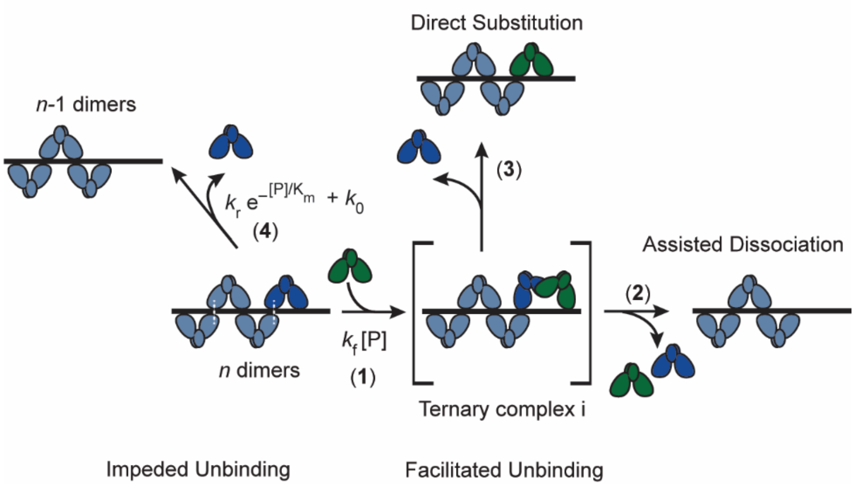


**Supplementary Fig. 16 |** Schematics of impeded unbinding (left, Step 4) for Zur and facilitated unbinding for Zur and ZntR/CueR (right, Steps 1, 2 and 3). Freely diffusing proteins are shown in green. Incumbent proteins on DNA are shown in blue. [P]: concentration of Zur or ZntR protein.

For the impeded-followed-by-facilitated unbinding behavior of Zur, we have previously derived the following equation to quantitatively describe Zur's apparent 1st-order unbinding rate constant $k_{-1}$ from a tight-binding site as a function of its cellular concentration of freely diffusing component[2]:

$$k_{-1} = k_0^{\mathrm{off}} + k_{\mathrm{r}}\,(\mathrm{e}^{\frac{-[\mathrm{Zur}]}{K_{\mathrm{m}}}} - 1) + k_{\mathrm{f}}\,[\mathrm{Zur}] \qquad \text{Eq. S22}$$

where $k_{-1}$ is the apparent 1st-order apparent unbinding rate constant; $k_{\mathrm{f}}$ is a 2nd-order facilitated unbinding rate constant; $k_0^{\mathrm{off}} = k_0 + k_{\mathrm{r}}$, where $k_0$ is the intrinsic protein unbinding rate constant; $k_{\mathrm{r}}$ is a 1st-order impeded unbinding rate constant; and $K_{\mathrm{m}}$, is the effectively affinity constant (in concentration units) of Zur oligomerization on DNA.

##### 9.1 *Empirical kinetic equation for ZntR$_{apo}$-induced enhancement of Zur's facilitated unbinding and diminishment of Zur's impeded unbinding*

In this study we have uncovered the effects of ZntR$_{\mathrm{apo}}$ on Zur unbinding and our results show that $k_{\mathrm{r}}$ and $k_{\mathrm{f}}$ of Zur both depend on [ZntR$_{\mathrm{apo}}$] linearly with a positive slope and intercept (Fig. 2f, and Fig. 5c and e). Thus, we can replace $k_{\mathrm{r}}$ and $k_{\mathrm{f}}$ in Eq. S22 as:

$$k_{\mathrm{r}} = k_{\mathrm{r2}}[\mathrm{ZntR_{apo}}] + k_{\mathrm{r1}} \qquad \text{Eq. S23}$$

$$k_{\mathrm{f}} = k_{\mathrm{f2}}[\mathrm{ZntR_{apo}}] + k_{\mathrm{f1}} \qquad \text{Eq. S24}$$

where $k_{\mathrm{r1}}$, $k_{\mathrm{r2}}$, $k_{\mathrm{f1}}$, and $k_{\mathrm{f2}}$ are empirical constants. Eq. S22 becomes:

$$k_{-1} = k_0 + (k_{r2}[\text{ZntR}_{\text{apo}}] + k_{r1})(e^{\frac{-[\text{Zur}]}{K_m}}) + (k_{f2}[\text{ZntR}_{\text{apo}}] + k_{f1})[\text{Zur}] \quad \text{Eq. S25}$$

Eq. S25 empirically describes the effective unbinding rate constant $k_{-1}$ of Zur as a function of [Zur] and [$\text{ZntR}_{\text{apo}}$] in the cell. Below we will use the mechanistic model from Fig. 5F to derive this relationship between $k_{-1}$ and [Zur] and [$\text{ZntR}_{\text{apo}}$].

##### 9.2 *Kinetic derivation and justification of the mechanistic model for ZntR$_{apo}$-dependent Zur unbinding from DNA*

On the basis of the observed [Zur] and [$\text{ZntR}_{\text{apo}}$] dependence of Zur's unbinding kinetics, which are empirically described by Eq. S25 above, we proposed the mechanistic scheme of Zur unbinding, in which $\text{ZntR}_{\text{apo}}$ can act directly on DNA-bound Zur (Fig. 5F; Supplementary Fig. 17). Starting from oligomeric Zur dimers bound at a tight-binding site (i.e., *n* dimers), Zur can unbind spontaneously ($k_0$ component in Step 4) and its unbinding can also be impeded by its oligomerization on DNA due to the extra stability from inter-dimer interactions (the $k_{r1}\, e^{-\frac{[\text{Zur}]}{K_m}}$ component in Step 4), as we previously formulated[2]. In the presence of free Zur and $\text{ZntR}_{\text{apo}}$ in the surrounding, this mechanism can proceed by the formation of one ternary complex intermediate, i (Step 1), and two transition states, TS1 (a quaternary complex) and TS2 (a ternary complex ii). The formation of ternary complex intermediate i, using an incoming freely diffusing cytoplasmic Zur (Step 1), can lead to Zur's assisted dissociation and direct substitution (Steps 2 and 3), accounting for the facilitated unbinding of Zur that we previously discovered[2]. The formation of the heteromeric TS1 quaternary complex, using an incoming free $\text{ZntR}_{\text{apo}}$, accounts for $\text{ZntR}_{\text{apo}}$-enhanced facilitated unbinding of Zur (Step 5), while the formation of the heteromeric TS2 ternary complex ii, also using an incoming free $\text{ZntR}_{\text{apo}}$ (Step 6), accounts for $\text{ZntR}_{\text{apo}}$ dependent diminishment of Zur's impeded unbinding process.

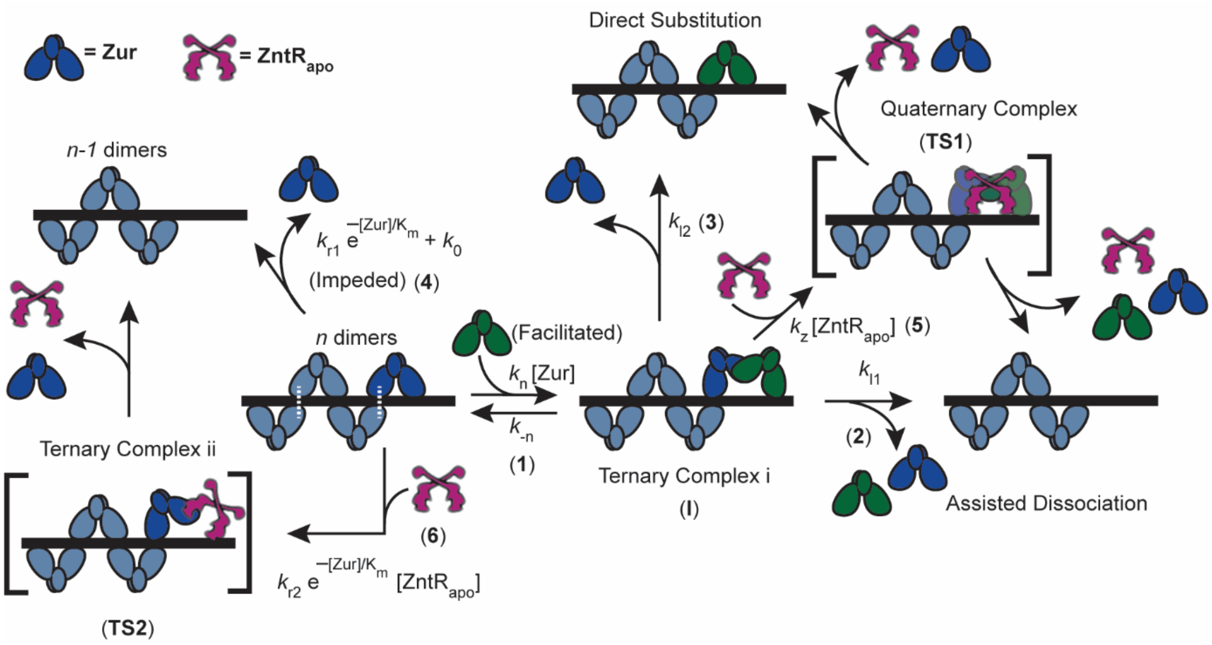


**Supplementary Fig. 17 | A "through-DNA" mechanistic model for $\text{ZntR}_{\text{apo}}$-dependent Zur unbinding kinetics.** Starting with *n* oligomerized Zur dimers at a tight-binding site on DNA, the unbinding of an incumbent Zur protein (dark blue) can be facilitated by a freely diffusing Zur (dark green) through the formation of a ternary complex i (Step 1), leading to assisted dissociation (Step 2) or direct substitution (Step 3); this facilitated unbinding of Zur can be enhanced by $\text{ZntR}_{\text{apo}}$ through the formation of a heteromeric quaternary complex (Step 5). The oligomer-induced impedance of Zur unbinding (Step 4) can be weakened by $\text{ZntR}_{\text{apo}}$ through the formation of a heteromeric ternary complex ii (Step 6), leading to faster Zur unbinding as well. White dashed lines denote salt bridge interactions between Zur dimers. The associated rate constants, *k*'s, are denoted on the respective kinetic steps in the mechanism.

The mechanism in Supplementary Fig. 17 can be separated into the following kinetic processes with their associated rate constants:

<u>Impeded and spontaneous unbinding pathways of Zur and the dependence on [ZntR$_{\mathrm{apo}}$]</u>

$$\mathrm{Zur_nD} \xrightarrow{k_{\mathrm{r1}}\, \mathrm{e}^{-\frac{[\mathrm{Zur}]}{K_m}} + k_0} \mathrm{Product}\ \ \text{(Step 4)} \qquad \text{Eq. S26}$$

$$\mathrm{Zur_nD} + \mathrm{ZntR_{apo}} \xrightarrow{k_{\mathrm{r2}}\, \mathrm{e}^{-\frac{[\mathrm{Zur}]}{K_m}}} \mathrm{Product}\ \ \text{(Step 6)} \qquad \text{Eq. S27}$$

where Zur$_n$D represents *n* Zur dimers bound at a tight-binding site on DNA (i.e., D). In step 4 (Eq. S26), the rate constant is $k_{\mathrm{r1}}\, \mathrm{e}^{-\frac{[\mathrm{Zur}]}{K_m}} + k_0$, where $K_{\mathrm{m}}$ is the effective dissociation constant of the protein oligomer, as we derived in our previous work to account for the impeded unbinding of Zur from DNA[2].

<u>Facilitated unbinding pathways of Zur and is enhancement by [ZntR$_{\mathrm{apo}}$]</u>

$$\mathrm{Zur_nD} + \mathrm{Zur} \underset{k_{-\mathrm{n}}}{\overset{k_{\mathrm{n}}}{\rightleftarrows}} \mathrm{I}\quad \text{(Step 1)} \qquad \text{Eq. S28}$$

$$\mathrm{I} \xrightarrow{k_{\mathrm{l1}}} \mathrm{Product}\ \ \text{(Assisted Dissociation) (Step 2)} \qquad \text{Eq. S29}$$

$$\mathrm{I} \xrightarrow{k_{\mathrm{l2}}} \mathrm{Product}\ \ \text{(Direct Substitution) (Step 3)} \qquad \text{Eq. S30}$$

Eq. S29 and Eq. S30 can be combinedly written as

$$\mathrm{I} \xrightarrow{k_{\mathrm{l}}} \mathrm{Product} \qquad \text{Eq. S31}$$

where, $k_{\mathrm{l}} = k_{\mathrm{l1}} + k_{\mathrm{l2}}$. For the ZntR$_{\mathrm{apo}}$-enhanced facilitated unbinding:

$$\mathrm{I} + \mathrm{ZntR_{apo}} \xrightarrow{k\mathrm{z}} \mathrm{Product}\ \text{(Step 5)} \qquad \text{Eq. S32}$$

Taking into account Eq. S26 – Eq. S32 , we can write the following rate equations:

$$\frac{\mathrm{d[I]}}{\mathrm{d}t} = k_{\mathrm{n}}[\mathrm{Zur_nD}][\mathrm{Zur}] - k_{-\mathrm{n}}[\mathrm{I}] - k_{\mathrm{l}}[\mathrm{I}] - k_{\mathrm{z}}[\mathrm{I}][\mathrm{ZntR_{apo}}] \qquad \text{Eq. S33}$$

$$\frac{\mathrm{d[Products]}}{\mathrm{dt}} = k_{\mathrm{r1}}\, \mathrm{e}^{-\frac{[\mathrm{Zur}]}{K_m}}[\mathrm{Zur_nD}] + k_{\mathrm{r2}}\, \mathrm{e}^{-\frac{[\mathrm{Zur}]}{K_m}}[\mathrm{Zur_nD}]\left[\mathrm{ZntR_{apo}}\right] + k_0\,[\mathrm{Zur_nD}] + k_{\mathrm{l}}\,[\mathrm{I}] + k_{\mathrm{z}}\,[\mathrm{I}]\left[\mathrm{ZntR_{apo}}\right] \qquad \text{Eq. S34}$$

Here "Products" represent all product species that resulted from the Zur unbinding from tight-binding sites via all possible pathways. Assuming steady state approximation to Eq. S33:

$$\frac{\mathrm{d[I]}}{\mathrm{dt}} = 0$$

We then have:

$$k_{\mathrm{n}}[\mathrm{Zur_nD}][\mathrm{Zur}] - k_{-\mathrm{n}}[\mathrm{I}] - k_{\mathrm{l}}[\mathrm{I}] - k_{\mathrm{z}}[\mathrm{I}][\mathrm{ZntR_{apo}}] = 0$$

$$k_{-\mathrm{n}}[\mathrm{I}] + k_{\mathrm{l}}[\mathrm{I}] + k_{\mathrm{z}}[\mathrm{I}][\mathrm{ZntR_{apo}}] = k_{\mathrm{n}}[\mathrm{Zur_nD}][\mathrm{Zur}]$$

$$\therefore [\mathrm{I}] = \frac{k_{\mathrm{n}}[\mathrm{Zur_nD}][\mathrm{Zur}]}{k_{-\mathrm{n}} + k_{\mathrm{l}} + k_{\mathrm{z}}[\mathrm{ZntR_{apo}}]}$$

Replacing the expression for [I] in Eq. S34

$$\frac{\mathrm{d[Products]}}{\mathrm{dt}} = k_{\mathrm{r1}}\, \mathrm{e}^{-\frac{[\mathrm{Zur}]}{K_m}}[\mathrm{Zur_nD}] + k_{\mathrm{r2}}\, \mathrm{e}^{-\frac{[\mathrm{Zur}]}{K_m}}[\mathrm{Zur_nD}][\mathrm{ZntR_{apo}}] + k_0\,[\mathrm{Zur_nD}] + k_{\mathrm{l}}\, \frac{k_{\mathrm{n}}[\mathrm{Zur_nD}][\mathrm{Zur}]}{k_{-\mathrm{n}} + k_{\mathrm{l}} + k_{\mathrm{z}}[\mathrm{ZntR_{apo}}]} + k_{\mathrm{z}}\, \frac{k_{\mathrm{n}}[\mathrm{Zur_nD}][\mathrm{Zur}]}{k_{-\mathrm{n}} + k_{\mathrm{l}} + k_{\mathrm{z}}[\mathrm{ZntR_{apo}}]}\,[\mathrm{ZntR_{apo}}] \quad \text{Eq. S35}$$

To simply, we can make the approximation that the Ternary Complex I is not a stable species and its dissociation is fast compared with the formation of the heteromeric Quaternary Complex TS1, i.e., $k_{-\mathrm{n}} >> k_{\mathrm{z}}[\mathrm{ZntR_{apo}}]$. Moreover, the direct substitution rate constant $k_{\mathrm{l2}}$ (Step 3) is effectively the same as that $k_{-\mathrm{n}}$, and thus the same approximation gives $k_{\mathrm{l2}} >> k_{\mathrm{z}}[\mathrm{ZntR_{apo}}]$ and therefore $k_{\mathrm{l}} = k_{\mathrm{l1}} + k_{\mathrm{l2}} >> k_{\mathrm{z}}[\mathrm{ZntR_{apo}}]$. Consequently, the dominator in the 4$^{\mathrm{th}}$ and 5$^{\mathrm{th}}$ term in Eq. S35 can be simplified to:

$$\frac{\mathrm{d[Products]}}{\mathrm{dt}} = k_{\mathrm{r1}}\, \mathrm{e}^{-\frac{[\mathrm{Zur}]}{K_m}}[\mathrm{Zur_nD}] + k_{\mathrm{r2}}\, \mathrm{e}^{-\frac{[\mathrm{Zur}]}{K_m}}[\mathrm{Zur_nD}][\mathrm{ZntR_{apo}}] + k_0\,[\mathrm{Zur_nD}] + k_{\mathrm{l}}\, \frac{k_{\mathrm{n}}[\mathrm{Zur_nD}]}{k_{-\mathrm{n}} + \mathrm{k_l}}[\mathrm{Zur}] + k_{\mathrm{z}}\, \frac{k_{\mathrm{n}}[\mathrm{Zur_nD}]}{k_{-\mathrm{n}} + \mathrm{k_l}}\,[\mathrm{Zur}][\mathrm{ZntR_{apo}}] \quad \text{Eq. S36}$$

Also, from the three-state model (Fig. 2c) using which we extracted the apparent unbinding rate constant $k_{-1}$, we also have the following,

$$\frac{\mathrm{d[Products]}}{\mathrm{dt}} = k_{-1}[\mathrm{Zur_nD}] \quad \text{Eq. S37}$$

Equating Eq. S36 and Eq. S37, we have the following:

$$k_{-1} = k_{\mathrm{r1}}\, \mathrm{e}^{-\frac{[\mathrm{Zur}]}{\mathrm{K}_m}} + k_{\mathrm{r2}}\, \mathrm{e}^{-\frac{[\mathrm{Zur}]}{\mathrm{K}_m}}[\mathrm{ZntR_{apo}}] + k_0 + k_{\mathrm{l}}\, \frac{k_{\mathrm{n}}}{k_{-\mathrm{n}} + k_{\mathrm{l}}}[\mathrm{Zur}] + k_{\mathrm{z}}\, \frac{k_{\mathrm{n}}}{k_{-\mathrm{n}} + k_{\mathrm{l}}}\,[\mathrm{Zur}][\mathrm{ZntR_{apo}}] \quad \text{Eq. S38}$$

Replacing in Eq. S38 by:

$$k_{\mathrm{f2}} \equiv k_z\, \frac{k_n}{k_{-n} + k_l} \qquad \text{and } k_{\mathrm{f1}} \equiv k_{\mathrm{l}}\, \frac{k_{\mathrm{n}}}{k_{-\mathrm{n}} + k_{\mathrm{l}}}$$

we can arrive at the following equation, which has the same form as the empirical Eq. S25,

$$k_{-1} = k_0 + (k_{\mathrm{r2}}[\mathrm{ZntR_{apo}}] + k_{\mathrm{r1}})(\mathrm{e}^{\frac{-[\mathrm{Zur}]}{K_{\mathrm{m}}}}) + (k_{\mathrm{f2}}[\mathrm{ZntR_{apo}}] + k_{\mathrm{f1}})\,[\mathrm{Zur}] \quad \text{Eq. S39}$$

#### 10 Within the physiological concentration range of Zur and ZntR, $\mathrm{ZntR_{apo}}$ can enhance the apparent unbinding rate constant of $\mathrm{Zur_{Zn}}$ by ~130% and that of $\mathrm{Zur_{C88S}}$ by ~50%

Under physiological expression from their chromosomal loci, the cellular concentration of Zur ranges from ~10 to ~300 nM[2] and that of ZntR ranges from ~30 to ~400 nM (Extended Data Fig. 7b). We sorted the cells within the physiological [Zur] range, and extracted the apparent Zur unbinding rate constant,

$k_{-1,}$ for different [$ZntR_{apo}$] concentration groups within the physiological [$ZntR_{apo}$] range. In this range of cellular protein concentration of Zur and ZntR, $ZntR_{apo}$ was found to enhance $k_{-1}$ of the repressor form of $Zur_{Zn}$ from ~17 to ~38 $s^{-1}$, i.e., by ~130% (Extended Data Fig. 7a, purple). In the same [$ZntR_{apo}$] range, $k_{-1}$ of the non-repressor form of $Zur_{C88S}$ increases from ~18 to ~26 $s^{-1}$, i.e., by ~50% (Extended Data Fig. 7a, magenta). The apparent less enhancement on $Zur_{C88S}$ unbinding likely results from the fact that the non-repressor form of $Zur_{C88S}$ binds DNA tightly at sites that do not share consensus with the Zur binding box and these non-consensus $Zur_{C88S}$ binding sites may not always have nearby sequences that partially overlap with the recognition sequence of ZntR.

#### 11 Additional data and figures

**a**

E coli Zur WT: MEKTTTQELL AQAEKICAQR NVRLTPQRLE VLRLMSLQDG AISAYDLLDL
Zur variant 1: MEKTTTQELL AQAEKISAQR NVRLTPQRLE VLRLMSLQDG AISAYDLLDL
Zur variant 2: MEKTTTQELL AQAEKISAQR NVRLTPQRLE VLRLMSLQDG AISAYDLLAL
Zur variant 3: MEKTTTQELL AQAEKISAQR NVRLTPQRLE VLRLMSLQDG AISAYDLLDL

E coli Zur WT: LREAEPQAKP PTVYRALDFL LEQGFVHKVE STNSYVLCHL FDQPTHTSAM
Zur variant 1: LREAEPQAKP PTVYRALDFL LEQGFVHKVE STNSYVLCHL FDQPTHTSAM
Zur variant 2: LREAEPQAKP PTVYRALDFL LEQGFVHKVE STNSYVLCHL FDQPTHTSAM
Zur variant 3: LREAEPQAKP PTVYRALDFL LEQGFVHKVE STNSYVLCHL FDQPTHTSAM

E coli Zur WT: FICDRCGAVK EECAEGVEDI MHTLAAKMGF ALRHNVIEAH GLCAACVEVE
Zur variant 1: FICDRCGAVK EECAEGVEDI MHTLAAKMGF ALRHNVIEAH GLCAACVEVE
Zur variant 2: FICDRCGAVK EECAEGVEDI MHTLAAKMGF ALRHNVIEAH GLCAACVEVE
Zur variant 3: FICDRCGAVK EESAEGVEDI MHTLAAKMGF ALRHNVIEAH GLCAACVEVE

E coli Zur WT: ACRHPEQCQH DHSVQVKKKP R
Zur variant 1: ASRHPEQSQH DHSVQVKKKP R
Zur variant 2: ASRHPEQSQH DHSVQVKKKP R
Zur variant 3: ASRHPEQCQH DHSVQVKKKP R

Zur variant 1 for $Zur^{Cy5\text{-}C113}$: E coli Zur (C17S, C152S, C158S)- expected mass 19205.6 Da
Zur variant 2 for $Zur^{Cy5\text{-}C113}_{D49A}$: E coli Zur (C17S, C152S, C158S, D49A)- expected mass 19161.6 Da
Zur variant 3 for $Zur^{Cy5\text{-}C158}$: E coli Zur (C17S, C113S, C152S)- expected mass 19205.6 Da

**b**

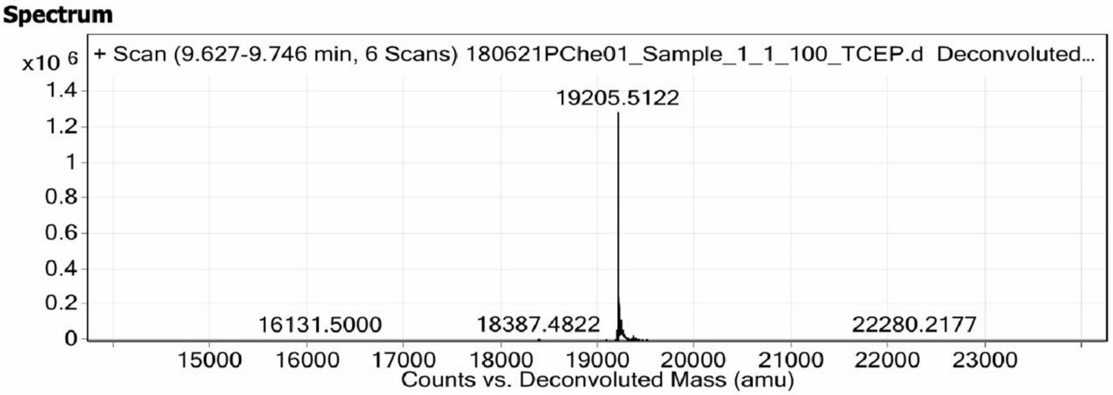


**c**

**PChe01 (100%), 19,161.6 Da**
**Zinc uptake regulation protein (Zur), E. coli, C17S, D49A, C152S, C158S [mutant of P0AC51 (ZUR_ECOLI)]**
**79 exclusive unique peptides, 159 exclusive unique spectra, 578 total spectra, 156/171 amino acids (91% coverage)**

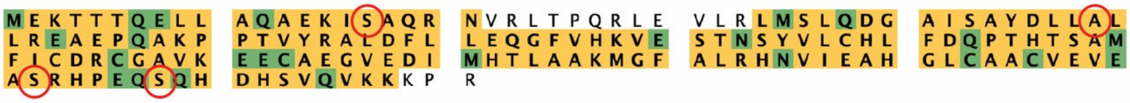


**d**

**PChe02 (100%), 19,205.6 Da**
**Zinc uptake regulation protein, E. coli, C113S, C152S [mutant of P0AC51 (ZUR_ECOLI)]**
**32 exclusive unique peptides, 81 exclusive unique spectra, 247 total spectra, 168/171 amino acids (98% coverage)**

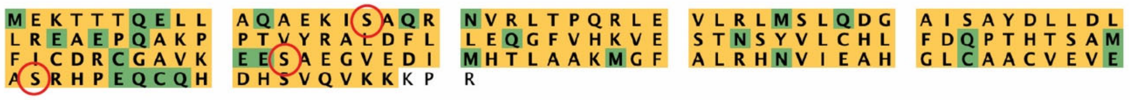

**Supplementary Fig. 18 | Identities of recombinant Zur variants are confirmed with mass spectrometry. a,** Amino-acid sequences of *E. coli* Zur and our designed variants. The specific mutant residues and expected mass for each variant are written below the sequences. **b,** Mass of Zur variant 1 is determined by ESI-TOF. The protein mass agrees with the expected value. **c-d,** Amino acid sequences of Zur variant 2 (c) and 3 (d) are determined by LC-MS/MS. The results are visualized via the Scaffold software (Proteome Software). Amino-acids matched to a MS/MS spectrum are highlighted (yellow/green). Amino-acids in green have a post-translational modification. Each variant is observed with >90% coverage and all mutation residues circled in red are confirmed.

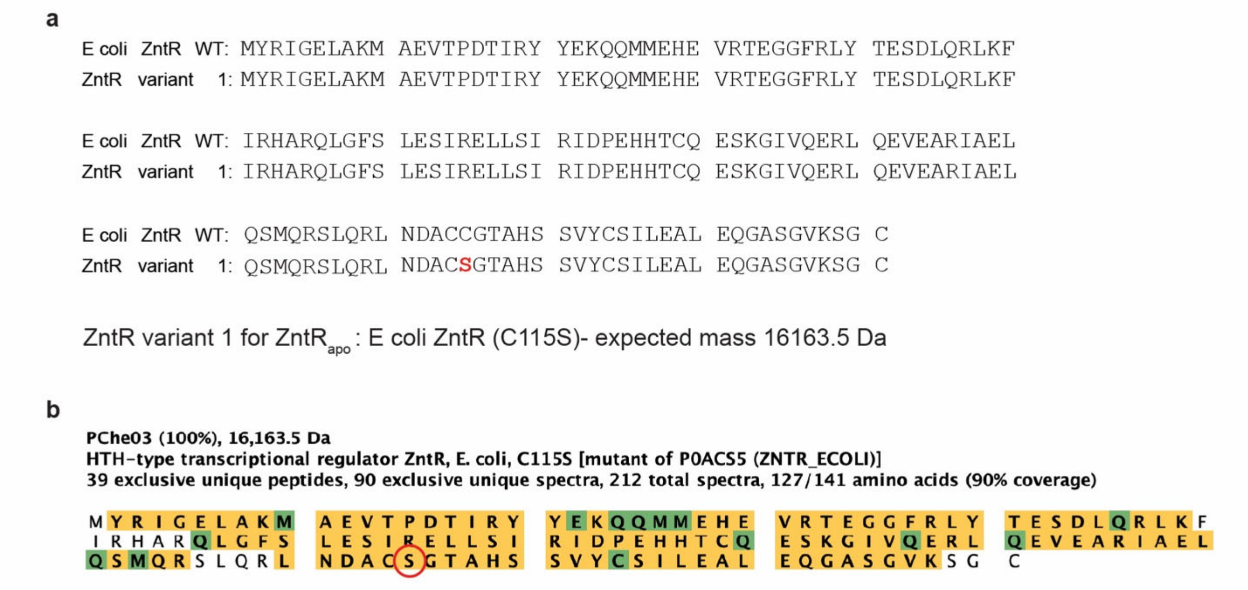


**Supplementary Fig. 19 | Identity of recombinant ZntR variant is confirmed with mass spectrometry. a,** Amino-acid sequences of *E. coli* ZntR and our designed variant. The specific mutant residue and expected mass for the variant are written below the sequences. **b,** Amino acid sequence of ZntR variant 1 is determined by LC-MS/MS. The results are visualized via the Scaffold software (Proteome Software). Amino-acids matched to a MS/MS spectrum are highlighted (yellow/green). Amino-acids in green have a post-translational modification. ZntR variant is observed with 90% coverage confirming the C115S mutation circled in red.

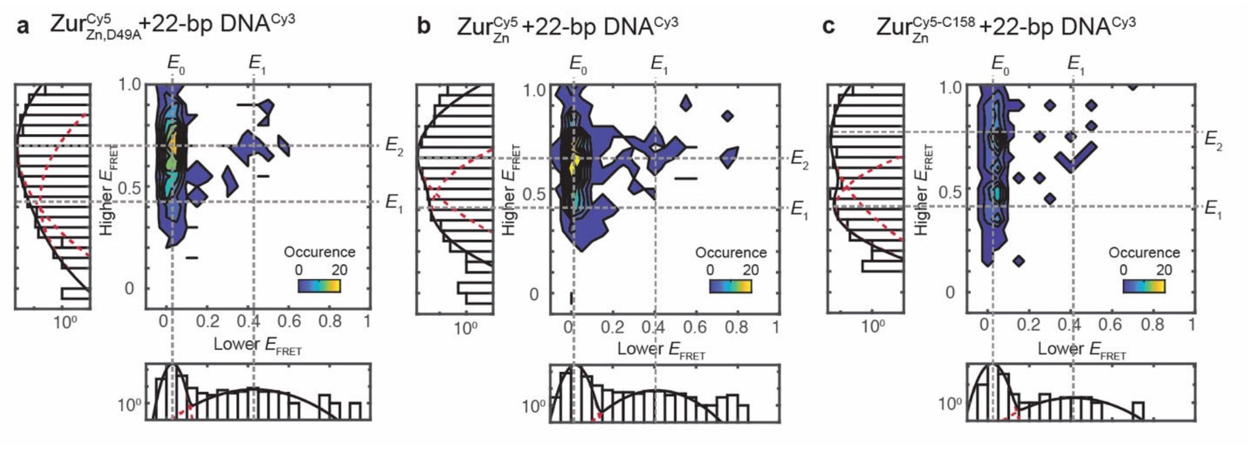


**Supplementary Fig. 20 |** Two-dimensional histogram of the lower vs. higher $E_{FRET}$ state values from single-molecule $E_{FRET}$ trajectories of an immobilized 22-bp truncated DNA$^{Cy3}$ interacting with 4 nM of **a,** Zur$^{Cy5}_{Zn,D49A}$ (Cy5 at C113), **b,** Zur$^{Cy5}_{Zn}$ (Cy5 at C113), and **c,** Zur$^{Cy5-C158}_{Zn}$. Left and bottom: corresponding one-dimensional projections. Red dashed lines: Gaussian-resolved fits; black lines: overall fits.

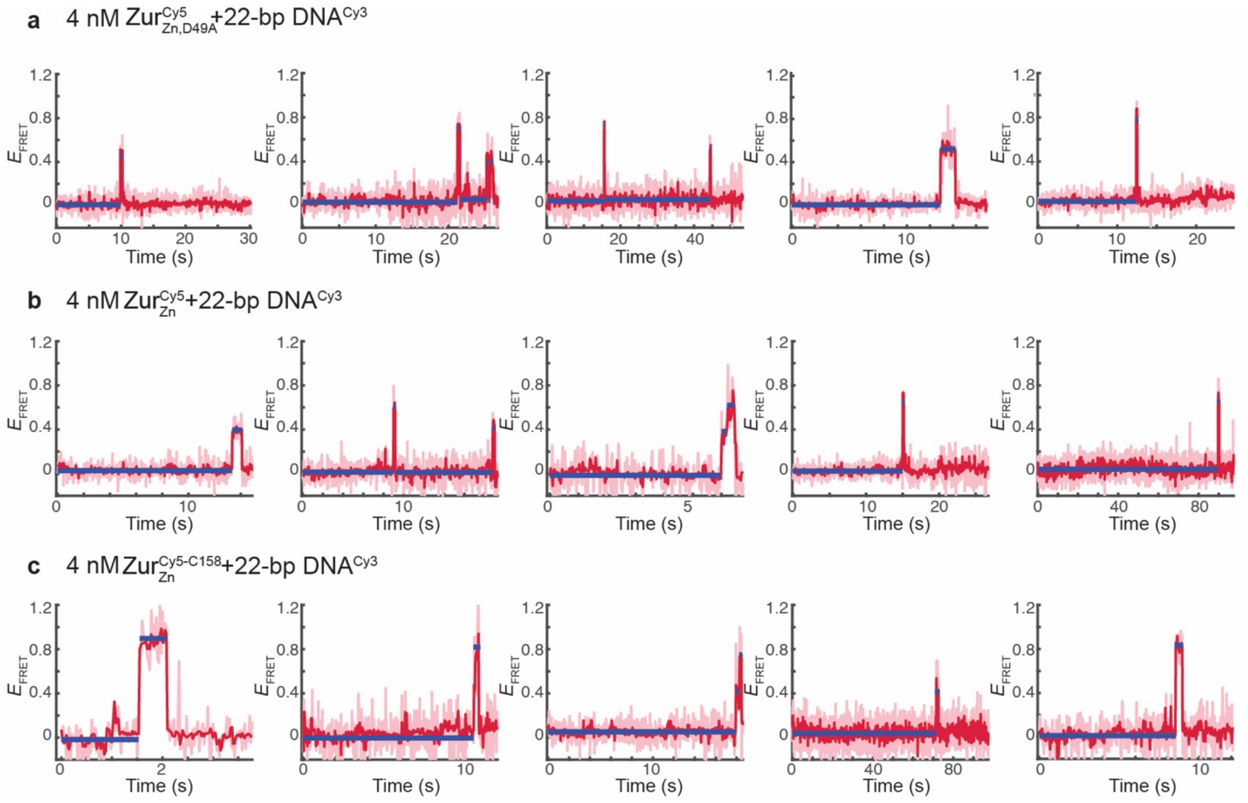


**Supplementary Fig. 21** | Examples of single-molecule $E_{FRET}$ trajectories of an immobilized 22-bp truncated DNA$^{Cy3}$ interacting with 4 nM of **a,** Zur$^{Cy5}_{Zn,D49A}$ (Cy5 at C113), **b,** Zur$^{Cy5}_{Zn}$ (Cy5 at C113), and **c,** Zur$^{Cy5-C158}_{Zn}$. Pink lines: raw data; red lines: after non-linear filtering; blue lines: mean value of each $E_{FRET}$ state.

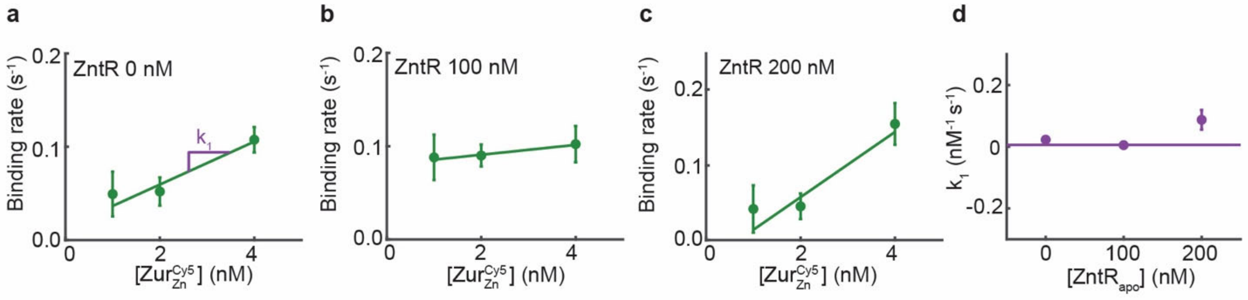


**Supplementary Fig. 22 | Binding rate constant of Zur$^{Cy5}_{Zn}$ (Cy5 at C113) on 31-bp DNA$^{Cy3}$ is independent of ZntR$_{apo}$ protein concentration.** Binding rate constant ($k_1$) is the slope of the graph of binding rate vs. [Zur$^{Cy5}_{Zn}$] in the presence of **a,** 0 nM, **b,** 100 nM, **c,** 200 nM of ZntR$_{apo}$. **d,** [ZntR$_{apo}$]-independent binding rate constant ($k_1$) of Zur$^{Cy5}_{Zn}$. Lines indicate linear fit of each graph.

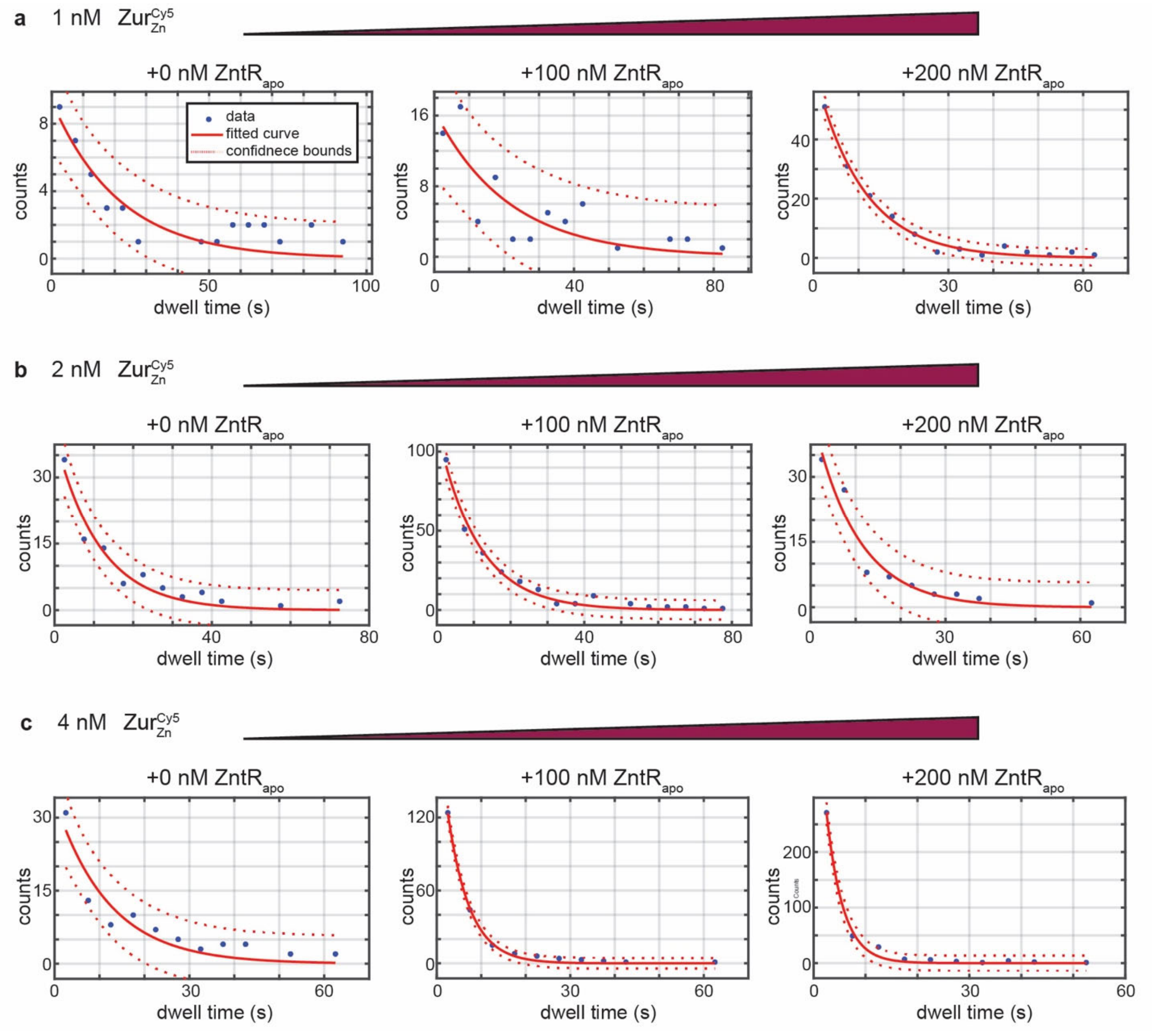


**Supplementary Fig. 23 |** The distributions of dwell time (i.e., all $\tau_{bound}$) from Zur$^{Cy5}_{Zn}$(Cy5 at C113) + 31-bp DNA$^{Cy3}$ interactions using *in vitro* smFRET measurement as in Fig. 3b at a protein concentration of **a,** 1 nM, **b,** 2 nM, **c,** 4 nM in the presence of 0-200 nM ZntR$_{apo}$. The corresponding single exponential fits ($y = A$*exp(-$k_{eff}$* $\tau$)) are shown in red solid lines. Red dashed lines are 90% confidence bounds. Rate constants are summarized in Extended Data Table 2. All bin sizes: 0.10 s.

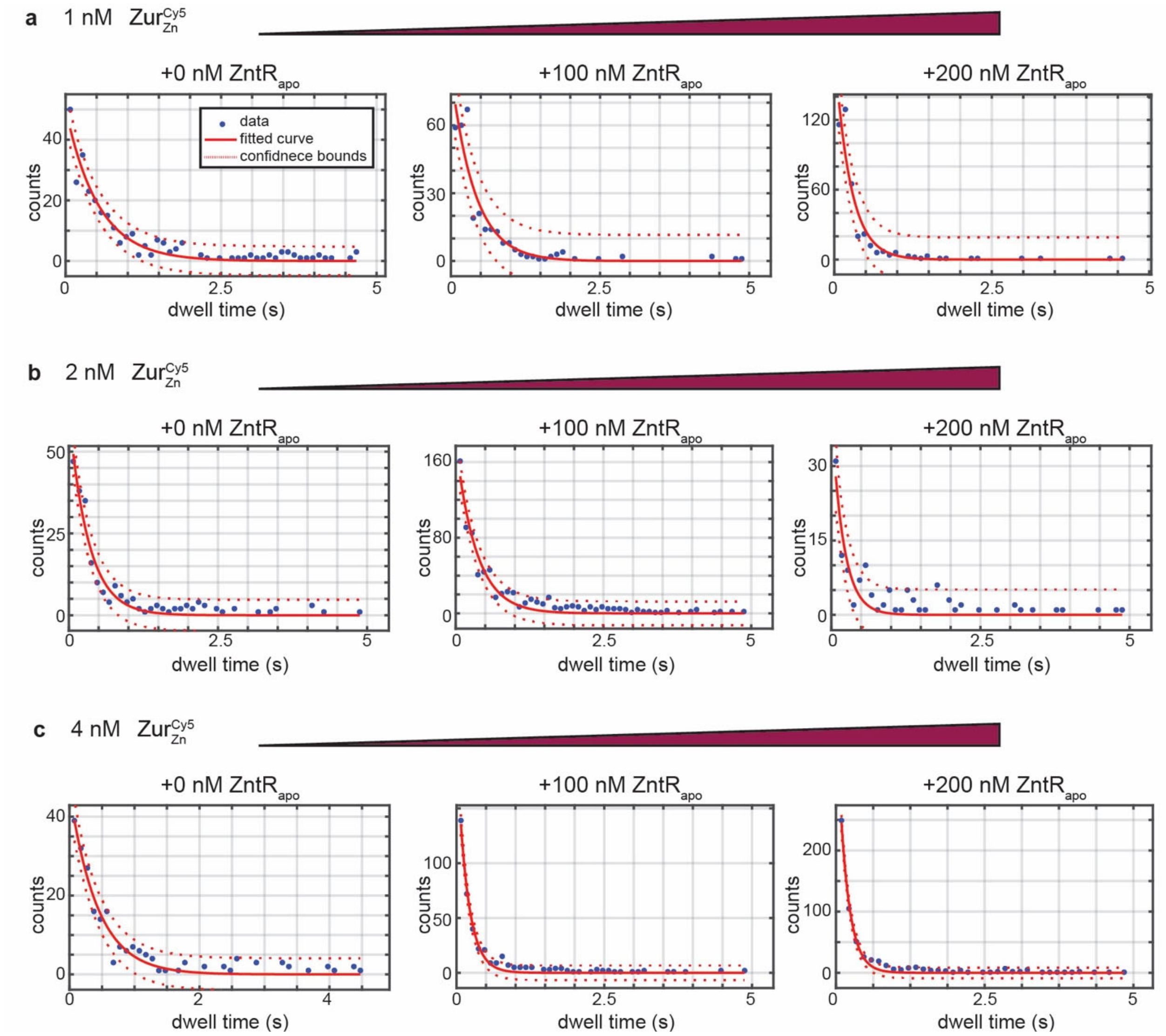


**Supplementary Fig. 24 |** The distributions of $\tau_{\text{unbound}}$ from $\text{Zur}_{\text{Zn}}^{\text{Cy5}}$(Cy5 at C113) + 31-bpDNA$^{\text{Cy3}}$ interactions using *in vitro* smFRET measurement as in Fig. 3b at a protein concentration of **a,** 1 nM, **b**, 2 nM, **c,** 4 nM in the presence of 0-200 nM ZntR$_{\text{apo}}$. The corresponding single exponential fits ($y = A$*exp($-k_{\text{eff}}$*$\tau$)) are shown in red solid lines. Red dashed lines are 90% confidence bounds. Rate constants are summarized in Extended Data Table 2. All bin sizes: 5 s.
